# FAM237A, rather than peptide PEN and proCCK56-63, is a ligand of the orphan receptor GPR83

**DOI:** 10.1101/2022.09.27.509696

**Authors:** Hao-Zheng Li, Ya-Fen Wang, Xiao-Xia Shao, Ya-Li Liu, Zeng-Guang Xu, Shi-Long Wang, Zhan-Yun Guo

**Author notes:** **Correspondence to Z.-Y. Guo**, Research Center for Translational Medicine at East Hospital, School of Life Sciences and Technology, Tongji University, 1239 Siping Road, Shanghai 200092, China. These authors contributed equally to this work (Hao-Zheng Li, Ya-Fen Wang).

## Abstract

G protein-coupled receptor 83 (GPR83) is primarily expressed in the brain and is implicated in the regulation of energy metabolism and some behaviors. Recently, the PCSK1N/proSAAS-derived peptide PEN, the procholecystokinin-derived peptide proCCK56-63, and family with sequence similarity 237 member A (FAM237A) were all reported as agonists of GPR83. However, these results have not yet been reproduced by other laboratories and thus GPR83 is still officially an orphan receptor. The PEN and proCCK56-63 share sequence similarity; however, they are completely different from FAM237A, raising doubts that all of them are ligands of GPR83. To identify its actual ligand(s), in the present study we developed a NanoLuc Binary Technology (NanoBiT)-based ligand-binding assay, fluorescent ligand-based visualization, and a NanoBiT-based β-arrestin recruitment assay for human GPR83. Using these assays, we demonstrated that mature human FAM237A could bind to GPR83 with nanomolar range affinity, which activated this receptor and induced its internalization in transfected human embryonic kidney 293T cells. However, we did not detect any interaction of PEN and proCCK56-63 with GPR83 using these assays. Thus, the results confirmed that FAM237A is an agonist of GPR83, but did not support PEN and proCCK56-63 as ligands of this receptor. Clarification of its actual endogenous agonist will pave the way for further functional studies of this brain-specific receptor. The present study also provided an efficient approach for the preparation of mature FAM237A, which would facilitate further functional studies of this difficult-to-make peptide in the future.

## Introduction

G protein-coupled receptor 83 (GPR83), also known as GPR72, glucocorticoid-induced receptor (GIR), or JP05, is an A-class G protein-coupled receptor (GPCR) primarily expressed in the brain. Previous studies suggested that GPR83 is involved in the regulation of energy metabolism and some anxiety-related behaviors [1□4]. GPR83 is also expressed in the thyroid and adrenal glands; however, its regulation of T-cell function is controversial [5□7]. GPR83 is widely present in vertebrates, and is highly conserved from fish to mammals (Fig. S1), suggesting it has important functions. The human *GPR83* gene is located on chromosome 11, and can produce two transcripts. Transcript 1 (NM_016540) encodes a longer isoform 1 (423 amino acids) that has a typical GPCR architecture with seven transmembrane domains (TMDs) and a cleavable N-terminal signal peptide (Fig. 1A), while transcript 2 (NM_001330345) encodes a shorter isoform 2 (381 amino acids) that lacks TMD3 compared with isoform 1. In previous publications and in the present study, GPR83 is refers to the longer isoform 1 unless otherwise stated. As predicted using the AlfaFold2 algorithm [8,9], mature human GPR83 (407 amino acids) has a typical seven helical-bundle structure (Fig. 1B). The highly conserved residues are mainly clustered in TMD3, TMD6, and TMD7 (Fig. 1A,B and Fig. S1). These residues seem responsible for structural integrity, ligand binding, and downstream signaling. Among them, some conserved extracellular residues, such as H331, L308, N309, N124, and Q151, might form a ligand-binding pocket (Fig. 1B).

**Fig. 1.**
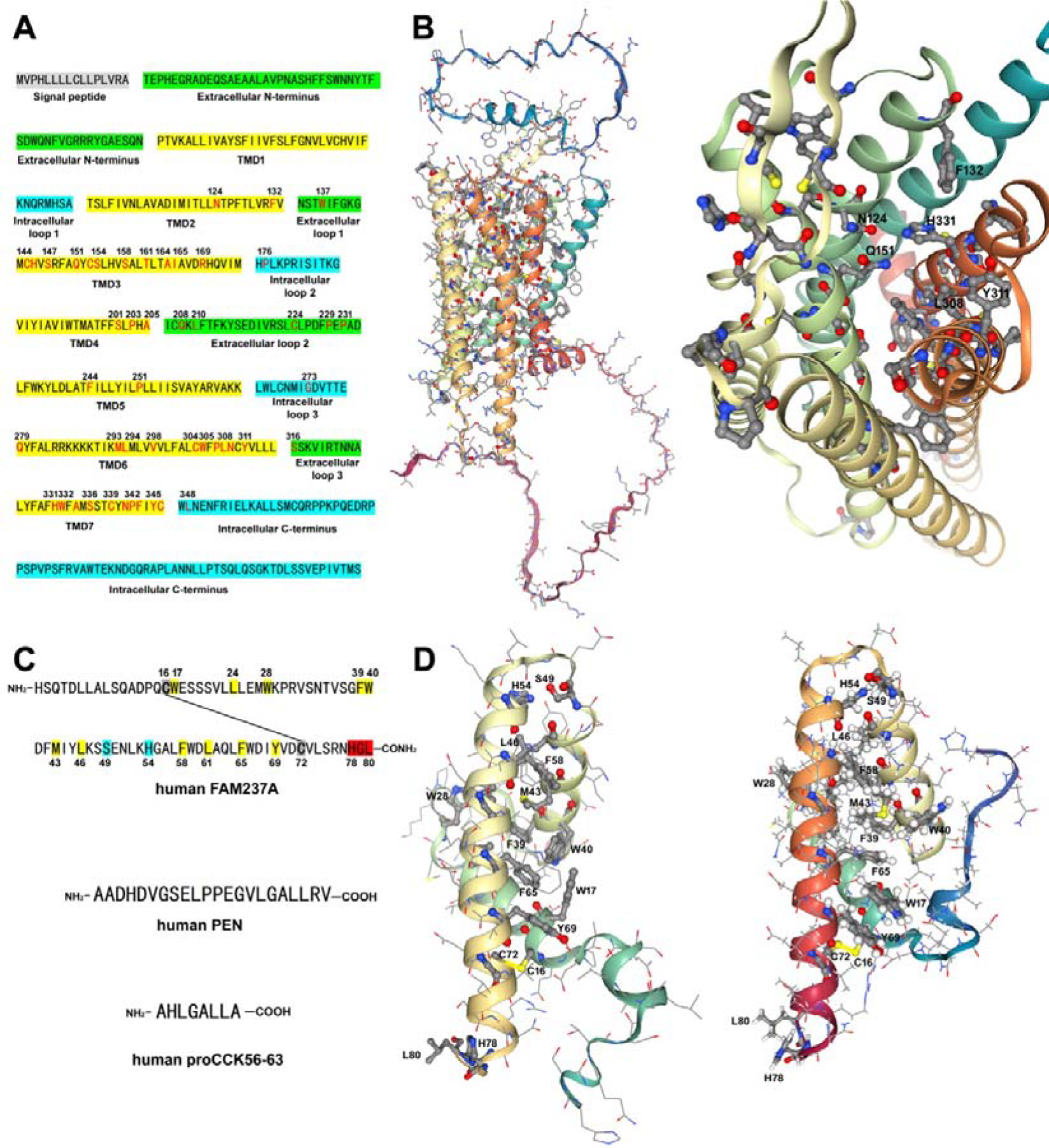
(**A**) Amino acid sequence of the full-length human GPR83. Positions of the predicted TMDs, extracellular or intracellular regions, and the signal peptide are highlighted by different colors and indicated. The highly conserved residues are shown in red. (**B**) The predicted three-dimensional structure (AF-Q9NYM4-F1-model_v2) of the mature human GPR83. Left penal, overall structure; right panel, the possible ligand-binding pocket. Side-chains of the highly conserved residues are shown as balls and sticks. (**C**) Amino acid sequence of the mature human FAM237A, PEN, and proCCK56-63. Highly conserved residues in FAM237A are highlighted by different colors, yellow for those forming the hydrophobic core, cyan for those forming a critical hydrogen bond, and red for those possibly interacting with receptor. (**D**) Three dimensional structures of the mature human FAM237A predicted by the AlphaFold algorithm (left panel, AF-A0A1B0GTK4-F1-model_v1) or by the Rosetta *Ab Initio* algorithm (right panel). The side-chains of the conserved residues responsible for structure integrity are shown as balls and sticks, the side-chains of the conserved residues possibly responsible for receptor-binding are shown as hyperballs.

In 2016, the neuropeptide PEN was reported as an agonist for GPR83 by Devi’s group [10]. Mature PEN comprises 22 amino acids with a free carboxyl terminus (Fig. 1C). It is derived from proprotein convertase subtilisin/kexin type 1 inhibitor (PCSK1N, also known as proSAAS), a protein that also releases some other possible neuropeptides, such as big SAAS, little SAAS, big LEN, and little LEN, after *in vivo* processing [11□13]. However, this finding has not yet been reproduced by other laboratories; thus, GPR83 is still officially designated as an orphan receptor. In 2022, Devi’s group reported that a procholecystokinin (proCCK)-derived peptide, proCCK56-63, can also bind to and activate GPR83 [14], because its amino acid sequence is similar to that of the C-terminus of PEN (Fig. 1C).

In 2020, another group reported that family with sequence similarity 237 member A (FAM237A) is an efficient agonist for GPR83 [15]. FAM237A is a rarely studied protein and no other related publications can be found in PubMed. However, FAM237A is widely present in all vertebrates, and is highly conserved from fish to mammals (Fig. S2), suggesting that it plays an important, but as-yet-unknown, function. FAM237A is primarily expressed in the brain (https://www.proteinatlas.org/ ENSG00000235118-FAM237A), and is synthesized *in vivo* as a precursor (Fig. S2). After a series of posttranslational processes, including removal of the N-terminal signal peptide and the C-terminal propeptide, C-terminal amidation, and the formation of an intramolecular disulfide bond, the expected mature FAM237A is released [15]. Mature human FAM237A comprises 80 amino acids, with an amidated C-terminus and an intramolecular disulfide bond (Fig. 1C). Two prediction algorithms generated quite consistent three-dimensional structures for mature human FAM237A (Fig. 1D). Three helical segments are stabilized by the disulfide bond and a hydrophobic core is formed by some highly conserved residues, including W17, L24, W28, F39, W40, M43, L46, F58, L61, and F65 (Fig. 1D). The extruded C-terminus contains three highly conserved residues, H78, G79, and L80, which might insert into the ligand binding pocket of its receptor (Fig. 1D). To date, pairing of FAM237A with GPR83 has not been confirmed by other laboratories.

FAM237A has no sequence similarity with PEN or proCCK56-63 (Fig. 1C), thus it seemed strange that all of them could bind to and activate GPR83. In the present study, we developed a novel NanoLuc® Binary Technology (NanoBiT)-based binding assay, fluorescent ligand-based visualization, and NanoBiT-based β-arrestin recruitment assay for human GPR83, and demonstrated that mature human FAM237A binds to and activates human GPR83 with high efficiency, whereas human PEN and proCCK56-63 showed no detectable binding with this receptor. Our study also demonstrated that the three C-terminal residues, H78, G79, and L80, are essential for FAM237A binding to GPR83. The present study clarified that FAM237A is an agonist for GPR83, thus paving the way for further functional studies of the FAM237A-GPR83 axis.

## Results

### Preparation of mature human FAM237A via bacterial overexpression, intein-mediated amidation, and in vitro refolding

As demonstrated in a recent study [15], the C-terminus of the mature FAM237A is amidated and this posttranslational modification is essential for its activity. A dibasic cleavage site and an extended Gly residue responsible for removal of the C-terminal propeptide and subsequent α-amidation are absolutely conserved in all known FAM237A orthologs from fish to mammals (Fig. S2), suggesting that C-terminal amidation is essential for FAM237A in all vertebrates. FAM237A also contains two absolutely conserved Cys residues that form an intramolecular disulfide bond (Fig. S2). In addition, mature FAM237A seems quite hydrophobic, with hydrophobic residues accounting for ∼40% of the protein, including 5 Trp residues and 13 Leu residues. To prepare mature FAM237A with the correct posttranslational modifications, in the present study, we employed an intein-mediated amidation approach that has been used to prepare some other C-terminally amidated peptides in previous studies [16,17].

An intein molecule was genetically fused to the C-terminus of human FAM237A (Fig. S3), and the fusion protein was efficiently overexpressed in *E. coli* and formed inclusion bodies, as analyzed by sodium dodecyl sulfate-polyacrylamide gel electrophoresis (SDS-PAGE) shown in Fig. 2A. After solubilization via *S*-sulfonation and purification using a Ni^2+^ column, ∼50 mg of fusion protein could be typically obtained from one liter of culture broth. After purification, the fused intein needs to be renatured to restore its self-cleavage activity. However, the reversibly modified *S*-sulfonate moieties have to be removed from the fusion protein under denatured condition before the renaturation step to release the active site cysteine of the intein. Thus, we first treated the eluent from the Ni^2+^ column with a high concentration of reductant tris(2-carboxyethyl)phosphine (TCEP) and sodium 2-mercptoethanesulfonate (MES), and then diluted the eluent into L-arginine solution for renaturation and self-cleavage. During self-cleavage, MES is expected to form a thioester bond with the C-terminal carboxyl moiety of the released FAM237A peptide. As analyzed by SDS-PAGE (Fig. 2A), ∼70% of the fusion protein could self-cleave after overnight incubation. A weak band (indicated using an asterisk) at the bottom of the gel, presumably the released FAM237A peptide, could be observed after self-cleavage of the fusion protein (Fig. 2A).

**Fig. 2.**
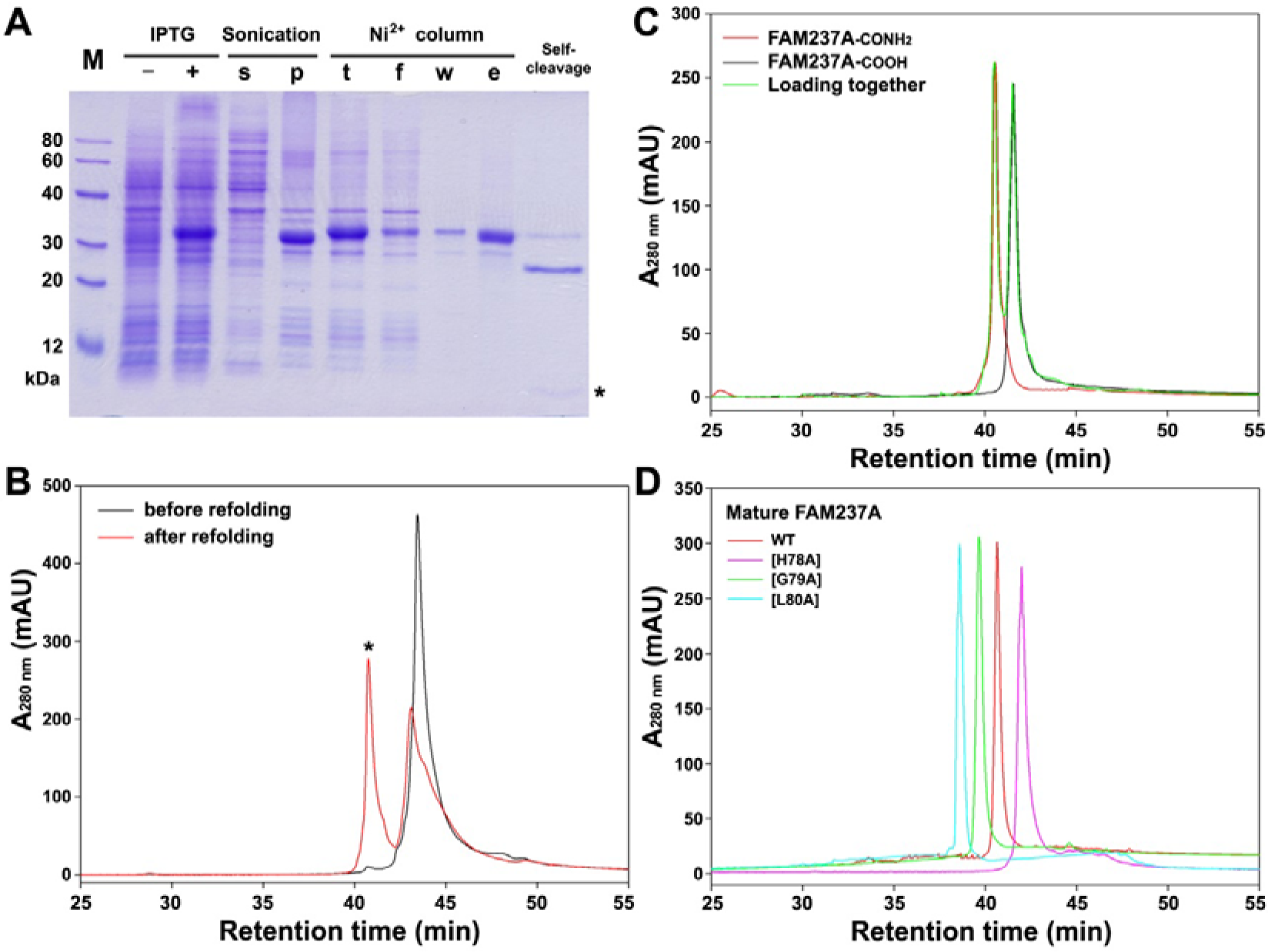
Preparation of mature human FAM237A. (**A**) SDS-PAGE analysis of the FAM237A-intein-6×His fusion protein. After overexpression in *E. coli*, samples at different stages were loaded onto a 15% SDS-gel. After electrophoresis, the gel was stained by Coomassie brilliant blue R250. The band of *S*-sulfonated FAM237A is indicated by an asterisk. The sample loading amount was from 50 µl culture broth. Lane (–), before IPTG induction; lane (+), after IPTG induction; lane (s), supernatant after sonication; lane (p), pellet after sonication; lane (t), the *S*-sulfonated fraction loading to the Ni^2+^ column; lane (f), flow-through fraction; lane (w), washed by 30 mM imidazole; lane (e), eluent by 250 mM imidazole. (**B**) HPLC analysis of the *S*-sulfonated FAM237A before and after in vitro refolding. ∼100 µg of sample was loaded onto a C_4_ reverse-phase column and eluted by an acidic acetonitrile gradient. The peak of the folded FAM237A is indicated by an asterisk. (**C**) HPLC analysis of the refolded FAM237A with or without C-terminal amidation. The FAM237A samples with or without amidation were loaded onto a C_18_ reverse-phase column either separately or together, and then eluted from the column by an alkaline acetonitrile gradient. (**D**) HPLC analysis of mature FAM237A mutants. The wild-type FAM237A and its mutants were loaded onto a C_18_ reverse-phase column separately, and then eluted from the column by an alkaline acetonitrile gradient.

Thereafter, the C-terminal thioester of the released FAM237A peptide was ammoniated via reaction with NH_4_HCO_3_ under high pH conditions, forming the C-terminally α-amidated product. During ammoniation, most of the FAM238A peptide and some uncleaved fusion protein formed precipitate that was collected and solubilized again via *S*-sulfonation. To prevent FAM237A sticking to the Ni^2+^ column, 0.5% Tween-20 was added to the sulfonation solution. Thereafter, the uncleaved fusion protein was absorbed using a Ni^2+^ column, and the flow-through fraction containing the *S*-sulfonated FAM237A was applied to high performance liquid chromatography (HPLC). The *S*-sulfonated FAM237A could be eluted from a C_4_ reverse-phase column using an acidic acetonitrile gradient; however, it could not be eluted from a C_18_ reverse-phase column using the acidic acetonitrile gradient because of its high hydrophobicity. Finally, the *S*-sulfonated FAM237A was subjected to *in vitro* refolding to form its intramolecular disulfide bond. As analyzed by HPLC, a new peak (indicated using an asterisk) with a shorter retention time, presumably the expected refolded product, was eluted from the C_4_ reverse-phase column using the acidic acetonitrile gradient (Fig. 2B). As estimated from the peak area, the refolding yield of FAM237A was ∼50%.

To confirm the FAM237A prepared via the intein-mediated approach was actually amidated, we also prepared non-amidated FAM237A by conducting self-cleavage of the fusion protein in the presence of dithiothreitol only. As analyzed by HPLC, the amidated FAM237A (FAM237A-CONH_2_) and the non-amidated peptide (FAM237A-COOH) had different retention times when eluted from a C_18_ reverse-phase column using an alkaline acetonitrile gradient (Fig. 2C), confirming that the mature FAM237A was correctly amidated via the intein-based approach.

In the predicted structures (Fig. 1D), the C-terminal fragment of FAM237A extends outside and might insert into the ligand-binding pocket of its receptor. To test the importance of the C-terminal fragment, three highly conserved C-terminal residues, H78, G79, and L80, were replaced by an alanine residue, respectively. The resultant FAM237A mutants were prepared using the intein-mediated approach. As analyzed by HPLC, these mutants all displayed a single symmetric peak with different retention times when eluted from the C_18_ reverse-phase column using the alkaline acetonitrile gradient (Fig. 2D).

### Mature FAM237A binds to GPR83 with high affinity and high specificity

To test whether mature FAM237A is a ligand of GPR83, we first employed the NanoBiT-based ligand□receptor binding assay that was recently developed by our laboratory and has been validated using certain GPCRs, such as relaxin family peptide receptor (RXFP) 3 and 4, and growth hormone secretagogue receptor type 1a (GHSR1a) [18–20]. To set up this binding assay, an inactive secretory large NanoLuc fragment for NanoBiT (sLgBiT) was genetically fused to the extracellular N-terminus of mature human GPR83 (Fig. S4), and a low-affinity small complementation tag for NanoBiT (SmBiT) was genetically fused to the N-terminus of mature FAM237A (Fig. S3). If the SmBiT-FAM237A tracer could bind to sLgBiT-GPR83, a proximity effect would induce complementation of the receptor-fused sLgBiT with the ligand-fused SmBiT and thus restore luciferase activity (Fig. 3A). In theory, this NanoBiT-based binding assay can detect specific binding of the sLgBiT-fused receptor and the SmBiT-fused ligand, because the endogenously expressed receptors in the host cells have no sLgBiT fusion and thus would not cause interference.

**Fig. 3.**
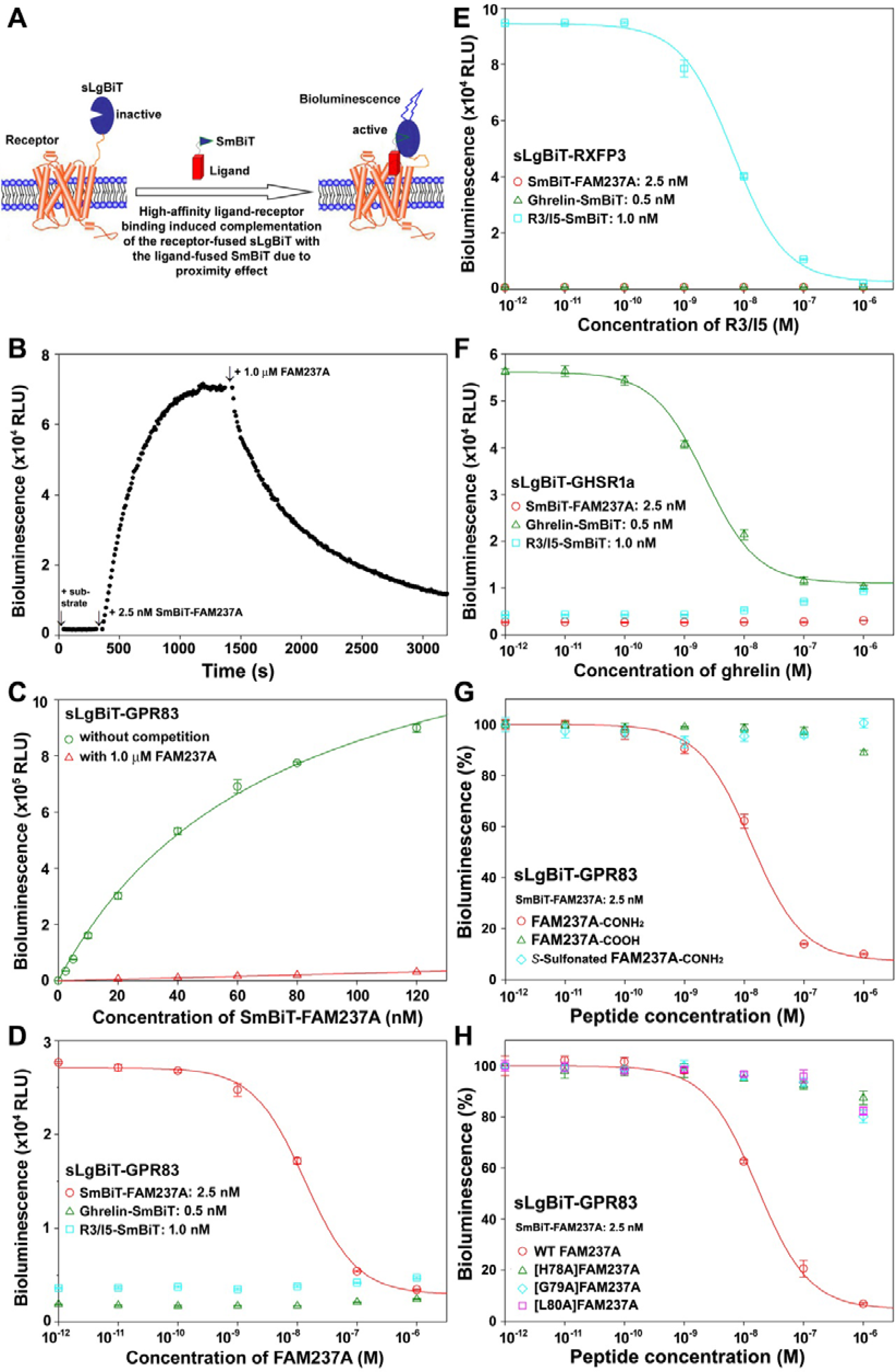
NanoBiT-based binding of FAM237A with GPR83. (**A**) Schematic presentation of the NanoBiT-based ligand–receptor binding assay. (**B**) Association and dissociation of SmBiT-FAM237A with sLgBiT-GPR83 assayed on living HEK293T cells. At the indicated time, NanoLuc substrate, SmBiT-FAM237A, and mature FAM237A were sequentially added to living HE293T cells overexpressing sLgBiT-GPR83, and bioluminescence was monitored continuously on a plate reader with an interval of 10 s. (**C**) Saturation binding of SmBiT-FAM237A with sLgBiT-GPR83 assayed on living HEK293T cells. (**D**) Binding of various SmBiT-based tracers to sLgBiT-GPR83 and competition by mature FAM237A. (**E**) Binding of various SmBiT-based tracers to sLgBiT-GHSR1a and competition by ghrelin. (**F**) Binding of various SmBiT-based tracers to sLgBiT-RXFP3 and competition by mature R3/I5. (**G**,**H**) Binding potency of some FAM237A analogs with GPR83 measured by the NanoBiT-based competition binding assay. In panel C–H, the measured binding data are expressed as mean ± SD (*n* = 3), and fitted to one-site binding model using SigmaPlot 10.0 software.

After the NanoLuc substrate was added to living HEK293T cells transiently overexpressing sLgBiT-GPR83, the background bioluminescence was quite low (Fig. 3B), confirming that the receptor-fused sLgBiT itself was enzymatically inactive. After the addition of 2.5 nM SmBiT-FAM237A, the measured bioluminescence increased rapidly, reaching a plateau within 15 min, with an association half-life of ∼3 min. The rapid increase in bioluminescence was presumably caused by the binding of SmBiT-FAM237A with sLgBiT-GPR83. After the addition of 1.0 µM of mature FAM27A as a competitor, the measured bioluminescence decreased rapidly, with a dissociation half-life of ∼7.5 min. This rapid decrease in bioluminescence was presumably caused by competitive displacement of SmBiT-FAM237A from sLgBiT-GPR83 by mature FAM237A. Thus, it seemed that SmBiT-FAM237A can reversibly bind to sLgBiT-GPR83 and their association and dissociation can be conveniently monitored in a real-time manner according to the measured bioluminescence.

Thereafter, we measured the binding affinity of SmBiT-FAM237A with sLgBiT-GPR83 via a saturation binding assay (Fig. 3C). As the tracer concentration increased, the measured bioluminescence increased in a typical hyperbolic manner, with a calculated dissociation constant (K_d_) of ∼70 nM (Fig. 3C). The presence of 1.0 μM mature FAM237A drastically lowered the measured bioluminescence (Fig. 3C), confirming that the tracer-induced bioluminescence was originated from binding of SmBiT-FAM237A with sLgBiT-GPR383. Thus, it seemed that SmBiT-FAM237A can bind to sLgBiT-GPR83 with high affinity.

To confirm the specific binding of FAM237A with GPR83, we employed other SmBiT-based tracers and the corresponding sLgBiT-fused receptors (ghrelin-SmBiT versus sLgBiT-GHSR1a; R3/I5-SmBiT versus sLgBiT-RXFP3) generated in our previous studies [18,19]. Towards sLgBiT-GPR83, only the addition of SmBiT-FAM237A could induce high bioluminescence, and the addition of mature FAM237A could decrease it in a sigmoidal manner (Fig. 3D), with a calculated IC_50_ value of 13.5 ± 0.9 nM (*n* = 3). In contrast, the addition of FAM237A had no effect on the background bioluminescence in R3/I5-SmBiT and ghrelin-SmBiT groups (Fig. 3D). Towards sLgBiT-GHSR1a and sLgBiT-RXFP3, the addition of SmBiT-FAM237A could not induce high bioluminescence; however, the addition of their cognate tracer could (Fig. 3E,F). Moreover, addition of their cognate ligands could only decrease the bioluminescence induced by their cognate tracers (Fig. 3D,E). Thus, it seemed that FAM237A binds to GPR83 with high specificity.

Mature FAM237A contains one intramolecular disulfide bond and an amidated C-terminus. To test the importance of these posttranslational modifications, we measured the receptor-binding potency of the non-amidated FAM237A (FAM237A-COOH) and the *S*-sulfonated FAM237A (with an amidated C-terminus, but without a disulfide bond). In the NanoBiT-based competition binding assay (Fig. 3G), mature FAM237A with the correct posttranslational modifications could efficiently displace the SmBiT-FAM237A tracer from sLgBiT-GPR83, with a calculated IC_50_ of 12.9 ± 0.9 nM (*n* = 3). However, the non-amidated FAM237A and the *S*-sulfonated FAM237A had almost no effects (Fig. 3G), confirming that the C-terminal amidation and the disulfide bond are essential for FAM237A binding to GPR83.

According to the predicted structures (Fig. 1D), the extended C-terminus might insert into the ligand-binding pocket of GPR83. The three C-terminal residues, H78, G79, and L80, are highly conserved in evolution (Fig. S1). After they were replaced by an alanine residue, the resultant FAM237A mutants almost lost their binding to GPR83 (Fig. 3H), implying that these C-terminal residues are critical for FAM237A function. As a positive control, the wild-type FAM237A could efficiently displace the tracer, with a calculated IC_50_ value of 16.9 ± 1.3 nM (*n* = 3). The large H78 and L80 residues might directly interact with counterpart receptor residues, while the smallest G79 might provide sufficient flexibility for L80 and H78 to efficiently interact with the receptor.

### PEN and proCCK56-63 have no detectable binding with GPR83

We first tested whether PEN and proCCK56-63 could bind to GPR83 via the NanoBiT-based competition binding assay using SmBiT-FAM237A as a tracer (Fig. 4A). As a positive control, mature FAM237A could displace SmBiT-FAM237A from sLgBiT-GPR83 in a typical sigmoidal manner (Fig. 4A). However, synthetic PEN and proCCK56-63 could not displace the tracer at all (Fig. 4A), suggesting they either had no binding with this receptor, or they bound to a distinct site without any interference to FAM237A binding. Considering that FAM237A, PEN, and proCCK56-63 are quite large peptides, it is difficult to imagine that they could bind simultaneously to GPR83 without any interference. As shown in Fig. 1B, it seemed that the ligand binding pocket of GPR83 is not large enough to simultaneously accommodate FAM237A and another peptide without any interference. Thus, the interpretation of our data was that PEN and proCCK56-63 cannot bind to GPR83.

**Fig. 4.**
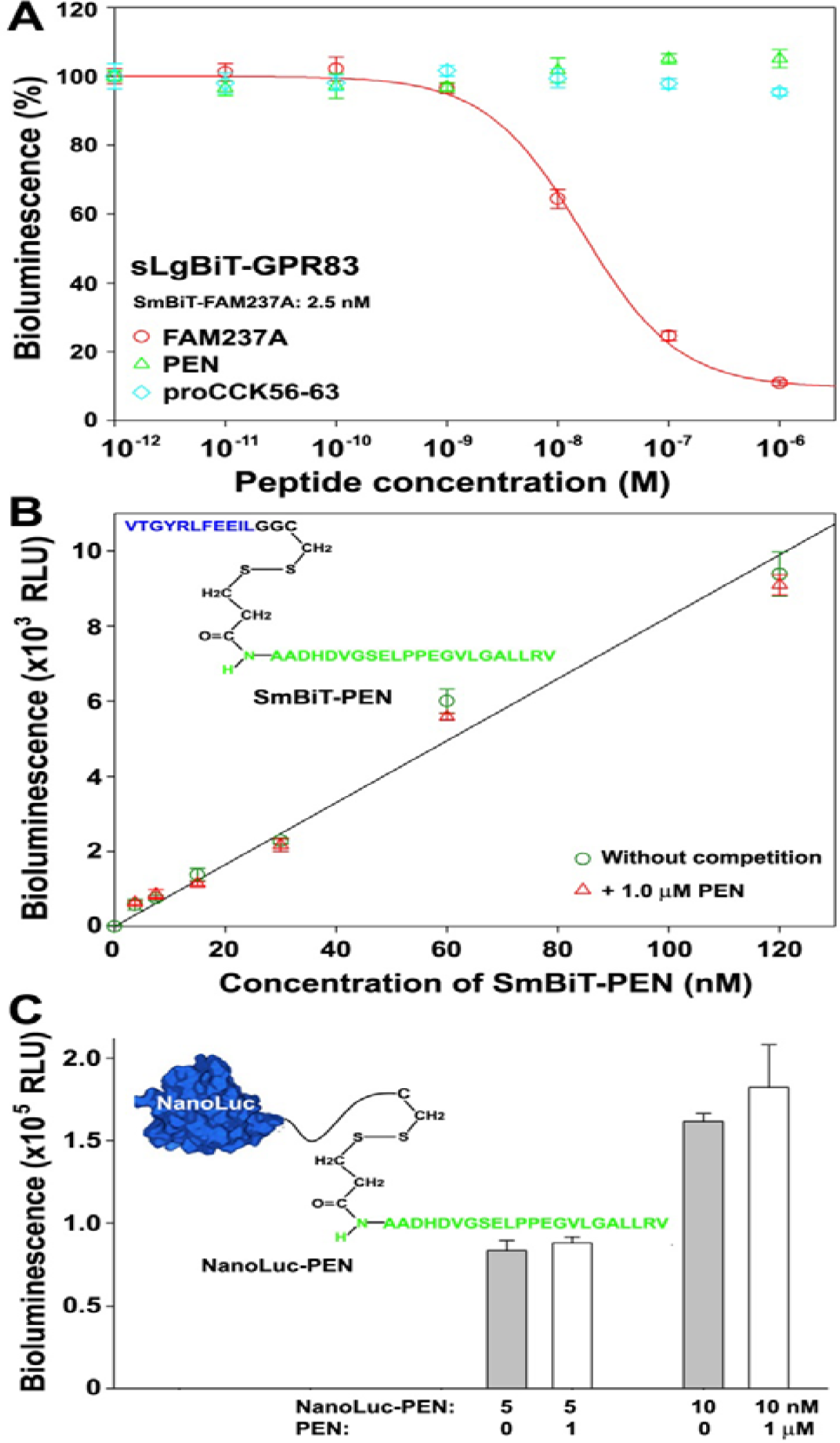
Detection of binding of PEN and proCCK56-63 with GPR83. (**A**) Binding detection via NanoBiT-based competition binding assay using SmBiT-FAM237A as a tracer. (**B**) Binding detection using SmBiT-PEN as a tracer assayed on living HEK293T cells overexpressing sLgBiT-GPR83. (**C**) Binding detection using NanoLuc-PEN as a tracer assayed on living HEK293T cells overexpressing untagged GPR83. The measured binding data are expressed as mean ± SD (*n* = 3).

We prepared a SmBiT-PEN tracer via chemical conjugation of a C-terminally Cys-tagged SmBiT to the N-terminus of synthetic PEN (Fig. 4B, inner panel). When SmBiT-PEN was added to living HEK293T cells overexpressing sLgBiT-GPR83, the measured bioluminescence was very low, and competition with 1.0 µM PEN had no effect, suggesting that the SmBiT-PEN tracer either could not bind to sLgBiT-GPR83 or its binding could not induce complementation of its SmBiT with the receptor-fused sLgBiT because of unexpected steric hindrance.

We also prepared a NanoLuc-based tracer, NanoLuc-PEN, by chemical conjugation of a C-terminally Cys-tagged NanoLuc to the N-terminus of synthetic PEN (Fig. 4C, inner panel). The NanoLuc-based binding tracers have been widely validated using different receptors in our previous studies [21–28], such as RXFP1–4, GHSR1a, leukemia inhibitory factor receptor, erythropoietin receptor, and fibroblast growth factor receptor. Using NanoLuc-PEN as a tracer, we conducted washing-based binding assays using living HEK293T cells overexpressing untagged human GPR83 (Fig. 4C). Unfortunately, the measured total binding data (without competition) were identical to the corresponding non-specific binding data (competed by 1.0 µM PEN), suggesting that the NanoLuc-PEN tracer had no detectable binding to untagged GPR83.

### Visualization of FAM237A binding to GPR83 and the induction of internalization

The above binding assays confirmed that FAM237A can bind to GPR83 with nanomolar range affinity; however, the binding of PEN and proCCK56-63 with this receptor could not be detected. To confirm these results, we prepared red fluorescent dye-labeled FAM237A and PEN, and attempted to visualize whether these fluorescent peptides could bind to GPR83 using a confocal microscope. For this purpose, we generated two Tet-On inducible expression constructs that either express a C-terminally enhanced green fluorescent protein (EGFP)-fused GPR83 or coexpress an untagged GPR83 and an EGFP mediated via an internal ribosome entry site (IRES).

After HEK293T cells were treated with 10 ng/ml doxycycline (Dox) to induce expression of GPR83-EGFP fusion protein, green fluorescence (originating from the receptor-fused EGFP) could be observed in the plasma membrane and some intracellular compartments (primarily the Golgi apparatus and the endoplasmic reticulum) of some cells (presumably transfected cells) under a confocal microscope (Fig. 5A). Thus, it seemed that GPR83-EGFP can be trafficked to the plasma membrane in the transfected cells, albeit with low efficiency.

**Fig. 5.**
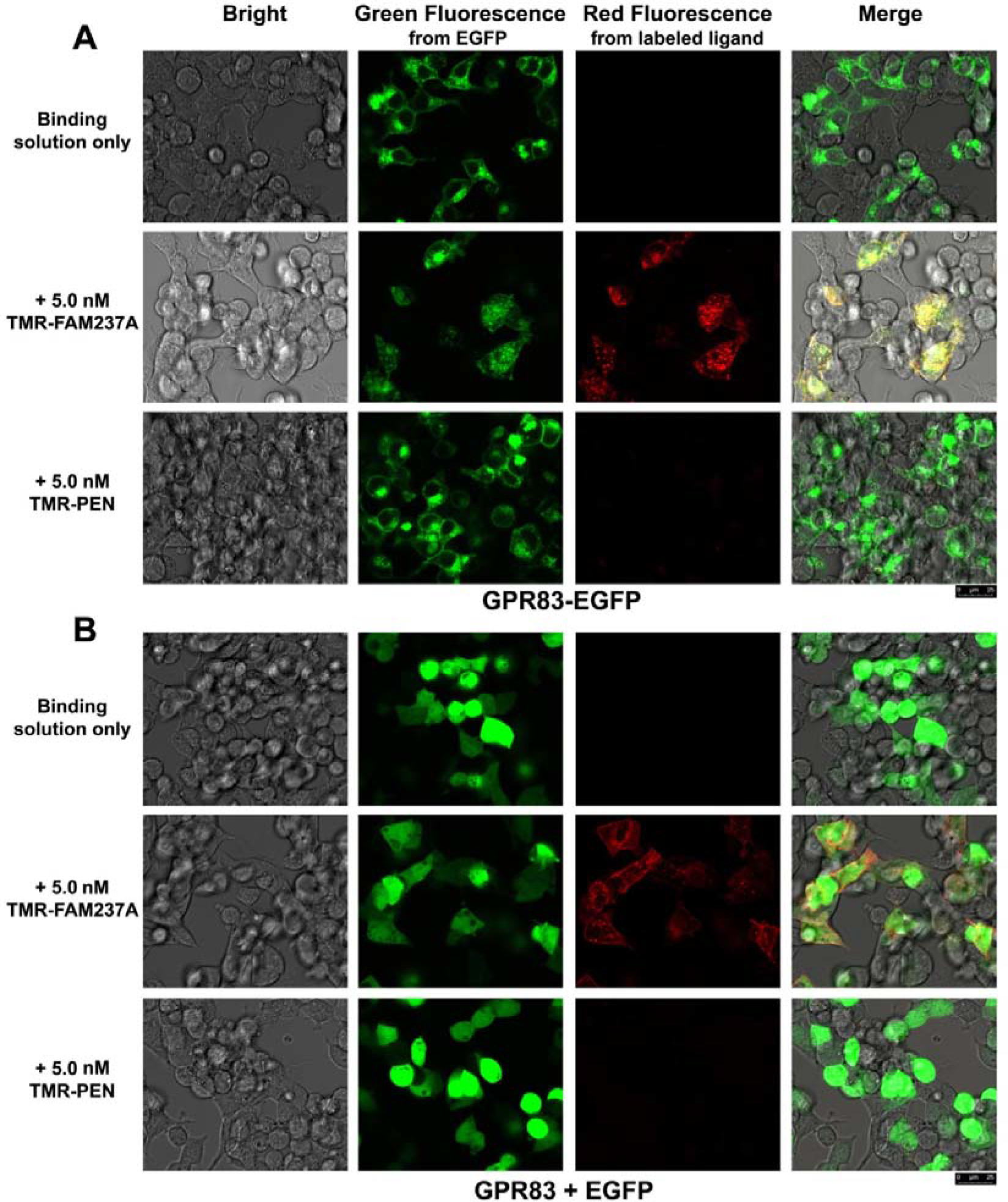
Fluorescent microscopic observation. (**A**) Fluorescent microscopic observation of living HEK293T cells expressing GPR83-EGFP fusion protein. (**B**) Fluorescent microscopic observation of living HEK293T cells coexpressing untagged GPR83 and EGFP. The living HEK293T cells were incubated either with binding solution only, or with 5 nM TMR-FAM237A, or with 5 nM TMR-PEN at 37°C for 2 h, and then observed under a confocal fluorescent microscope.

After the Dox-induced HEK293T cells were incubated with 5 nM tetramethylrhodamine (TMR)-labeled FAM237A at 37 °C for 2 h, both green fluorescence (from GPR83-fused EGFP) and red fluorescence (from TMR-FAM237A) could be observed in the plasma membrane and some intracellular dots of the transfected cells (Fig. 5A), suggesting the fluorescent dye-labeled FAM237A can bind to GPR83-EGFP and induce its internalization. For those untransfected cells (without green fluorescence), no red fluorescence could be observed (Fig. 5A), suggesting that TMR-FAM237A specifically binds to GPR83-EGFP. However, after the induced cells were incubated with 5 nM TMR-PEN, no red fluorescence could be observed in any cells (Fig. 5A), suggesting that the fluorescent dye-labeled PEN has no binding with GPR83-EGFP.

After HEK293T cells were treated with 10 ng/ml Dox to induce coexpression of the untagged GPR83 and EGFP, green fluorescence (originating from the coexpressed EGFP) was evenly distributed in the transfected cells (Fig. 5B), because EGFP is expressed as a cytosolic protein. After these cells were incubated with 5 nM TMR-FAM237A at 37 °C for 2 h, red fluorescence (from TMR-FAM237A) could be observed in the plasma membrane and some intracellular dots of the transfected cells (Fig. 5B). For those untransfected cells (without green fluorescence), no red fluorescence could be observed (Fig. 5B). Thus, it seemed that the fluorescent dye-labeled FAM237A can specifically bind to the untagged GPR83 and be internalized into cells together with the receptor. In contrast, no red fluorescence could be observed in any cells after incubation with 5 nM TMR-PEN (Fig. 5B), suggesting that the fluorescent PEN has no detectable binding with untagged GPR83.

### FAM237A activates GPR83 efficiently, but PEN and proCCK56-63 have no effects

To measure activation of GPR83, we employed the newly developed NanoBiT-based β-arrestin recruitment assay, which has been validated using other GPCRs in recent studies [29–31]. For this purpose, an inactive LgBiT protein was genetically fused to the C-terminus of the full-length human GPR83, and a low-affinity SmBiT tag was genetically fused to the N-terminus of human β-arrestin 2 (SmBiT-ARRB2; Fig. S5). Once GPR83-LgBiT was activated by an agonist, SmBiT-ARRB2 would bind to the activated receptor and thus the proximity effect would induce complementation of the β-arrestin-fused SmBiT with the GPR83-fused LgBiT and restore NanoLuc activity (Fig. 6A). Overexpression of GPR83 is harmful to host cells; therefore, we employed a Tet-On inducible expression vector, pTRE3G-BI, which can coexpress two proteins in a controllable manner via an inducible bi-directional promoter. In theory, the NanoBiT-based β-arrestin recruitment assay can specifically measure GPR83 activation, since no endogenously expressed receptors could cause interference because they do not carry LgBiT.

**Fig. 6.**
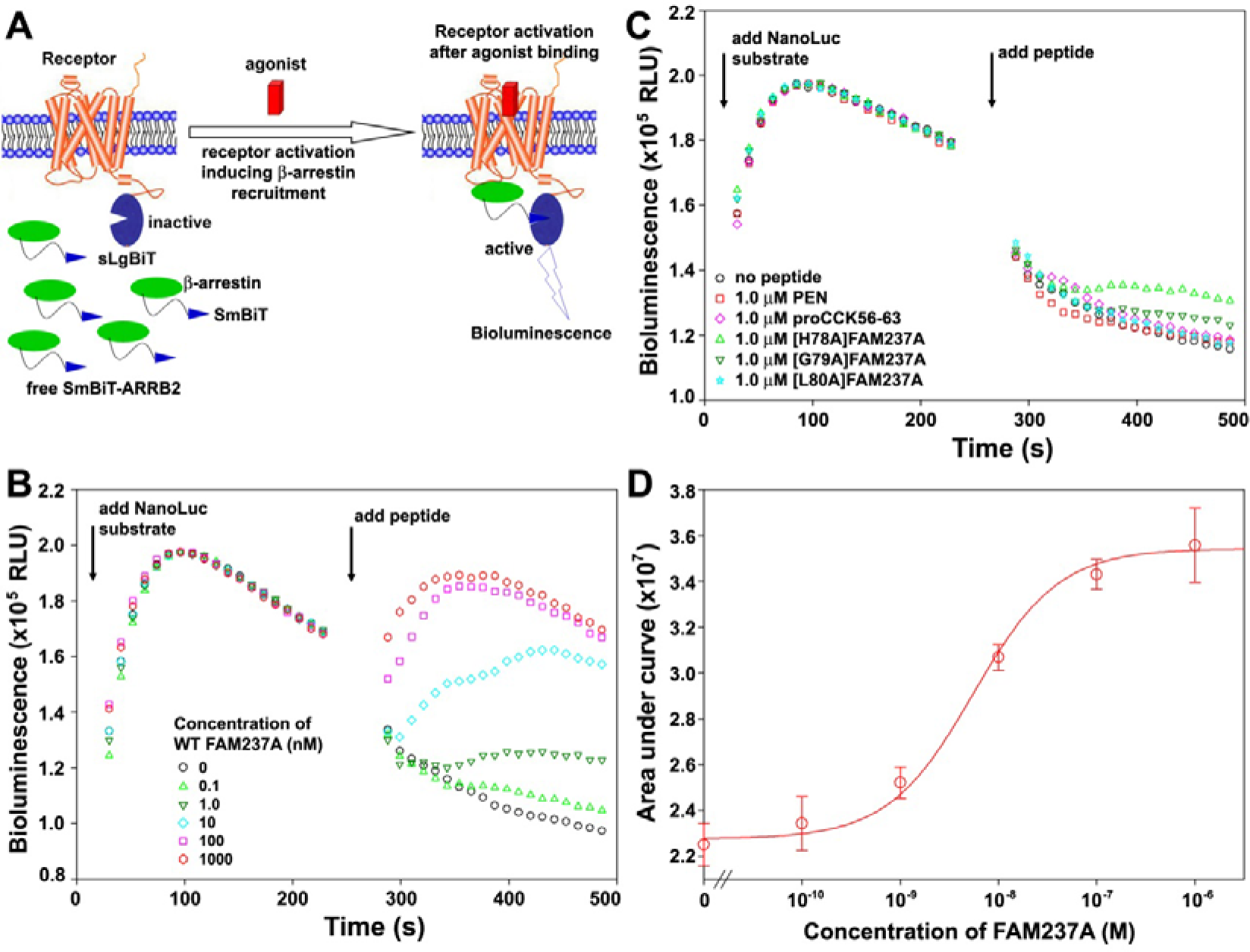
NanoBiT-based β-arrestin recruitment assay for GPR83. (**A**) Schematic presentation of the NanoBiT-based β-arrestin recruitment assay. (**B**) A typical bioluminescence change after sequential addition of NanoLuc substrate and different concentrations of mature FAM237A. (**C**) A typical bioluminescence change after sequential addition of NanoLuc substrate and 1.0 µM of different peptides. (**D**) The dose curve of FAM237A activating GPR83 assayed by the NanoBiT-based β-arrestin recruitment. The calculated AUC data are expressed as mean ± SD (*n* = 3), and plotted with FAM237A concentrations using SigmaPlot 10.0 software.

After transient transfection, expression of GPR83-LgBiT and SmBiT-ARRB2 in HEK293T cells was induced using 2.5 ng/ml Dox. After an activation solution containing the NanoLuc substrate was added to these cells, bioluminescence increased rapidly to a peak and then decreased slowly (Fig. 6B), suggesting that both GPR83-LgBiT and SmBiT-ARRB2 were expressed after induction. After mature FAM237A was added, the bioluminescence increased rapidly in a dose-dependent manner (Fig. 6B), implying that FAM237A could activate GPR83-LgBiT and induce recruitment of SmBiT-ARRB2 to the activated receptor. When the values of the area under curve (AUC) calculated from Fig. 6B were plotted with the concentrations of FAM237A, a typical sigmoidal curve was obtained (Fig. 6D), with a calculated EC_50_ value of ∼5.5 nM. Thus, mature FAM237A can efficiently activate GPR83, as assayed by β-arrestin recruitment.

However, no activation effect could be observed when up to 1.0 µM of peptide PEN or proCCK56-63 were tested (Fig. 6C), suggesting that neither peptide can activate GPR83. When the three C-terminal FAM237A mutants were tested, only [H78A]FAM237A had a weak effect (Fig. 7C), suggesting that the three C-terminal residues are important for FAM237A activation of GPR83, consistent with their low receptor-binding potency.

## Discussion

In the present study, we demonstrated that mature FAM237A is an efficient agonist of GPR83 using a NanoBiT-based ligand–receptor binding assay, fluorescent ligand-based visualization, and a NanoBiT-based β-arrestin recruitment assay, thus confirming the recent breakthrough discovery by Sallee et al [15]. We deduced that FAM237A is an endogenous agonist of GPR83 based on following considerations. (1) FAM237A binds to GPR83 with nanomolar range affinity and activates it with nanomolar efficiency. (2) Both FAM237A and GPR83 are primarily expressed in the brain. (3) Both FAM237A and GPR83 are ubiquitously present in all vertebrates and are highly conserved from fish to mammals. Deorphanization of GPR83 will stimulate further functional studies of this important receptor. Preparation of mature FAM237A is quite difficult, because of its complex posttranslational modifications and hydrophobic nature. The present study also provided an efficient approach to prepare highly active FAM237A and thus will promote further functional studies.

Unfortunately, our present results did not support the hypothesis that PEN and proCCK56-63 are ligands of GPR83. Neither peptide displayed detectable binding and activating effects towards GPR83 in our assays. As shown in Fig. S1, GPR83 is ubiquitously present in all vertebrates and is highly conserved from fish to mammals. In contrast, the peptide PEN precursor PCSK1N/proSAAS is not so conserved in evolution (Fig. S6). Moreover, no PCSK1N/proSAAS orthologs can be found in birds in the Gene database at NCBI (https://ncbi.nlm.nih.gov/gene). Thus, the evolutionary conservation of GPR83 and PCSK1N/proSAAS are quite different.

Previous studies [10,14] mainly relied on radioligand binding assay to detect binding of PEN and proCCK56-63 with GPR83. However, their experiments had some problems. (1) The quality of their radioligand was poor, because its reported specific activity (54.1 Ci/mmol) was much lower than the theoretical value (2715 Ci/mmol) of the mono-iodinated radioligand. Thus, only ∼2% of Tyr-rPEN was labeled by iodine-125 in their experiments, and ∼98% of Tyr-rPEN was not radio-iodinated at all. The unlabeled peptide in the tracer would function as a competitor, and thus inhibit binding of the radioligand with receptors, resulting in incorrect B_max_ and K_d_ values. During radio-iodination, the authors used too much Tyr-rPEN (200 µg, equal to 81 nmol) to react with Na^125^I (probably 1.0 mCi, equal to 0.46 nmol), thus only a small percent (∼0.6%) of Tyr-rPEN could be labeled by iodine-125. (2) Their binding assay was conducted in 50 mM Tris-Cl (pH 7.4) buffer without bovine serum albumin (BSA). In fact, 0.5–1.0% BSA is typically essential to prevent non-specific binding of radioligands, especially for hydrophobic peptides, such as PEN. (3) Strangely, their measured B_max_ values for membranes from mouse hypothalamic, cultured Neuro2A cells, or CHO cells overexpressing human GPR83 were all at similar levels. Generally, overexpression can provide much higher expression levels than endogenous expression, thus the B_max_ value of CHO cells overexpressing human GPR83 should be much higher than that of hypothalamic and cultured Neuro2A cells. Moreover, the authors likely mis-calculated their B_max_ values (1000-fold higher), their reported values likely have a unit of fmol/mg, rather than pmol/mg, as judged from their reported cpm values in some figures. Thus, it seemed that pairing of GPR83 with peptide PEN and proCCK56-63 lacks solid experimental support.

## Experimental procedures

### Chemical synthesis of peptide PEN and proCCK56-63

The peptide PEN and proCCK56-63 were chemically synthesized at GL Biotech (Shanghai, China) via solid-phase peptide synthesis using standard Fmoc methodology. The synthetic crude peptides were applied to HPLC sequentially using a semi-preparative C_18_ reverse-phase column and an analytical C_18_ reverse-phase column (Zorbax 300SB-C18, 9.4 or 4.6 × 250 mm; Agilent Technologies, Santa Clara, CA, USA). These peptides were eluted from the reverse-phase columns by an acidic acetonitrile gradient composed of acidic solvent A and B. The acidic solvent A is 0.1% aqueous trifluoroacetic acid (TFA), and the acidic solvent B is acetonitrile containing 0.1% TFA. The eluted peptide fractions were manually collected, lyophilized, and confirmed by electrospray mass spectrometry. After purification, the lyophilized peptide powder was weighed using a precise balance and dissolved in 1.0 mM aqueous hydrochloride (pH3.0) as stock solution for later activity assays.

### Preparation of mature FAM237A via bacterial overexpression, intein-mediated amidation, and *in vitro* refolding

The nucleotide sequence encoding the C-terminally intein-fused human FAM237A (FAM237A-Intein) was chemically synthesized at GeneWiz (Suzhou, China) according to the published sequence of the gyrase A intein from *Mycobacterium xenopi* [16,17] and the codon-optimized human FAM237A sequence (Fig. S3). After cleavage with restriction enzymes NsiI and NotI, the synthetic DNA fragment was ligated into a pET vector, resulting in the expression construct pET/FAM237A-Intein-6×His (Fig. S3). The coding sequence of the N-terminally SmBiT-tagged FAM237A-Intein (SmBiT-FAM237A-Intein) was amplified by polymerase chain reaction (PCR) using pET/FAM237A-Intein-6×His as a template. After cleavage with restriction enzymes NsiI and NotI, the amplified DNA fragment was ligated into the pET vector, resulting in the expression construct pET/SmBiT-FAM237A-Intein-6×His (Fig. S3).

Thereafter, the FAM237A fusion proteins were overexpressed in *Escherichia coli* strain BL21(DE3) according to standard protocols. Briefly, the transformed cells were cultured in liquid LB medium (plus 100 µg/ml ampicillin) to OD_600_ = 1.5–2.0 at 37 °C with vigorous shaking, and then induced by 1.0 mM isopropyl β-D-thiogalactoside (IPTG) at 37 °C for 6–8 h. Subsequently, the cells were harvested by centrifugation (5000 *g*, 10 min), re-suspended in lysis buffer (20 mM phosphate buffer, pH 7.4, 0.5 M NaCl), and lysed by sonication. After centrifugation (15000 *g*, 30 min), inclusion bodies were collected and resuspended in solubilizing buffer (6 M guanidine chloride in lysis buffer) and subjected to *S*-sulfonation by adding solid sodium sulfite (Na_2_SO_3_) and sodium tetrathionate (Na_2_S_4_O_6_) to the final concentration of 100 mM and 80 mM, respectively. After shaking at room temperature for ∼4 h, the reaction mixture was subjected to centrifugation (15000 *g*, 30 min), and the supernatant was applied to an immobilized metal ion affinity chromatography (Ni^2+^ column). The *S*-sulfonated FAM237A fusion protein was eluted from the Ni^2+^ column by 250 mM imidazole (in solubilizing buffer), and its concentration in the eluent was calculated according to the measured ultra-violet (UV) absorbance at 280 nm and its empirical extinction coefficient (in unit of M^−1^cm^−1^) that was calculated according to its Trp and Tyr numbers (ε_280nm_ = 5500 × N_Trp_ + 1490 × N_Tyr_).

According to the measured concentration of the FAM237A fusion protein, tris(2-carboxyethyl)phosphine (TCEP) and sodium 2-mercptoethanesulfonate (MES) was added to the eluent to insure that the subsequent renaturation system contains 2.0 mM TCEP, 100 mM MES, and ∼0.15 mg/ml FAM237A fusion protein after dilution. After incubation at 37 °C for ∼1 h, the eluent was pre-cooled on ice and then diluted (typically over 10-fold) into the ice-cold renaturation buffer (0.5 M L-arginine, pH7.4). After overnight incubation at room temperature, solid ammonium bicarbonate (NH_4_HCO_3_) was added to the final concentration of 2.0 M and pH of the solution was adjusted to 8.5–9.0 by adding appropriate amount of aqueous ammonia. After stirred at room temperature for ∼3 h, the pellet was collected by centrifugation (12000 *g*,45 min).

The above pellet was dissolved in the solubilizing buffer and subjected to *S*-sulfonation again. After shaking at room temperature for ∼3 h, Tween-20 was added to the final concentration of 0.5%, and the solution was continuously shaken for ∼0.5 h. After centrifugation (12000 *g*, 10 min), the supernatant was applied to a Ni^2+^ column and the flow-through fraction was collected, acidified to pH3–4 by adding appropriate amount of TFA and subjected to HPLC purification using a C_4_ reverse-phase column (Hi-Pore reversed-phase column, 4.6 × 250 mm, Bio-Rad, Hercules, CA, USA). The *S*-sulfonated FAM237A was eluted from the C_4_ reverse-phase column by an acidic acetonitrile gradient composed of the acidic solvent A (0.1% aqueous TFA) and the acidic solvent B (acetonitrile plus 0.1% TFA). After lyophilization, the purified *S*-sulfonated FAM237A was dissolved in 1.0 mM aqueous hydrochloride (pH3.0), quantified by UV absorbance, and diluted into the ice-cold refolding buffer [0.5 M L-arginine, pH 8.5, 1.0 mM oxidative glutathione (GSSG), and 2.0 mM reduced glutathione (GSH)] to the final concentration of ∼0.2 mg/ml. After incubation on ice for ∼6 h, the refolding mixture was acidified to pH3–4 by adding appropriate amount of TFA and subjected to HPLC purification. The refolded mature FAM237A was eluted from the C_4_ reverse-phase column (Hi-Pore reversed-phase column, 4.6 × 250 mm, Bio-Rad) by the acidic acetonitrile gradient.

Purity of the wild-type or mutant FAM237A peptides was analyzed by HPLC using an analytical C_18_ reverse-phase column (Zorbax 300SB-C18, 4.6 × 250 mm, Agilent Technologies). The FAM237A peptides were eluted from the C_18_ reverse-phase column by an alkaline acetonitrile gradient composed of alkaline solvent A and B [15]. The alkaline solvent A is 20 mM aqueous NH_4_HCO_3_ (pH8.5) containing 20% (v/v) acetonitrile, the alkaline solvent B is 20 mM aqueous NH_4_HCO_3_ (pH8.5) containing 70% (v/v) acetonitrile. The purified wild-type or mutant FAM237A peptides were dissolved in 1.0 mM aqueous hydrochloride (pH3.0) and their concentrations were calculated according to the measured UV absorbance at 280 nm and the empirical extinction coefficient (ε_280nm_) of 30480 M^−1^cm^−1^. The FAM237A stock solutions were stored at -80°C for later activity assays.

### Labeling of FAM237A and PEN

To prepare fluorescent FAM237A for fluorescent microscopic visualization, FAM237A stock solution (50 µM in 1.0 mM aqueous HCl) and the fluorescent dye 5(6)-carboxytetramethylrhodamine N-succinimidyl ester (Macklin, Shanghai, China) stock solution (10 mM in acetonitrile) were mixed in phosphate buffer, and the final reaction system contained 25 µM FAM237A, 50 µM fluorescent dye, 100 mM phosphate buffer (pH8.0), and 33.3% (v/v) acetonitrile. After incubation at room temperature for 30 min, the reaction mixture was acidified to pH3–4 by adding TFA, applied to HPLC, and eluted from the C_4_ reverse-phase column (Hi-Pore reversed-phase column, 4.6 × 250 mm, Bio-Rad) by the acidic acetonitrile gradient.

The synthetic PEN was first modified by SPDP [3-(2-Pyridyldithio)propionic acid N-hydroxysuccinimide ester, from Sigma-Aldrich, St. Louis, MO, USA]. For this reaction, the PEN stock solution (600 µM in 1.0 mM aqueous HCl) and the SPDP stock solution (10 mM in acetonitrile) were mixed in the phosphate buffer, and the final reaction system contained 300 µM PEN, 3 mM SPDP, and 100 mM phosphate buffer (pH8.0). After incubation at room temperature for 30 min, the reaction mixture was acidified to pH3–4 by adding TFA, applied to HPLC, and eluted from the C_18_ reverse-phase column (Zorbax 300SB-C18, 4.6 × 250 mm, Agilent Technologies) by the acidic acetonitrile gradient. After lyophilization, the SPDP-modified PEN was reacted with cysteamine in the reaction system containing 200 µM SPDP-modified PEN, 600 mM cysteamine, and 100 mM phosphate buffer (pH8.0). After incubation at room temperature for 30 min, the reaction mixture was acidified to pH3–4 by adding TFA, and purified by HPLC using the C_18_ reverse-phase column (Zorbax 300SB-C18, 4.6 × 250 mm, Agilent Technologies). Finally, the above modified PEN was further reacted with the fluorescent dye 5(6)-carboxytetramethylrhodamine N-succinimidyl ester in the reaction system containing 280 µM modified PEN, 1.0 mM fluorescent dye, 100 mM phosphate buffer (pH8.0), and 30% (v/v) acetonitrile. After incubation at room temperature for 30 min, the reaction mixture was acidified to pH3–4, and purified by HPLC using the C_18_ reverse-phase column (Zorbax 300SB-C18, 4.6 × 250 mm, Agilent Technologies).

After lyophilization, the tetramethylrhodamine (TMR)-labeled peptides, TMR-FAM237A and TMR-PEN, were dissolved in 1.0 mM aqueous hydrochloride (pH3.0) and their concentrations were calculated from the measured absorbance at 553 nm and the empirical extinction coefficient of TMR (ε_530nm_ = 92000 M^−1^cm^−1^). The stock solutions were stored at -80°C for later activity assays.

To prepare the SmBiT-PEN tracer, the SPDP-modified PEN was reacted with a synthetic C-terminally Cys-tagged SmBiT (SmBiT-Cys) in the reaction system containing 200 µM SPDP-modified PEN, 400 µM SmBiT-Cys, and 20 mM phosphate buffer (pH8.0). After incubation at room temperature for 30 min, the reaction mixture was acidified to pH3–4, applied to HPLC, and eluted from the C_18_ reverse-phase column (Zorbax 300SB-C18, 4.6 × 250 mm, Agilent Technologies) by the acidic acetonitrile gradient.

The C-terminally Cys-tagged NanoLuc (Luc-Cys) was designed in our previous study for NanoLuc conjugation with various peptides carrying a free Cys residue [24]. After overexpression in *E. coli*, Luc-Cys was purified according to previous procedure [24]. For conjugation, the SDSP-modified PEN and the purified Luc-Cys [eluted from a DEAE column by NaCl gradient in 20 mM phosphate buffer (pH8.0)] were mixed together in reaction system containing 20 µM Luc-Cys, 40 µM SPDP-modified PEN, and 20 mM phosphate buffer (pH8.0). After incubation at room temperature for 30 min, the reaction mixture was loaded onto a spin Ni^2+^ column. After washing by 20 mM phosphate buffer (pH8.0), the PEN-Luc conjugate was eluted by 250 mM imidazole, analyzed by SDS-PAGE and bioluminescence, and stored at -80 °C for later binding assay.

### Generation of the expression constructs for human GPR83

The cDNA of human GPR83 was purchased from Immunogen (Shanghai, China), its nucleotide sequence is identical to the reference sequence (NM_016540) except for a synonymous mutation. The full-length coding region (without stop codon) of human GPR83 was PCR amplified, cleaved by restriction enzymes NheI and AgeI, and then ligated into a modified pcDNA6 vector or modified pTRE3G-BI vectors. The resultant construct pcDNA6/GPR83 expresses a full-length human GPR83 (Fig. S4), the construct pTRE3G-BI/GPR83-EGFP expresses a C-terminally EGFP-fused full-length human GPR83, and the construct pTRE3G-BI/GPR83-IRES-EGFP coexpresses a full-length GPR83 and a free EGFP mediated by an IRES element (Fig. S4). The coding region (with stop codon) of the mature human GPR83 (17–423 residues) was PCR amplified, cleaved by restriction enzymes KpnI and AgeI, and then ligated into a pcDNA6/sLgBiT vector, resulting in the construct pcDNA6/sLgBiT-GPR83 encoding an N-terminally sLgBiT-fused mature GPR83 (Fig. S4). To generate the C-terminally LgBiT-fused GPR83 (GPR83-LgBiT), the coding region of full-length human GPR83 (without stop codon) and the coding region of LgBiT were PCR amplified separately. Thereafter, the partially overlapped two DNA fragments were annealed together and PCR amplified again. The resultant full-length coding fragment for GPR83-LgBiT was cleaved by restriction enzymes EcoRI and NdeI, and ligated into the multiple cloning site 2 (MCS2) of the pTRE3G-BI vector (Clontech, Mountain View, CA, USA), resulting in the construct pTRE3G-BI/GPR83-LgBiT. The coding region of the N-terminally SmBiT-fused β-arrestin 2 (SmBiT-ARRB2) was PCR amplified using the human β-arrestin 2 (NM_004313) expression construct pENTER/ARRB2 (WZ Biosciences Inc., Jinan, China) as a template. The amplified DNA fragment was cleaved by restriction enzymes BamHI and NotI, and ligated into the MCS1 of pTRE3G-BI/GPR83-LgBiT, resulting in the construct pTRE3G-BI/GPR83-LgBiT:SmBiT-ARRB2 that coexpresses GPR83-LgBiT and SmBiT-ARRB2 under a controllable manner via a bidirectional promoter (Fig. S5).

### The NanoBiT-based ligand–receptor binding assays

The expression construct pcDNA6/sLgBiT-GPR83 was transiently transfected into HEK293T cells using the transfection reagent Lipo8000 (Beyotime Technology, Shanghai, China). Next day, the transfected cells were trypisinized, seeded into white opaque 96-well plates, and continuously cultured for ∼24 h to ∼90% confluence. To conduct the binding assays, the medium was removed and the binding solution (serum-free DMEM plus 0.1% BSA and 0.01% Tween-20) was added to the living cells (50 µl/well). For kinetic binding assay, the binding solution contains 1.0 µl of NanoLuc substrate stock solution (Promega, Madison, WI, USA). After addition to one well, bioluminescence data were immediately collected for ∼10 min on a SpectraMax iD3 plate reader (Molecular Devices, Sunnyvale, CA, USA) with an interval of 10 s. Thereafter, 2.0 µl of SmBiT-FAM237A stock solution was added to the final concentration of 2.5 nM, and bioluminescence data were continuously collected for ∼20 min with an interval of 10 s. Finally, 2.0 µl of mature FAM237A stock solution was added to the final concentration of 1.0 µM, and bioluminescence data were continuously collected for ∼30 min with an interval of 10 s. For saturation binding assays, the binding solution contains varied concentrations of SmBiT-FAM237A or SmBiT-PEN; for competition binding assays, the binding solution contains 2.5 nM of SmBiT-FAM237A and varied concentrations of competitor. After incubation at 22°C for ∼30 min, diluted NanoLuc substrate was added (10 µl/well, 30-fold dilution using the binding solution) and bioluminescence was immediately measured on a SpectraMax iD3 plate reader (Molecular Devices). The equilibrium binding data were expressed as mean ± standard deviation (SD) (*n* = 3), and fitted to one-site binding model by SigmaPlot 10.0 software (SYSTAT software, Chicago, IL, USA).

### Visualization of fluorescent ligand binding to GPR83

The Tet-On inducible expression construct pTRE3G-BI/GPR83-EGFP or pTRE3G-BI/GPR83-IRES-EGFP was transiently transfected into HEK293T cells together with the expression control vector pCMV-TRE3G (Clontech) using the transfection reagent Lipo8000 (Beyotime Technology). Next day, the transfected cells were trypsinized, suspended in the induction medium (complete DMEM medium plus 10 ng/ml Dox), seeded into 35-mm glass-bottomed dishes, and continuously cultured at 37 °C for ∼24 h. Thereafter, the induction medium was removed and the binding solution (serum-free DMEM medium plus 1% BSA) with or without 5.0 nM TMR-labeled peptide was added. After continuously cultured at 37 °C for ∼2 h, the living cells were observed under an SP8 confocal microscope (Leica Microsystems, Wetzlar, Germany).

### NanoBiT-based β-arrestin recruitment assay

The coexpression construct pTRE3G-BI/GPR83-LgBiT:SmBiT-ARRB2 and the expression control vector pCMV-TRE3G (Clontech) were cotransfected into HEK293T cells using the transfection reagent Lipo8000 (Beyotime Technology). Next day, the transfected cells were trypisinized, suspended in the induction medium (complete DMEM medium plus 2.5 ng/ml of Dox), seeded into white opaque 96-well plates, and continuously cultured at 37 °C for ∼24 h. Before the activation assay, the cells were placed at room temperature for ∼30 min. To conduct the assay, the medium was removed and pre-warmed activation solution (serum-free DMEM plus 1% BSA) containing NanoLuc substrate was added to the living cells (40 µl/well, containing 1.0 µl of NanoLuc substrate stock from Promega). Thereafter, bioluminescence data were immediately collected for ∼4 min on a SpectraMax iD3 plate reader (Molecular Devices) with an interval of 15 s. Subsequently, peptide solution (diluted in the activation solution) was added to these wells (10 µl/well), and bioluminescence data were continuously collected for ∼4 min with an interval of 15 s. The measured absolute signals were corrected for inter well variability by forcing all curves after addition of NanoLuc substrate (without ligand) to same level. The values of area under curve (AUC) after addition of ligands were then calculated based on these corrected activation curves by SigmaPlot 10.0 software (SYSTAT software). The calculated activation data were expressed as mean ± SD (*n* =3), and fitted to a sigmoidal curve by SigmaPlot 10.0 software (SYSTAT software).

## Abbreviations

BSA: bovine serum albumin
Dox: doxycycline
EGFP: enhanced green fluorescent protein
FAM237A: family with sequence similarity 237 member A
GPCR: G protein-coupled receptor
GPR83: G protein-coupled receptor 83
GSH: reduced glutathione
GSSG: oxidative glutathione
HPLC: high performance liquid chromatography
IPTG: isopropyl β-D-thiogalactoside
IRES: internal ribosome entry site
LgBiT: large NanoLuc fragment for NanoBiT
MCS: multiple cloning site
MES: sodium 2-mercptoethanesulfonate
NanoBiT: NanoLuc Binary Technology
PCR: polymerase chain reaction
PCSK1N: proprotein convertase subtilisin/kexin type 1 inhibitor
SDS-PAGE: sodium dodecyl sulfate-polyacrylamide gel electrophoresis
sLgBiT: secretory LgBiT
SmBiT: low-affinity small complementation tag for NanoBiT
SPDP: 3-(2-Pyridyldithio)propionic acid N-hydroxysuccinimide ester
TCEP: tris(2-carboxyethyl)phosphine
TFA: trifluoroacetic acid
TMD: transmembrane domain
TMR: tetramethylrhodamine
UV: ultra-violet.

## Acknowledgments

We thank former laboratory members, Ben-Jun Ji, Yu Liu, Ge Song, Lei Zhang, Yu-Qi Guo, Qing-Ping Wu, Li-Li Shou, and Ning Li, for generation of over 200 GPCR expression constructs. This work was supported by grants from the National Natural Science Foundation of China (31971193).

