## Supplementary Fig. S1-S6 for "FAM237A, rather than peptide PEN and proCCK56-63, is a ligand of the orphan receptor GPR83"

**Contents:**

**Fig. S1**. Amino acid sequence alignment of GPR83 orthologs from fish to mammals.

**Fig. S2.** Amino acid sequence alignment of FAM237A orthologs from fish to mammals.

**Fig. S3.** The nucleotide and amino acid sequences of human FAM237A constructs overexpressed in *E. coli*.

**Fig. S4.** The nucleotide and amino acid sequences of human GPR83 constructs expressed in mammalian cells.

**Fig. S5.** The nucleotide and amino acid sequences of GPR83-LgBiT and SmBiT-ARRB2 in pTRE3G-BI vector for coexpression in mammalian cells under a controllable manner.

**Fig. S6.** Amino acid sequence alignment of PCSK1N/proSAAS orthologs from different species.

Alignment of GPR83 orthologs from fish to mammals

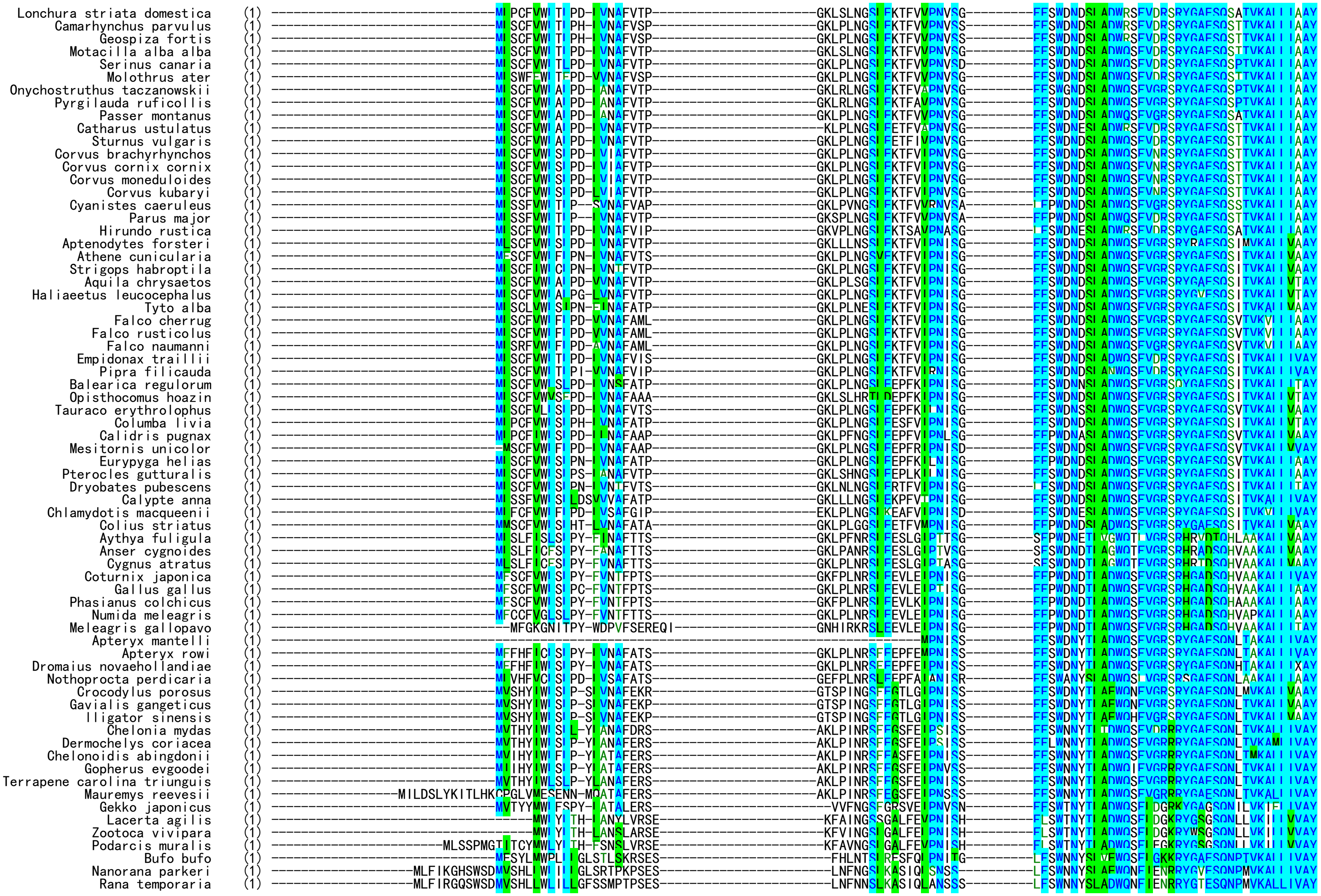

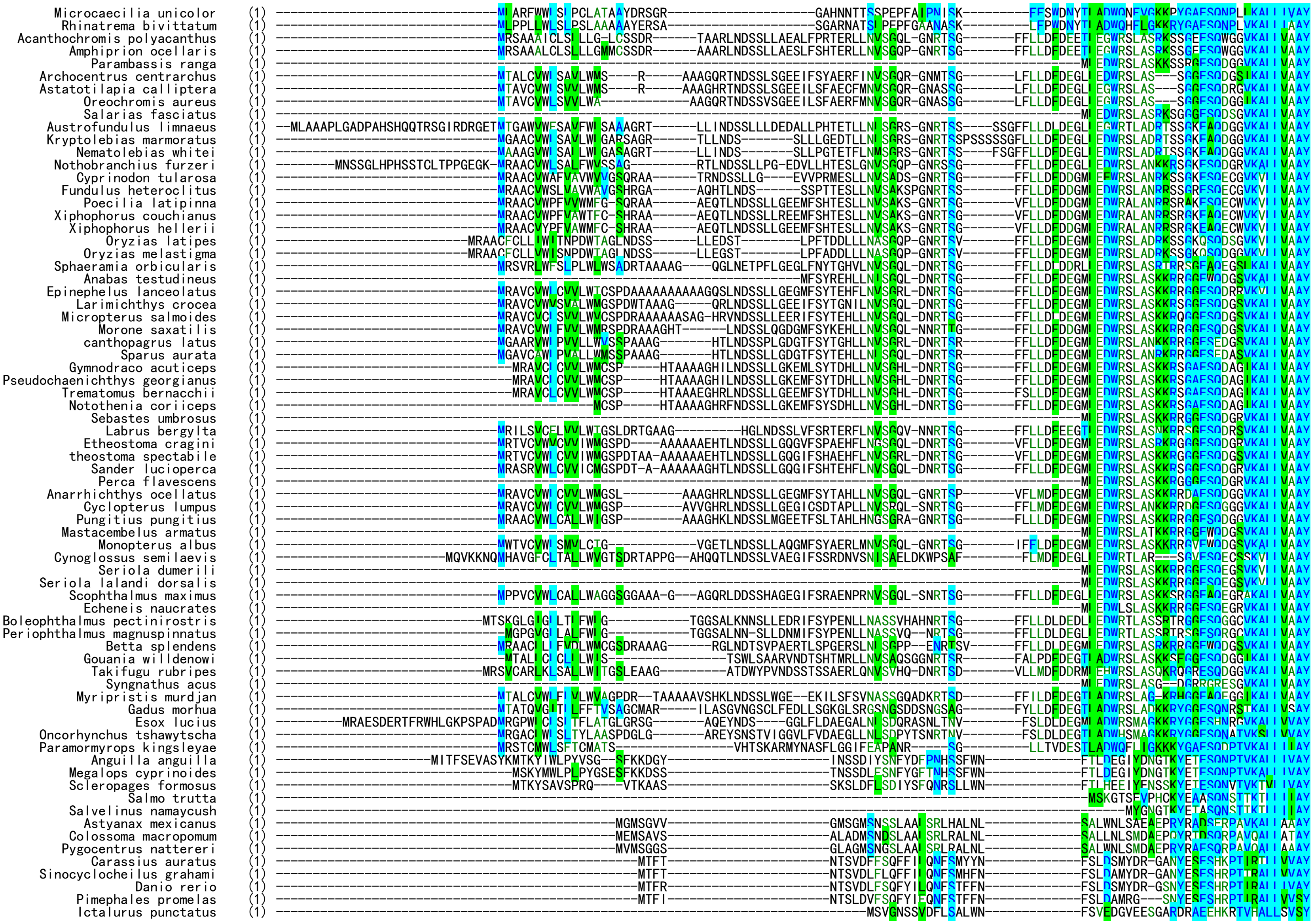
Microcaecilia unicolor
Rhinatrema bivittatum
Acanthochromis polyacanthus
Amphiprion ocellaris
Parambassis ranga
Archocentrus centrarchus
Astatotilapia calliptera
Oreochromis aureus
Salarias fasciatus
Austrofundulus limnaeus
Kryptolebias marmoratus
Nematolebias whitei
Nothobranchius furzeri
Cyprinodon tularosa
Fundulus heteroclitus
Poecilia latipinna
Xiphophorus couchianus
Xiphophorus hellerii
Oryzias latipes
Oryzias melastigma
Sphaeramia orbicularis
Anabas testudineus
Epinephelus lanceolatus
Larimichthys crocea
Micropterus salmoides
Morone saxatilis
canthopagrus latus
Sparus aurata
Gymnodraco acuticeps
Pseudochaenichthys georgianus
Trematomus bernacchii
Notothenia coriiceps
Sebastes umbrosus
Labrus bergylta
Etheostoma cragini
theostoma spectabile
Sander lucioperca
Perca flavescens
Anarrhichthys ocellatus
Cyclopterus lumpus
Pungitius pungitius
Mastacembelus armatus
Monopterus albus
Cynoglossus semilaevis
Seriola dumerili
Seriola lalandi dorsalis
Scophthalmus maximus
Echeneis naucrates
Boleophthalmus pectinirostris
Periophthalmus magnuspinnatus
Betta splendens
Gouania willdenowi
Takifugu rubripes
Syngnathus acus
Myripristis murdjan
Gadus morhua
Esox lucius
Oncorhynchus tshawytscha
Paramormyrops kingsleyae
Anguilla anguilla
Megalops cyprinoides
Scleropages formosus
Salmo trutta
Salvelinus namaycush
Astyanax mexicanus
Colossoma macropomum
Pygocentrus nattereri
Carassius auratus
Sinocyclocheilus grahami
Danio rerio
Pimephales promelas
Ictalurus punctatus

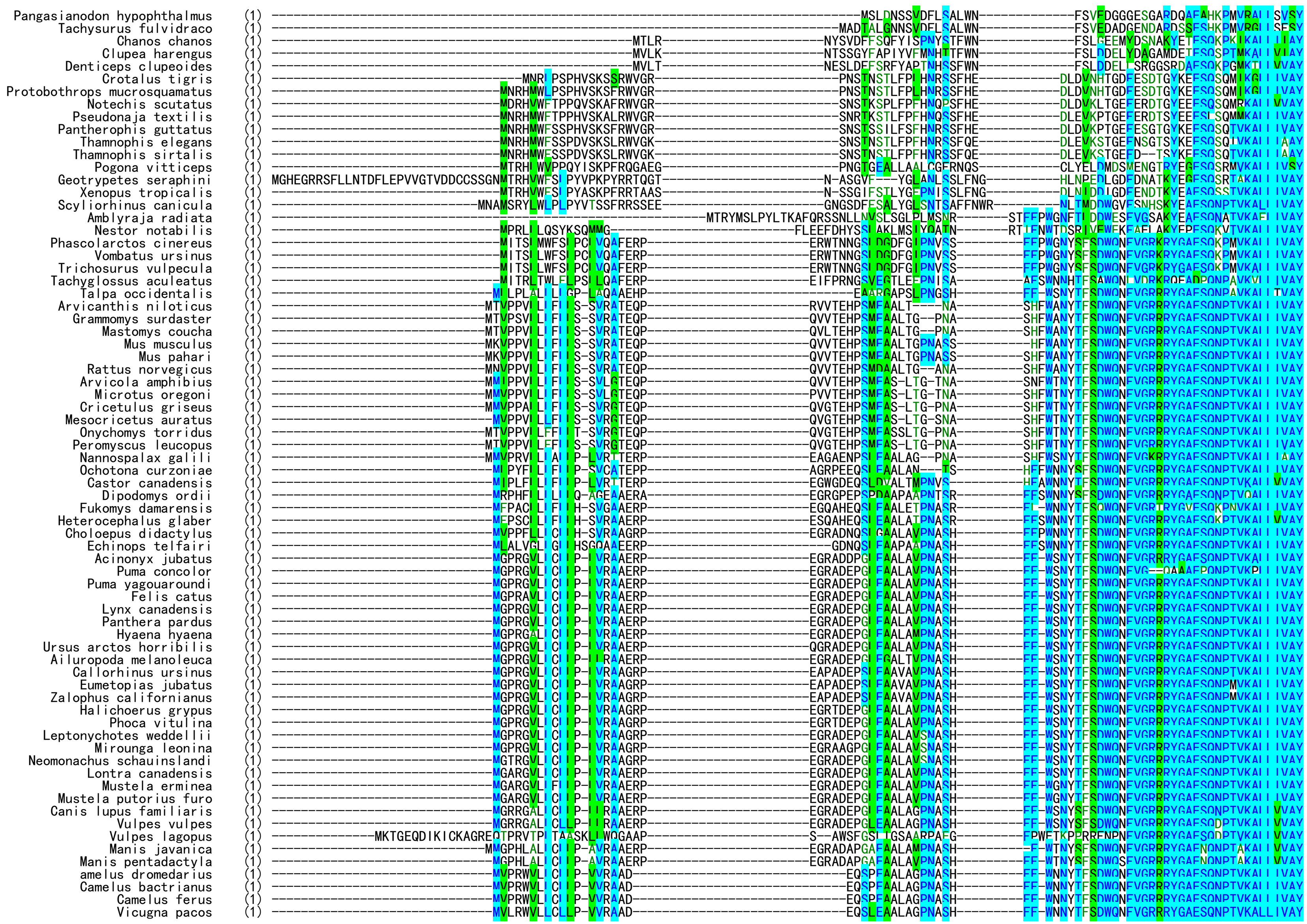
Pangasianodon hypophthalmus
Tachysurus fulvidraco
Chanos chanos
Clupea harengus
Denticeps clupeoides
Crotalus tigris
Protobothrops mucrosquamatus
Notechis scutatus
Pseudonaja textilis
Pantherophis guttatus
Thamnophis elegans
Thamnophis sirtalis
Pogona vitticeps
Geotrypetes seraphini
Xenopus tropicalis
Scyliorhinus canicula
Amblyraja radiata
Nestor notabilis
Phascolarctos cinereus
Vombatus ursinus
Trichosurus vulpecula
Tachyglossus aculeatus
Talpa occidentalis
Arvicanthis niloticus
Grammomys surdaster
Mastomys coucha
Mus musculus
Mus pahari
Rattus norvegicus
Arvicola amphibius
Microtus oregoni
Cricetulus griseus
Mesocricetus auratus
Onychomys torridus
Peromyscus leucopus
Nannospalax galili
Ochotona curzoniae
Castor canadensis
Dipodomys ordii
Fukomys damarensis
Heterocephalus glaber
Choloepus didactylus
Echinops telfairi
Acinonyx jubatus
Puma concolor
Puma yagouaroundi
Felis catus
Lynx canadensis
Panthera pardus
Hyaena hyaena
Ursus arctos horribilis
Ailuropoda melanoleuca
Callorhinus ursinus
Eumetopias jubatus
Zalophus californianus
Halichoerus grypus
Phoca vitulina
Leptonychotes weddellii
Mirounga leonina
Neomonachus schauinslandi
Lontra canadensis
Mustela erminea
Mustela putorius furo
Canis lupus familiaris
Vulpes vulpes
Vulpes lagopus
Manis javanica
Manis pentadactyla
amelus dromedarius
Camelus bactrianus
Camelus ferus
Vicugna pacos

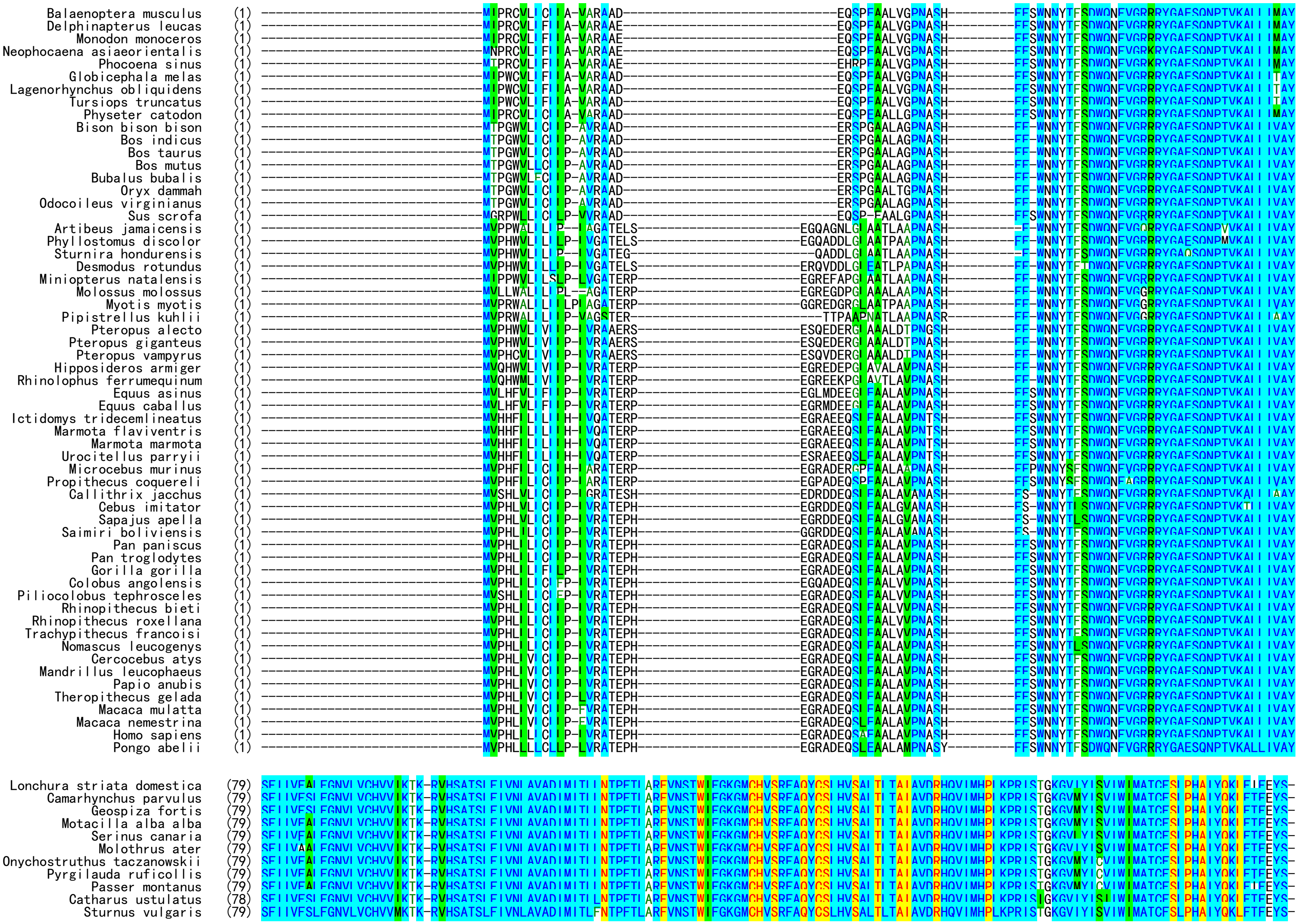
Balaenoptera musculus
Delphinapterus leucas
Monodon monoceros
Neophocaena asiaeorientalis
Phocoena sinus
Globicephala melas
Lagenorhynchus obliquidens
Tursiops truncatus
Physeter catodon
Bison bison bison
Bos indicus
Bos taurus
Bos mutus
Bubalus bubalis
Oryx dammah
Odocoileus virginianus
Sus scrofa
Artibeus jamaicensis
Phyllostomus discolor
Sturnira hondurensis
Desmodus rotundus
Miniopterus natalensis
Molossus molossus
Myotis myotis
Pipistrellus kuhlii
Pteropus alecto
Pteropus giganteus
Pteropus vampyrus
Hipposideros armiger
Rhinolophus ferrumequinum
Equus asinus
Equus caballus
Ictidomys tridecemlineatus
Marmota flaviventris
Marmota marmota
Urocitellus parryii
Microcebus murinus
Propithecus coquereli
Callithrix jacchus
Cebus imitator
Sapajus apella
Saimiri boliviensis
Pan paniscus
Pan troglodytes
Gorilla gorilla
Colobus angolensis
Piliocolobus tephrosceles
Rhinopithecus bieti
Rhinopithecus roxellana
Trachypithecus francoisi
Nomascus leucogenys
Cercocebus atys
Mandrillus leucophaeus
Papio anubis
Theropithecus gelada
Macaca mulatta
Macaca nemestrina
Homo sapiens
Pongo abelii
Lonchura striata domestica
Camarhynchus parvulus
Geospiza fortis
Motacilla alba alba
Serinus canaria
Molothrus ater
Onychostruthus taczanowskii
Pyrgilauda ruficollis
Passer montanus
Catharus ustulatus
Sturnus vulgaris

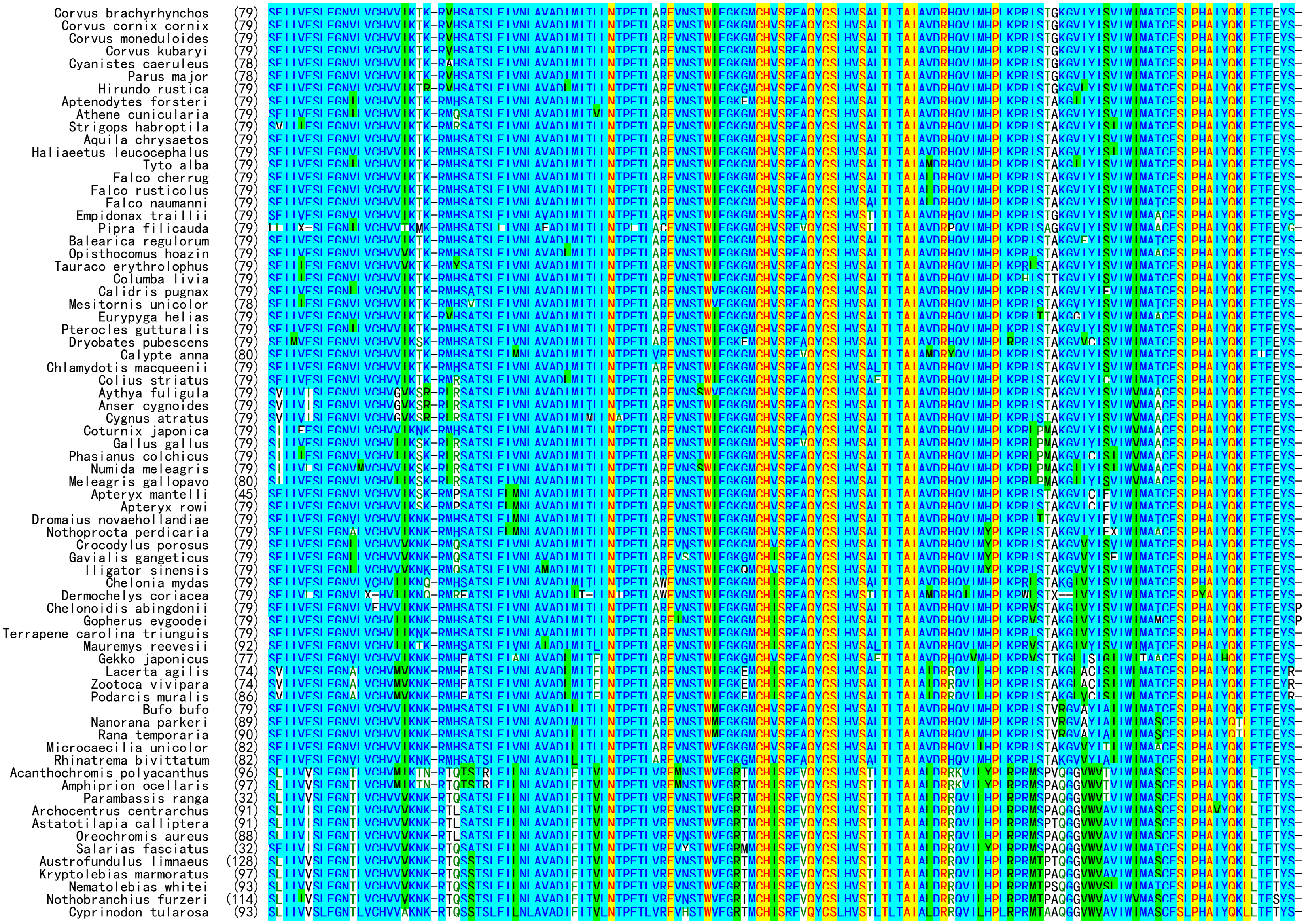
Corvus brachyrhynchos (79)
Corvus cornix cornix (79)
Corvus monedulоides (79)
Corvus kubaryi (79)
Cyanistes caeruleus (78)
Parus major (78)
Hirundo rustica (79)
Aptenodytes forsteri (79)
Athene cunicularia (79)
Strigops habroptila (79)
Aquila chrysaetos (79)
Haliaeetus leucocephalus (79)
Tyto alba (79)
Falco cherrug (79)
Falco rusticolus (79)
Falco naumanni (79)
Empidonax traillii (79)
Pipra filicauda (79)
Balearica regulorum (79)
Opisthocomus hoazin (79)
Tauraco erythrolophus (79)
Columba livia (79)
Calidris pugnax (79)
Mesitornis unicolor (78)
Eurypyga helias (79)
Pterocles gutturalis (79)
Dryobates pubescens (79)
Calypte anna (80)
Chlamydotis macqueenii (79)
Colius striatus (79)
Aythya fuligula (79)
Anser cygnoides (79)
Cygnus atratus (79)
Coturnix japonica (79)
Gallus gallus (79)
Phasianus colchicus (79)
Numida meleagris (79)
Meleagris gallopavo (80)
Apteryx mantelli (45)
Apteryx rowi (79)
Dromaius novaehollandiae (79)
Nothoprocta perdicaria (79)
Crocodylus porosus (79)
Gavialis gangeticus (79)
lligator sinensis (79)
Chelonia mydas (79)
Dermochelys coriacea (79)
Chelonoidis abingdonii (79)
Gopherus evgoodei (79)
Terrapene carolina triunguis (79)
Mauremys reevesii (92)
Gekko japonicus (77)
Lacerta agilis (74)
Zootoca vivipara (74)
Podarcis muralis (86)
Bufo bufo (79)
Nanorana parkeri (89)
Rana temporaria (90)
Microcaecilia unicolor (82)
Rhinatrema bivittatum (82)
Acanthochromis polyacanthus (96)
Amphiprion ocellaris (97)
Parambassis ranga (32)
Archocentrus centrarchus (91)
Astatotilapia calliptera (91)
Oreochromis aureus (88)
Salarias fasciatus (32)
Austrofundulus limnaeus (128)
Kryptolebias marmoratus (97)
Nematolebias whitei (93)
Nothobranchius furzeri (114)
Cyprinodon tularosa (93)

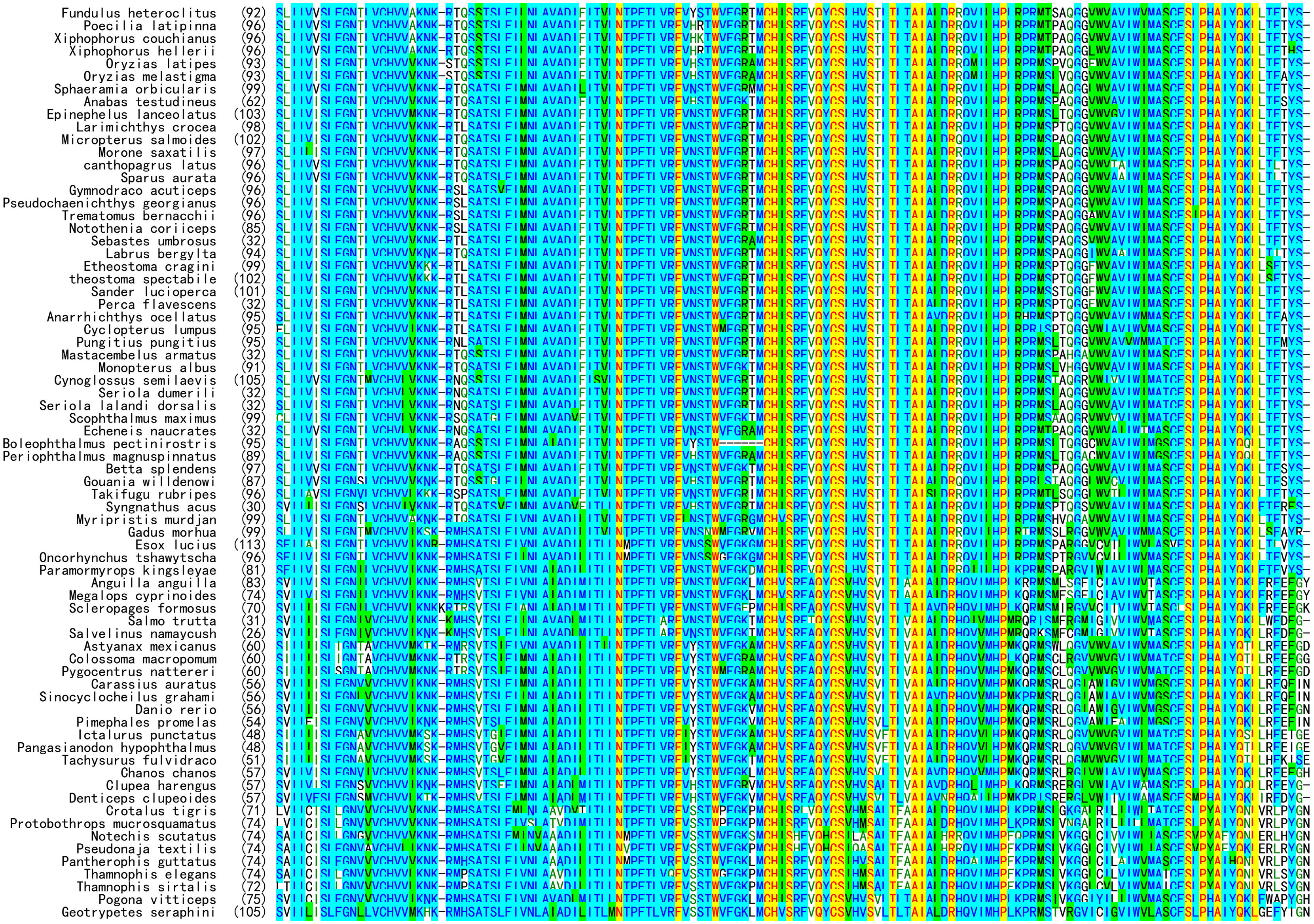

Fundulus heteroclitus (92)
Poecilia latipinna (96)
Xiphophorus couchianus (96)
Xiphophorus hellerii (96)
Oryzias latipes (93)
Oryzias melastigma (93)
Sphaeramia orbicularis (99)
Anabas testudineus (62)
Epinephelus lanceolatus (103)
Larimichthys crocea (98)
Micropterus salmoides (102)
Morone saxatilis (97)
canthopagrus latus (96)
Sparus aurata (96)
Gymnodraco acuticeps (96)
Pseudochaenichthys georgianus (96)
Trematomus bernacchii (96)
Notothenia coriiceps (85)
Sebastes umbrosus (32)
Labrus bergylta (94)
Etheostoma cragini (99)
theostoma spectabile (102)
Sander lucioperca (101)
Perca flavescens (32)
Anarrhichthys ocellatus (95)
Cyclopterus lumpus (95)
Pungitius pungitius (95)
Mastacembelus armatus (32)
Monopterus albus (91)
Cynoglossus semilaevis (105)
Seriola dumerili (32)
Seriola lalandi dorsalis (32)
Scophthalmus maximus (99)
Echeneis naucrates (32)
Boleophthalmus pectinirostris (95)
Periophthalmus magnuspinnatus (89)
Betta splendens (97)
Gouania willdenowi (87)
Takifugu rubripes (96)
Syngnathus acus (30)
Myripristis murdjan (99)
Gadus morhua (99)
Esox lucius (113)
Oncorhynchus tshawytscha (96)
Paramormyrops kingsleyae (81)
Anguilla anguilla (83)
Megalops cyprinoides (74)
Scleropages formosus (70)
Salmo trutta (31)
Salvelinus namaycush (26)
Astyanax mexicanus (60)
Colossoma macropomum (60)
Pygocentrus nattereri (60)
Carassius auratus (56)
Sinocyclocheilus grahami (56)
Danio rerio (56)
Pimephales promelas (54)
Ictalurus punctatus (48)
Pangasianodon hypophthalmus (48)
Tachysurus fulvidraco (51)
Chanos chanos (57)
Clupea harengus (57)
Denticeps clupeoides (57)
Crotalus tigris (71)
Protobothrops mucrosquamatus (74)
Notechis scutatus (74)
Pseudonaja textilis (74)
Pantherophis guttatus (74)
Thamnophis elegans (74)
Thamnophis sirtalis (72)
Pogona vitticeps (75)
Geotrypetes seraphini (105)

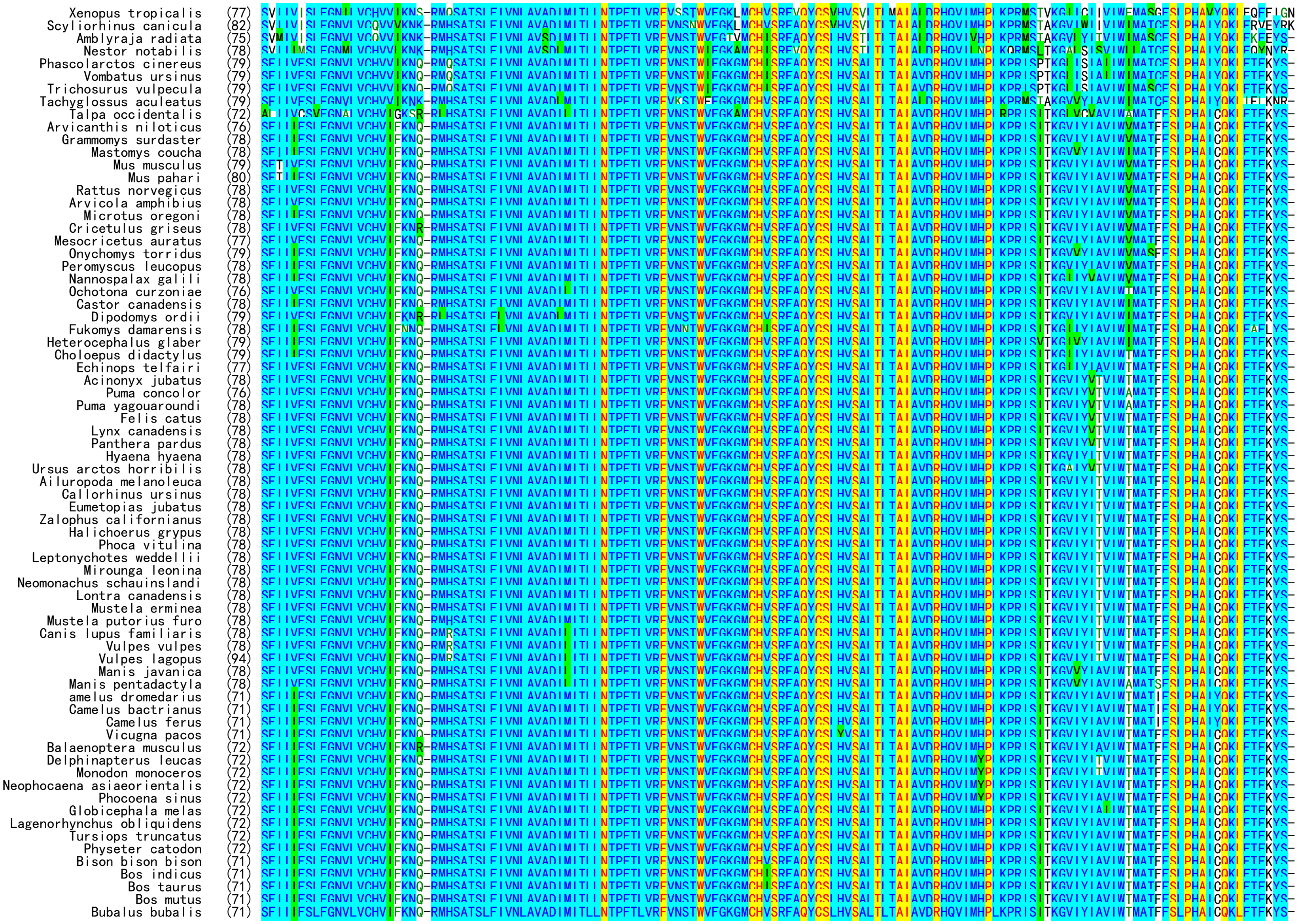

Xenopus tropicalis (77)
Scyliorhinus canicula (82)
Amblyraja radiata (75)
Nestor notabilis (78)
Phascolarctos cinereus (79)
Vombatus ursinus (79)
Trichosurus vulpecula (79)
Tachyglossus aculeatus (79)
Talpa occidentalis (72)
Arvicanthis niloticus (76)
Grammomys surdaster (78)
Mastomys coucha (78)
Mus musculus (79)
Mus pahari (80)
Rattus norvegicus (78)
Arvicola amphibius (78)
Microtus oregoni (78)
Cricetulus griseus (78)
Mesocricetus auratus (77)
Onychomys torridus (79)
Peromyscus leucopus (78)
Nannospalax galili (78)
Ochotona curzoniae (76)
Castor canadensis (78)
Dipodomys ordii (79)
Fukomys damarensis (78)
Heterocephalus glaber (79)
Choloepus didactylus (79)
Echinops telfairi (77)
Acinonyx jubatus (78)
Puma concolor (76)
Puma yagouaroundi (78)
Felis catus (78)
Lynx canadensis (78)
Panthera pardus (78)
Hyaena hyaena (78)
Ursus arctos horribilis (78)
Ailuropoda melanoleuca (78)
Callorhinus ursinus (78)
Eumetopias jubatus (78)
Zalophus californianus (78)
Halichoerus grypus (78)
Phoca vitulina (78)
Leptonychotes weddellii (78)
Mirounga leonina (78)
Neomonachus schauinslandi (78)
Lontra canadensis (78)
Mustela erminea (78)
Mustela putorius furo (78)
Canis lupus familiaris (78)
Vulpes vulpes (78)
Vulpes lagopus (94)
Manis javanica (78)
Manis pentadactyla (78)
amelus dromedarius (71)
Camelus bactrianus (71)
Camelus ferus (71)
Vicugna pacos (71)
Balaenoptera musculus (72)
Delphinapterus leucas (72)
Monodon monoceros (72)
Neophocaena asiaeorientalis (72)
Phocoena sinus (72)
Globicephala melas (72)
Lagenorhynchus obliquidens (72)
Tursiops truncatus (72)
Physeter catodon (72)
Bison bison bison (71)
Bos indicus (71)
Bos taurus (71)
Bos mutus (71)
Bubalus bubalis (71)

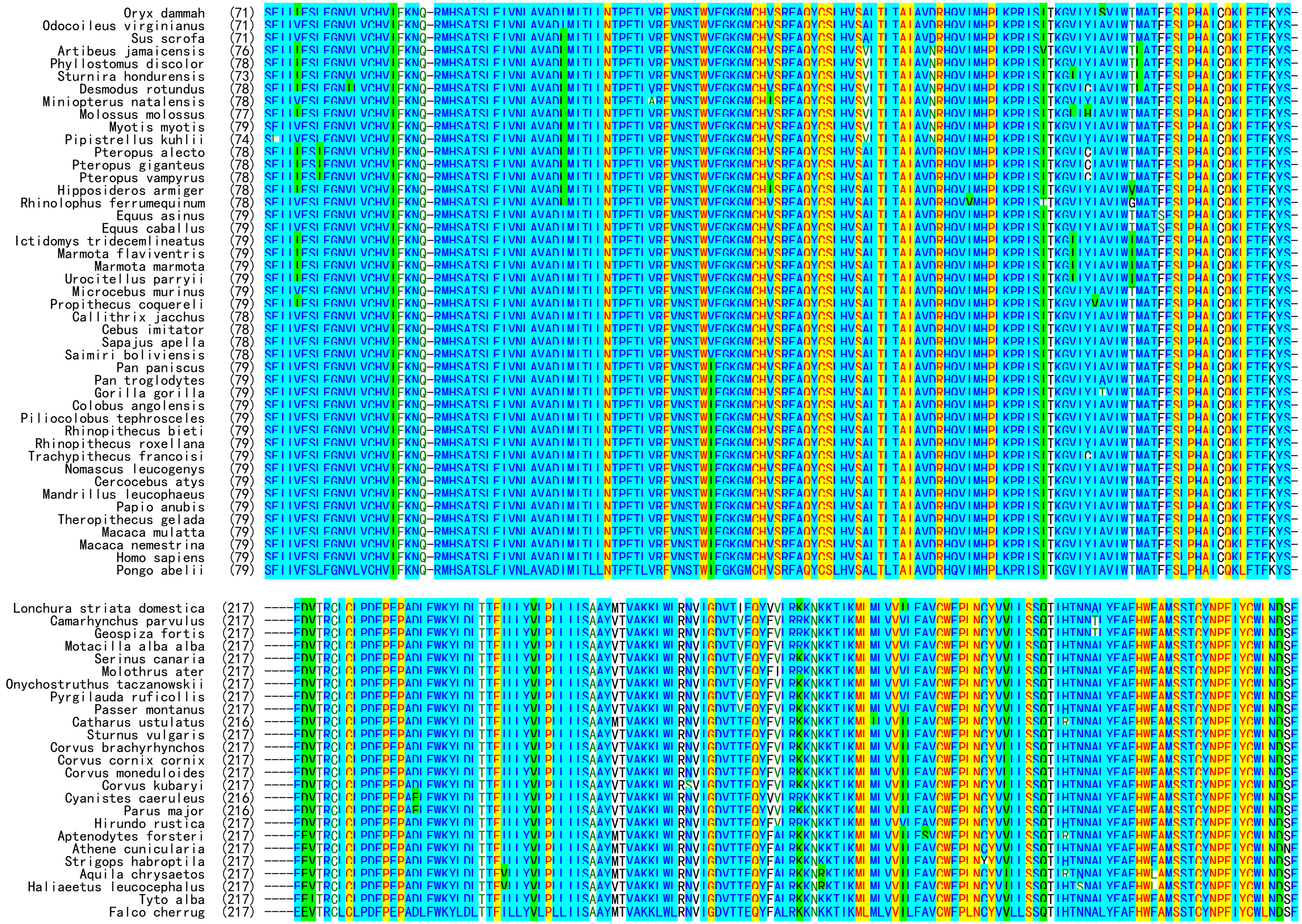

Oryx dammah (71)
Odocoileus virginianus (71)
Sus scrofa (71)
Artibeus jamaicensis (76)
Phyllostomus discolor (78)
Sturnira hondurensis (73)
Desmodus rotundus (78)
Miniopterus natalensis (78)
Molossus molossus (77)
Myotis myotis (79)
Pipistrellus kuhlii (74)
Pteropus alecto (78)
Pteropus giganteus (78)
Pteropus vampyrus (78)
Hipposideros armiger (78)
Rhinolophus ferrumequinum (78)
Equus asinus (79)
Equus caballus (79)
Ictidomys tridecemlineatus (79)
Marmota flaviventris (79)
Marmota marmota (79)
Urocitellus parryii (79)
Microcebus murinus (79)
Propithecus coquereli (79)
Callithrix jacchus (78)
Cebus imitator (78)
Sapajus apella (78)
Saimiri boliviensis (78)
Pan paniscus (79)
Pan troglodytes (79)
Gorilla gorilla (79)
Colobus angolensis (79)
Piliocolobus tephrosceles (79)
Rhinopithecus bieti (79)
Rhinopithecus roxellana (79)
Trachypithecus francoisi (79)
Nomascus leucogenys (79)
Cercocebus atys (79)
Mandrillus leucophaeus (79)
Papio anubis (79)
Theropithecus gelada (79)
Macaca mulatta (79)
Macaca nemestrina (79)
Homo sapiens (79)
Pongo abelii (79)
Lonchura striata domestica (217)
Camarhynchus parvulus (217)
Geospiza fortis (217)
Motacilla alba alba (217)
Serinus canaria (217)
Molothrus ater (217)
Onychostruthus taczanowskii (217)
Pyrgilauda ruficollis (217)
Passer montanus (217)
Catharus ustulatus (216)
Sturnus vulgaris (217)
Corvus brachyrhynchos (217)
Corvus cornix cornix (217)
Corvus moneduloides (217)
Corvus kubaryi (217)
Cyanistes caeruleus (216)
Parus major (216)
Hirundo rustica (217)
Aptenodytes forsteri (217)
Athene cunicularia (217)
Strigops habroptila (217)
Aquila chrysaetos (217)
Haliaeetus leucocephalus (217)
Tyto alba (217)
Falco cherrug (217)

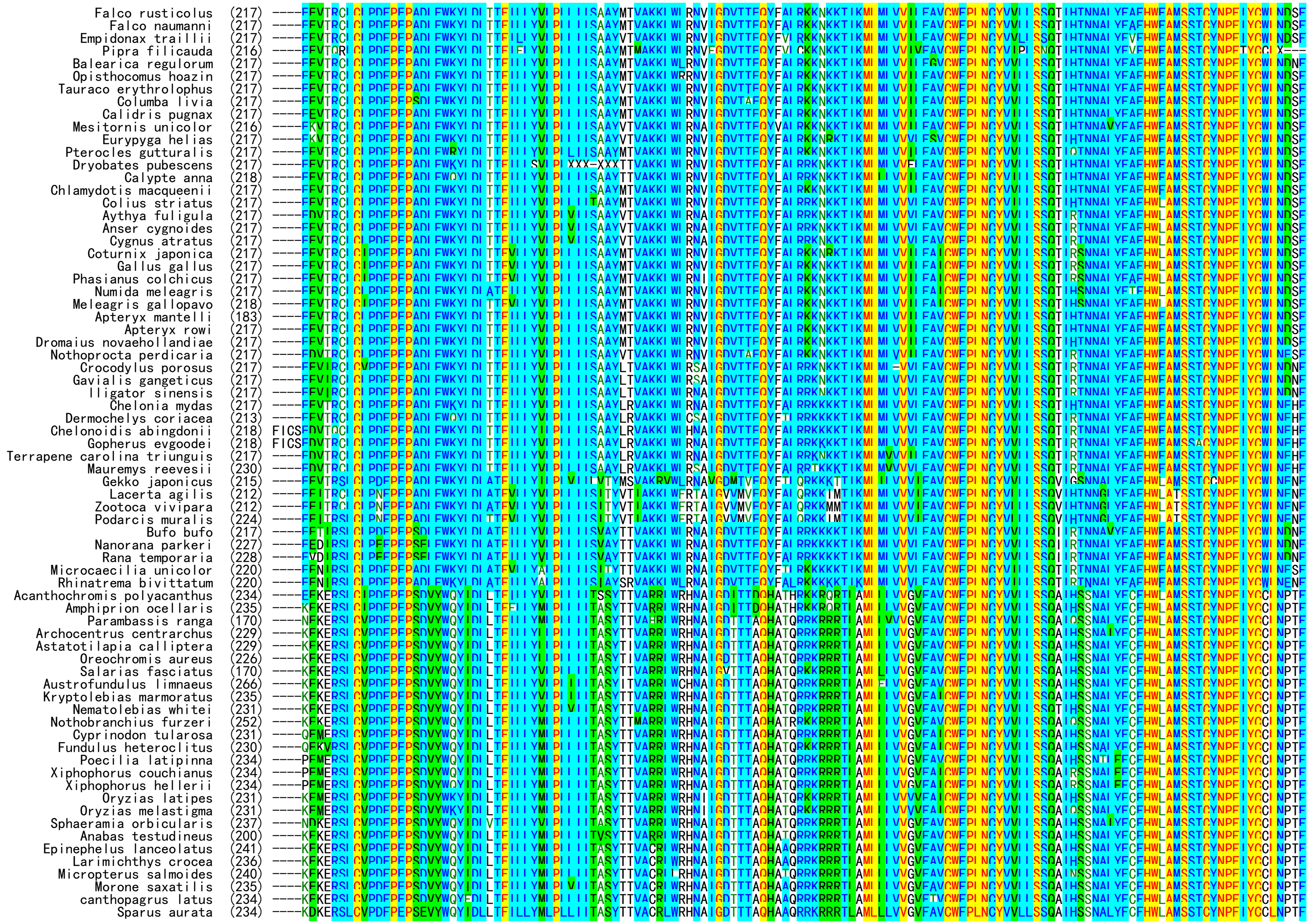
Falco rusticolus (217)
Falco naumanni (217)
Empidonax traillii (217)
Pipra filicauda (216)
Balearica regulorum (217)
Opisthocomus hoazin (217)
Tauraco erythrolophus (217)
Columba livia (217)
Calidris pugnax (217)
Mesitornis unicolor (216)
Eurypyga helias (217)
Pterocles gutturalis (217)
Dryobates pubescens (217)
Calypte anna (218)
Chlamydotis macqueenii (217)
Colius striatus (217)
Aythya fuligula (217)
Anser cygnoides (217)
Cygnus atratus (217)
Coturnix japonica (217)
Gallus gallus (217)
Phasianus colchicus (217)
Numida meleagris (217)
Meleagris gallopavo (218)
Apteryx mantelli (183)
Apteryx rowi (217)
Dromaius novaehollandiae (217)
Nothoprocta perdicaria (217)
Crocodylus porosus (217)
Gavialis gangeticus (217)
lligator sinensis (217)
Chelonia mydas (217)
Dermochelys coriacea (213)
Chelonoidis abingdonii (218)
Gopherus evgoodei (218)
Terrapene carolina triunguis (217)
Mauremys reevesii (230)
Gekko japonicus (215)
Lacerta agilis (212)
Zootoca vivipara (212)
Podarcis muralis (224)
Bufo bufo (217)
Nanorana parkeri (227)
Rana temporaria (228)
Microcaecilia unicolor (220)
Rhinatrema bivittatum (220)
Acanthochromis polyacanthus (234)
Amphiprion ocellaris (235)
Parambassis ranga (170)
Archocentrus centrarchus (229)
Astatotilapia calliptera (229)
Oreochromis aureus (226)
Salarias fasciatus (170)
Austrofundulus limnaeus (266)
Kryptolebias marmoratus (235)
Nematolebias whitei (231)
Nothobranchius furzeri (252)
Cyprinodon tularosa (231)
Fundulus heteroclitus (230)
Poecilia latipinna (234)
Xiphophorus couchianus (234)
Xiphophorus hellerii (234)
Oryzias latipes (231)
Oryzias melastigma (231)
Sphaeramia orbicularis (237)
Anabas testudineus (200)
Epinephelus lanceolatus (241)
Larimichthys crocea (236)
Micropterus salmoides (240)
Morone saxatilis (235)
canthopagrus latus (234)
Sparus aurata (234)

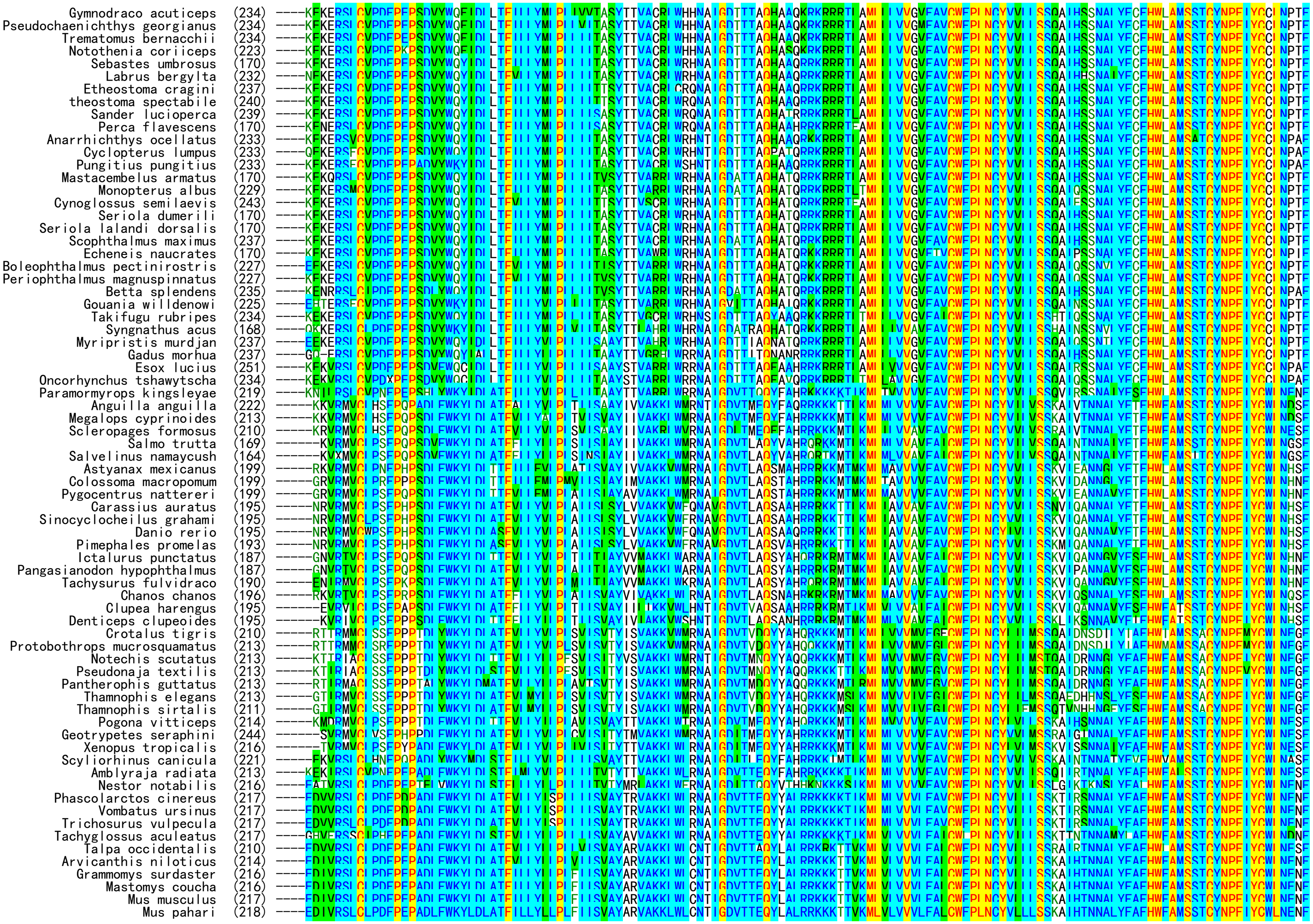
Gymnodraco acuticeps (234)
Pseudochaenichthys georgianus (234)
Trematomus bernacchii (234)
Notothenia coriiceps (223)
Sebastes umbrosus (170)
Labrus bergylta (232)
Etheostoma cragini (237)
theostoma spectabile (240)
Sander lucioperca (239)
Perca flavescens (170)
Anarrhichthys ocellatus (233)
Cyclopterus lumpus (233)
Pungitius pungitius (233)
Mastacembelus armatus (170)
Monopterus albus (229)
Cynoglossus semilaevis (243)
Seriola dumerili (170)
Seriola lalandi dorsalis (170)
Scophthalmus maximus (237)
Echeneis naucrates (170)
Boleophthalmus pectinirostris (227)
Periophthalmus magnuspinnatus (227)
Betta splendens (235)
Gouania willdenowi (225)
Takifugu rubripes (234)
Syngnathus acus (168)
Myripristis murdjan (237)
Gadus morhua (237)
Esox lucius (251)
Oncorhynchus tshawytscha (234)
Paramormyrops kingsleyae (219)
Anguilla anguilla (222)
Megalops cyprinoides (213)
Scleropages formosus (210)
Salmo trutta (169)
Salvelinus namaycush (164)
Astyanax mexicanus (199)
Colossoma macropomum (199)
Pygocentrus nattereri (199)
Carassius auratus (195)
Sinocyclocheilus grahami (195)
Danio rerio (195)
Pimephales promelas (193)
Ictalurus punctatus (187)
Pangasianodon hypophthalmus (187)
Tachysurus fulvidraco (190)
Chanos chanos (196)
Clupea harengus (195)
Denticeps clupeoides (195)
Crotalus tigris (210)
Protobothrops mucrosquamatus (213)
Notechis scutatus (213)
Pseudonaja textilis (213)
Pantherophis guttatus (213)
Thamnophis elegans (213)
Thamnophis sirtalis (211)
Pogona vitticeps (214)
Geotrypetes seraphini (244)
Xenopus tropicalis (216)
Scyliorhinus canicula (221)
Amblyraja radiata (213)
Nestor notabilis (216)
Phascolarctos cinereus (217)
Vombatus ursinus (217)
Trichosurus vulpecula (217)
Tachyglossus aculeatus (217)
Talpa occidentalis (210)
Arvicanthis niloticus (214)
Grammomys surdaster (216)
Mastomys coucha (216)
Mus musculus (217)
Mus pahari (218)

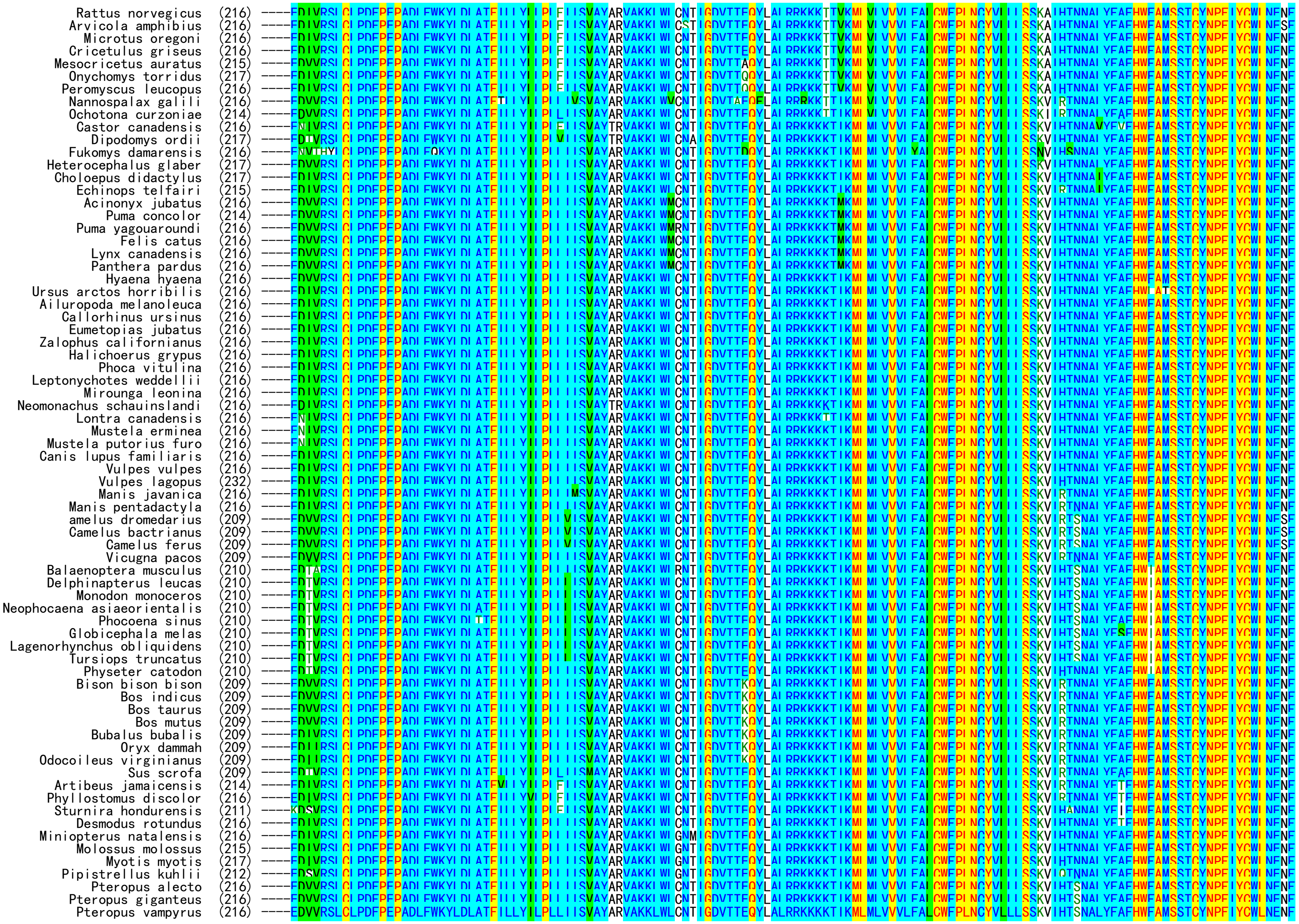

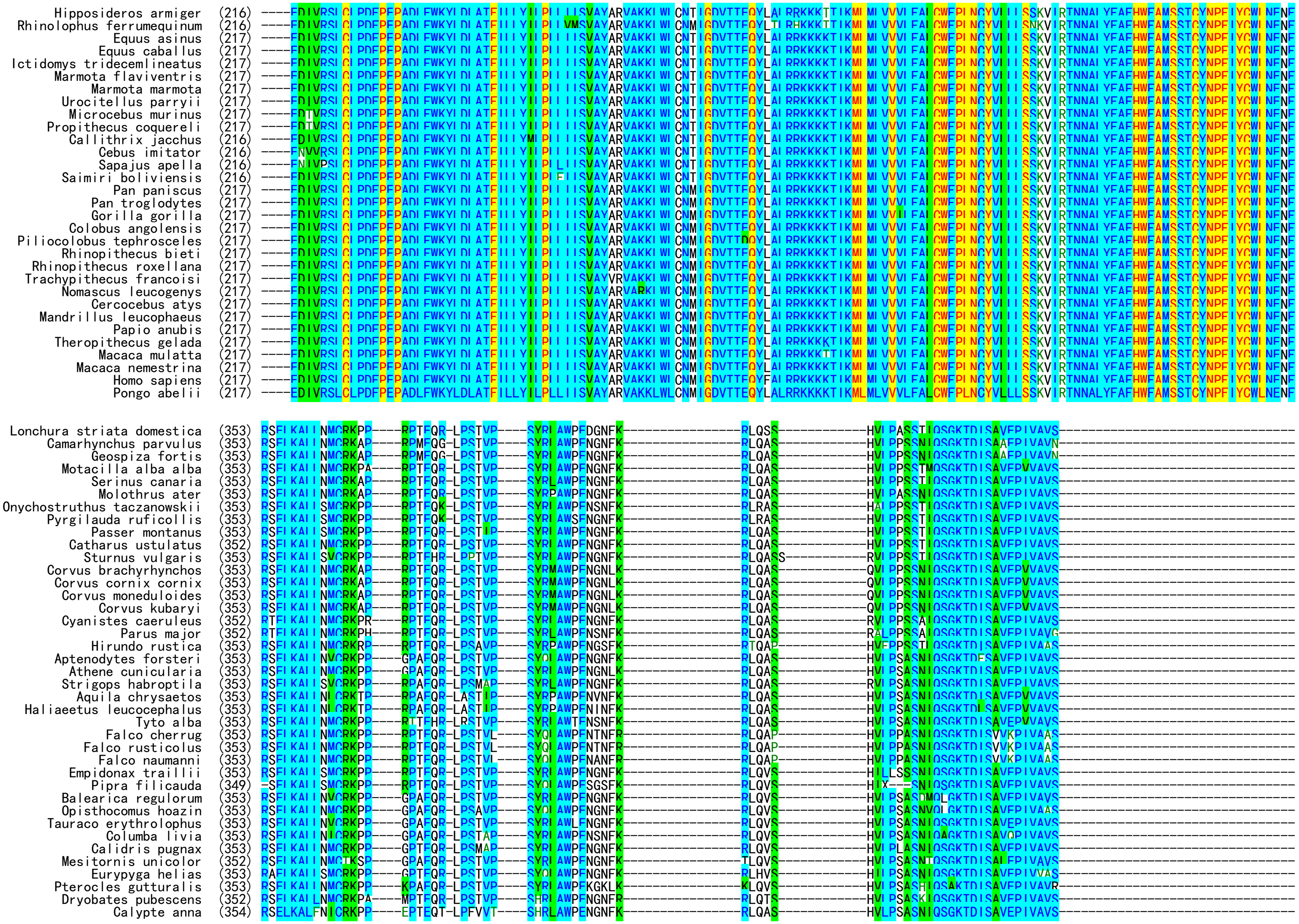
Hipposideros armiger (216)
Rhinolophus ferrumequinum (216)
Equus asinus (217)
Equus caballus (217)
Ictidomys tridecemlineatus (217)
Marmota flaviventris (217)
Marmota marmota (217)
Urocitellus parryii (217)
Microcebus murinus (217)
Propithecus coquereli (217)
Callithrix jacchus (216)
Cebus imitator (216)
Sapajus apella (216)
Saimiri boliviensis (216)
Pan paniscus (217)
Pan troglodytes (217)
Gorilla gorilla (217)
Colobus angolensis (217)
Piliocolobus tephrosceles (217)
Rhinopithecus bieti (217)
Rhinopithecus roxellana (217)
Trachypithecus francoisi (217)
Nomascus leucogenys (217)
Cercocebus atys (217)
Mandrillus leucophaeus (217)
Papio anubis (217)
Theropithecus gelada (217)
Macaca mulatta (217)
Macaca nemestrina (217)
Homo sapiens (217)
Pongo abelii (217)
Lonchura striata domestica (353)
Camarhynchus parvulus (353)
Geospiza fortis (353)
Motacilla alba alba (353)
Serinus canaria (353)
Molothrus ater (353)
Onychostruthus taczanowskii (353)
Pyrgilauda ruficollis (353)
Passer montanus (353)
Catharus ustulatus (352)
Sturnus vulgaris (353)
Corvus brachyrhynchos (353)
Corvus cornix cornix (353)
Corvus moneduloides (353)
Corvus kubaryi (353)
Cyanistes caeruleus (352)
Parus major (352)
Hirundo rustica (353)
Aptenodytes forsteri (353)
Athene cunicularia (353)
Strigops habroptila (353)
Aquila chrysaetos (353)
Haliaeetus leucocephalus (353)
Tyto alba (353)
Falco cherrug (353)
Falco rusticolus (353)
Falco naumanni (353)
Empidonax traillii (353)
Pipra filicauda (349)
Balearica regulorum (353)
Opisthocomus hoazin (353)
Tauraco erythrolophus (353)
Columba livia (353)
Calidris pugnax (353)
Mesitornis unicolor (352)
Eurypyga helias (353)
Pterocles gutturalis (353)
Dryobates pubescens (352)
Calypte anna (354)

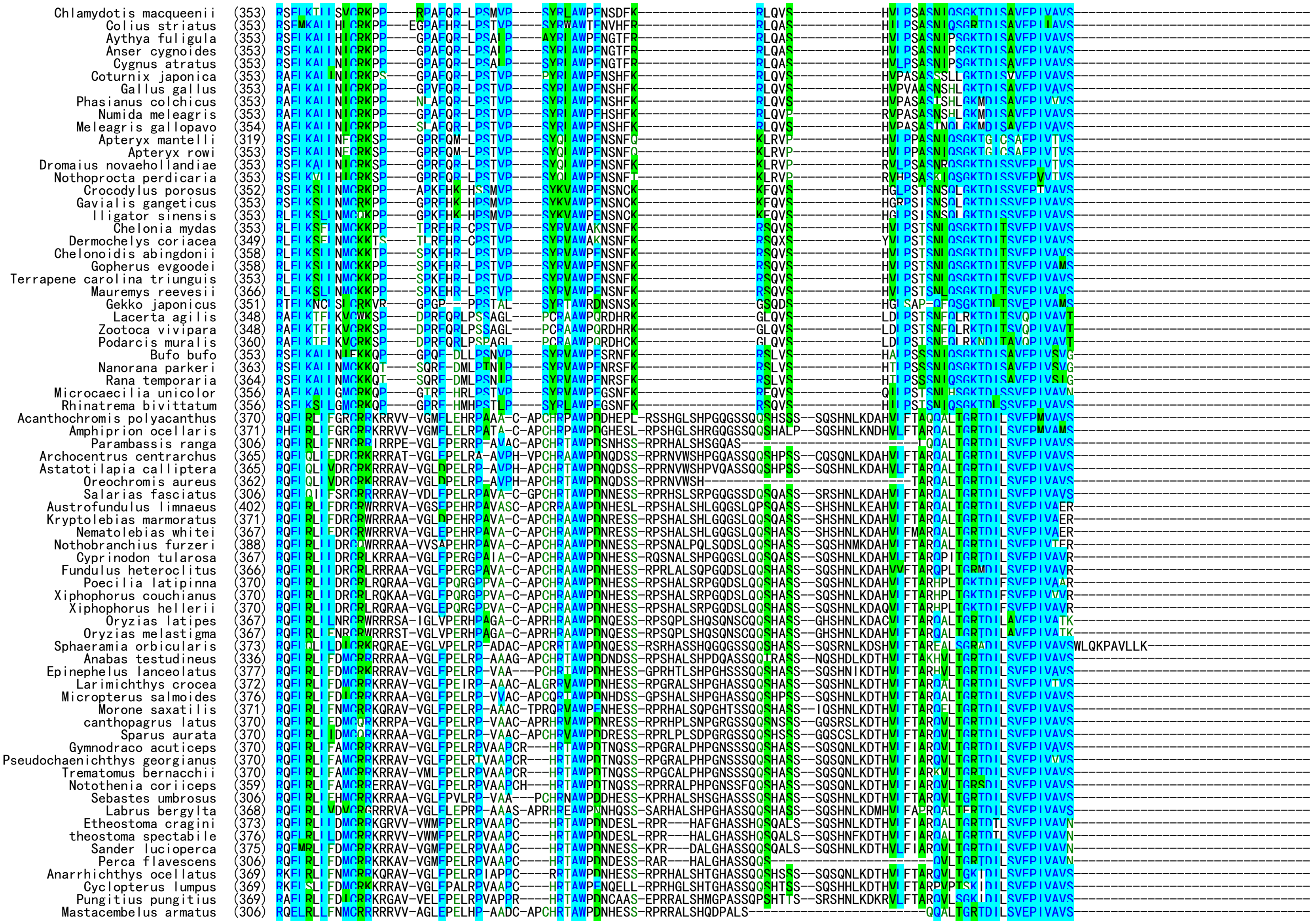

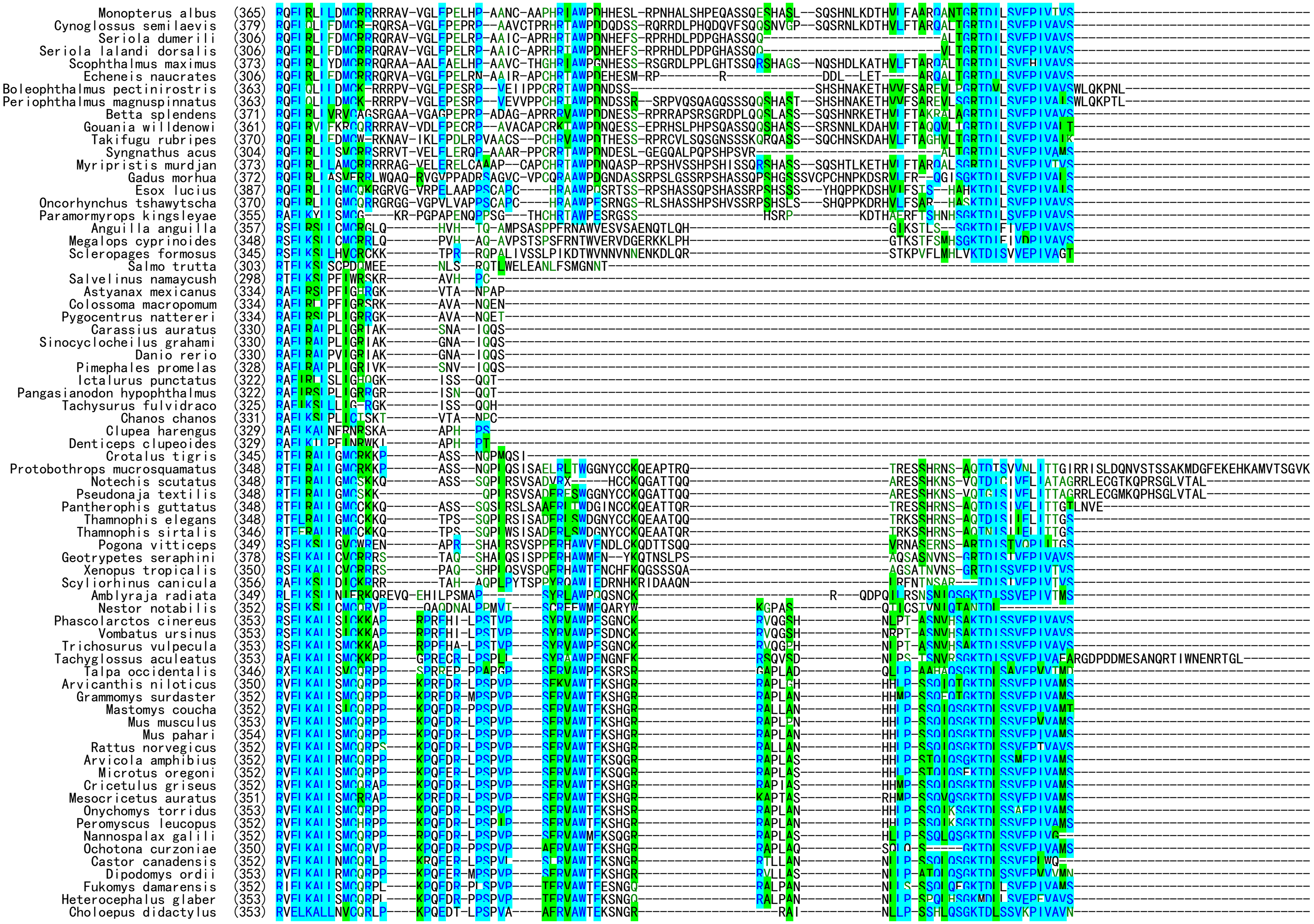
Monopterus albus (365)
Cynoglossus semilaevis (379)
Seriola dumerili (306)
Seriola lalandi dorsalis (306)
Scophthalmus maximus (373)
Echeneis naucrates (306)
Boleophthalmus pectinirostris (363)
Periophthalmus magnuspinnatus (363)
Betta splendens (371)
Gouania willdenowi (361)
Takifugu rubripes (370)
Syngnathus acus (304)
Myripristis murdjan (373)
Gadus morhua (372)
Esox lucius (387)
Oncorhynchus tshawytscha (370)
Paramormyrops kingsleyae (355)
Anguilla anguilla (357)
Megalops cyprinoides (348)
Scleropages formosus (345)
Salmo trutta (303)
Salvelinus namaycush (298)
Astyanax mexicanus (334)
Colossoma macropomum (334)
Pygocentrus nattereri (334)
Carassius auratus (330)
Sinocyclocheilus grahami (330)
Danio rerio (330)
Pimephales promelas (328)
Ictalurus punctatus (322)
Pangasianodon hypophthalmus (322)
Tachysurus fulvidraco (325)
Chanos chanos (331)
Clupea harengus (329)
Denticeps clupeoides (329)
Crotalus tigris (345)
Protobothrops mucrosquamatus (348)
Notechis scutatus (348)
Pseudonaja textilis (348)
Pantherophis guttatus (348)
Thamnophis elegans (348)
Thamnophis sirtalis (346)
Pogona vitticeps (349)
Geotrypetes seraphini (378)
Xenopus tropicalis (350)
Scyliorhinus canicula (356)
Amblyraja radiata (349)
Nestor notabilis (352)
Phascolarctos cinereus (353)
Vombatus ursinus (353)
Trichosurus vulpecula (353)
Tachyglossus aculeatus (353)
Talpa occidentalis (346)
Arvicanthis niloticus (350)
Grammomys surdaster (352)
Mastomys coucha (352)
Mus musculus (353)
Mus pahari (354)
Rattus norvegicus (352)
Arvicola amphibius (352)
Microtus oregoni (352)
Cricetulus griseus (352)
Mesocricetus auratus (351)
Onychomys torridus (353)
Peromyscus leucopus (352)
Nannospalax galili (352)
Ochotona curzoniae (350)
Castor canadensis (352)
Dipodomys ordii (353)
Fukomys damarensis (352)
Heterocephalus glaber (353)
Choloepus didactylus (353)

```
Echinops telfairi               (351) RVELKALLSMCQRLP----KAQEDR-HPSPVP----SFRVAWTEKSNGR-----------------RVPLVN-----------NLPP-SSHLQSGKTDLSSVEPIVTVS---------------------------------
Acinonyx jubatus                (352) RVELKALLSMCQRLP----KPQEDR-PPSPVP----SFRVAWTEKSSGR-----------------RALLAN-----------NLLP-SSQLQSGKTDLSSVEPIVAVS---------------------------------
Puma concolor                   (350) RVELKALLSMCQRLP----KPQEER-PPSPVP----SFRVAWTEKSSGR-----------------RALLAN-----------SLLP-SSQLQSGKTDLSSVEPIVAVS---------------------------------
Puma yagouaroundi               (352) RVELKALLSMCQRLP----KPQEER-PPSPVP----SFRVAWTEKSSGR-----------------RALLAS-----------NFLP-SSQLQSGKTDLSSVEPIVAVS---------------------------------
Felis catus                     (352) RVELKALLSMCQRLP----KPQEER-PPSPVP----SFRVAWTEKSSGR-----------------RAPLAN-----------NLLP-SSQLQSGKTDLSSVEPIVAVS---------------------------------
Lynx canadensis                 (352) RVELKALLSMCQRLP----KPQEER-PPSPVP----SFRVAWTEKSSGR-----------------RAPLAN-----------NLLP-SSQLQSGKTDLSSVEPIVAVS---------------------------------
Panthera pardus                 (352) RVELKALLSMCQRLP----KPQEER-PPSPVP----SFRVAWTEKSSGR-----------------RAPLAN-----------NLLP-SSQLQSGKTDLSSVEPIVAVS---------------------------------
Hyaena hyaena                   (352) RVELKALLSMCQRLP----KPQEER-PPSPIP----SFRVAWTEKTSGR-----------------RAPLAN-----------NLLP-SSQLQSGKTDLSSVEPIVAVS---------------------------------
Ursus arctos horribilis         (352) RVELKALLSMCQRLP----KPQEER-PPSPVP----SFRVAWTEKSSGR-----------------RVPLAS-----------NLLP-SSQLQSGKTDLSSVEPTVAMS---------------------------------
Ailuropoda melanoleuca          (352) RVELKALLSMCQRLP----KPQEER-PPSPVP----SFRVAWTE-SGGR-----------------RAPLAN-----------NLLP-SSQLQSGKTDLSSVEPIVAMS---------------------------------
Callorhinus ursinus             (352) RVELKALLSMCQRLP----KPQEER-PPSPVP----SFRVAWTEKSNGR-----------------RAPLAN-----------NLLP-SSQLQSGKTDLSSVEPIVAVS---------------------------------
Eumetopias jubatus              (352) RVELKALLSMCQRLP----KPQEER-PPSPVP----SFRVAWTEKSNGR-----------------RAPLAN-----------NLLP-SSQLQSGKTDLSSVEPIVAVS---------------------------------
Zalophus californianus          (352) RVELKALLSMCQRLP----KPQEER-PPSPVP----SFRVAWTEKSNGR-----------------RAPLAN-----------NLLP-SSQLQSGKTDLSSVEPIVAVS---------------------------------
Halichoerus grypus              (352) RVELKALLSMCQRLP----KPQEER-PPSPVP----SFRVAWTEKSNGR-----------------RAPLAN-----------NLLA-SSQLQPGKTDLSSVEPIVAVS---------------------------------
Phoca vitulina                  (352) RVELKALLSMCQRLP----KPQEER-PPSPVP----SFRVAWTEKSNGR-----------------RAPLAN-----------NLLA-SSQLQPGKTDLSSVEPIVAVS---------------------------------
Leptonychotes weddellii         (352) RVELKALLSMCQRLP----KPQEGR-PPSPVP----SFRVAWTEKSNGR-----------------RAPLAN-----------NLLP-SSQLQSGKTDLSSVEPIVAVS---------------------------------
Mirounga leonina                (352) RVELKALLSMCQRLP----KPQEGR-PPSPVP----SFRVAWTEKSNGR-----------------RGPLAN-----------NLLP-SSQLQSGKTDLSSVEPIVAVS---------------------------------
Neomonachus schauinslandi       (352) RVELKALLSMCQRLP----KPQEER-PPSPVP----SFRVAWTEKSNGR-----------------RAPLAN-----------NLLP-SSQLQSGKTDLSSVEPIVAVS---------------------------------
Lontra canadensis               (352) RVELKALLSMCQRLP----KAQEER-PPSPVA----SFRVAWTEKSDGR-----------------RAPLAS-----------NLLP-SSQLQSGKTDLSAVEPIVAVS---------------------------------
Mustela erminea                 (352) RVELKALLSMCQRLP----KPQEER-PPSPVA----SFRVAWTEKSDGR-----------------RAPLPN-----------NFLP-SSQLQSGKTDLSAVEPIVAVS---------------------------------
Mustela putorius furo           (352) RVELKALLSMCQRLP----KPQEER-PPSPVA----SFRVAWTEKSDGR-----------------RAPLAN-----------NFLP-SSQLQSGKTDLSAVEPIVAVS---------------------------------
Canis lupus familiaris          (352) RIELKALLSMCQRLP----KPQEER-PPSPVP----SFRVAWTEKSNGR-----------------RVPPAN-----------NLLL-SSHLHSGKTDLSSVEPIVAMS---------------------------------
Vulpes vulpes                   (352) RIELKALLSMCQRLP----KPQEER-PPSPVP----SFRVAWTEKNNGR-----------------RVPPAN-----------NLLL-SSHLQSGKTDLSSVETVVAMS---------------------------------
Vulpes lagopus                  (368) RIELKALLSMCQRLP----KPQEER-PPSPVP----SFRVAWTEKSNGR-----------------RVPPAN-----------NLLL-SSHFQSGKTDLSSVETVVAMS---------------------------------
Manis javanica                  (352) RVELKALLGMCQRPA----KPQEEQ-PPSPVP----SFRVAWTEESNGR-----------------RAPPAN-----------HLLP-SSQVQSGKTDLSSREPTVAMS---------------------------------
Manis pentadactyla              (352) RVEFKALLGMCQRPT----KPQEER-PPSPVP----SFRVAWTEKSNGR-----------------RAPLAN-----------HLLP-SSQVLSGKTDLSSVEPTVAMS---------------------------------
amelus dromedarius              (345) RVELKALLSMCRRPP----KPQEAQ-PPSPAPSFRVAFRVAWTEKGSGR-----------------RAPLAN-----------ALP--SSQLQSGKTDLSSVEPIVAVS---------------------------------
Camelus bactrianus              (345) RVELKALLSMCRRPP----KPQEAQ-PPSPAPSFRVAFRVAWTEKGSGR-----------------RAPLAN-----------ALP--SSQLQSGKTDLLSVEPIVAVS---------------------------------
Camelus ferus                   (345) RVELKALLSMCRRPP----KPQEAQ-PPSPAPSFRVAFRVAWTEKGSGR-----------------RAPLAN-----------ALP--SSQLQSGKTDLLSVEPIVAVS---------------------------------
Vicugna pacos                   (345) RVELKALLSMCRRPP----KPQEAQ-PPSPAPSFRLAFRVAWTEKGSGR-----------------RAPLAN-----------ALP--SSQLQSGKTDLSSVEPIVAMS---------------------------------
Balaenoptera musculus           (346) RVELKALLRMCQRSP----KPQEER-PPSPVP----SFRVAWTEKSSGL-----------------RAPLAN-----------SLLP-SSQLQSGKTDLSSVEPMVAF----------------------------------
Delphinapterus leucas           (346) RVALKALLRVCQRSP----KPREER-PPSPVP----SFRVAWTEKSSGR-----------------SAPLAH-----------SLLP-PSQLQSGKTDLSSVEPVVAVT---------------------------------
Monodon monoceros               (346) RVALKALLRVCQRSP----KPREER-PPSPVP----SFRVAWTEKSSGR-----------------SAPLAH-----------SLLP-PSQLQSGKTDLSSVEPVVAVT---------------------------------
Neophocaena asiaeorientalis     (346) RVALKALLRVCQRSP----KPREER-PPSPVP----SFRVAWTEKSSGR-----------------SAPLAH-----------SLLP-PSQLQSGKTDLSSVEPVVAVT---------------------------------
Phocoena sinus                  (346) RVALKALLRVCQRSP----KPREER-PPEPVP----SFRVAWTEKSSGR-----------------SAPLAH-----------SLLP-PSQLQSGKTDLSSVEPVVAVT---------------------------------
Globicephala melas              (346) RVALKALLRVCQRSP----KPREER-PPSLVP----SFRVAWTEKSSGR-----------------SAPLAN-----------SLLP-SSQLQSGKTDLLSVEPVVAVT---------------------------------
Lagenorhynchus obliquidens      (346) RVALKALLRVCQRSP----KPREER-PPSPVP----SFRVAWTEKSSGR-----------------SAPLAN-----------SLLP-SSQLQSGKTDLSSVEPVVAVT---------------------------------
Tursiops truncatus              (346) RVALKALLRVCQRSP----KPREER-PPSPIP----SFRVAWTEKSSGR-----------------SAPLAN-----------SLLP-SSQLQSGKTDLSSMEPVVAVT---------------------------------
Physeter catodon                (346) RVELKALLRVCQRSP----KPQEER-PPSPVP----SFRVAWTEKSSGR-----------------RAPLAN-----------ILLP-SSQLQSGKTDLSSVEPIVAVT---------------------------------
Bison bison bison               (345) RVELKALLHMCQRSP----KPQEGQ-PPSPVP----SLRVAWAEKSSGR-----------------RASPAN-----------SLLP-SSQVQSGRTDLASVEPIVTMT---------------------------------
Bos indicus                     (345) RVELKALLHXCQRSP----KPQEGQ-PXSPVP----SLRVAWAEKSSGR-----------------RASPAN-----------SLLP-SSQVQSGRTDLASVEPIVTMT---------------------------------
Bos taurus                      (345) RVELKALLHMCQRSP----KPQEGQ-PPSPVP----SLRVAWAEKSSGR-----------------RASPAN-----------SLLP-SSQVQSGRTDLASVEPIVTMT---------------------------------
Bos mutus                       (345) RVELKALLHMCQRSP----KPQEGQ-PPSHVP----SLRVAWAEKSSGC-----------------RASPAN-----------SLLP-SSQVQSGRTDLASVEPIVTMT---------------------------------
Bubalus bubalis                 (345) RVELKALLRMCQRSP----KPQEGQ-PPSPVP----SLRVAWAEKSSSR-----------------RALPAN-----------SLLP-SSQVQSGRTDLASVEPIVTMT---------------------------------
Oryx dammah                     (345) RVELKALLRMCQRSP----KPQEGQ-PPSPVP----SLRVAWAEKSSGR-----------------RAPPAS-----------SLLP-SSQVQSGRTDLASVEPTVTMT---------------------------------
Odocoileus virginianus          (345) RVELKALLRMCQRSP----KPHEGQ-PPSPVP----SLRVAWAERSSGR-----------------RAPPAN-----------SLLP-SSQLQSGRTDLASVEPIMTMT---------------------------------
Sus scrofa                      (345) RVELKALLSMCQRPP----KPPEER-PPSPVP----SFRMAWTEKSSGR-----------------RAPLAN-----------HLPP-SSQLQSGKTDLSSVEPIVAMT---------------------------------
Artibeus jamaicensis            (350) RVELKALLSMCQRLP----KPQEEQ-PPSPVP----SFRVAWTEKGNGW-----------------RSPVAN-----------SLLP-SPQLQSGKTYLSSFEPVVVMS---------------------------------
Phyllostomus discolor           (352) RVELKALLSMCQRLP----KPQEEQ-PPSPVP----SFRVAWTEKGNGW-----------------RSPVAN-----------SLLP-SPQLQSGKTYLSSVEPIVVMS---------------------------------
Sturnira hondurensis            (347) RVELKALLSMCQRLP----KPQEEQ-PPSPVP----SFRVAWTEKGNGW-----------------RSPVAN-----------SLLP-SPQLQSGKTYLSSAEPIVVMS---------------------------------
Desmodus rotundus               (352) RVELKALLSMCQRLP----KPQEEQ-PPSPVP----SFRVAWTEKGNGW-----------------RSPVVN-----------GLL--SPQLQSGKTDLSSVEPIVVMS---------------------------------
Miniopterus natalensis          (352) RVELKALLSMCQRPP----KPQEEQ-PPSPVP----SFRVAWTEKGNGR-----------------RAPLAN-----------NLLP-SPQLQSGKTDLLSVEPIVAMS---------------------------------
Molossus molossus               (351) RVELKALLSMFQRPP----KPQEEQ-PPSPVP----SFRVAWTEKSNGR-----------------RAPLAN-----------NLLP-SPQLQSGKTDLLSVEPIVAVS---------------------------------
Myotis myotis                   (353) RVELKALLSMCQRPP----KPQEER-PPSPVP----SFRMAWTEKGKGR-----------------RATLAN-----------NLLP-SPQLQSGKTDLLSVEPIVAMS---------------------------------
Pipistrellus kuhlii             (348) RVELKALLSMCQRPP----KPQEER-PPSPVP----SFRMAWTEKGKGR-----------------RATLAN-----------NLLP-SPQLQSGKTDLLSVEPIVAMS---------------------------------
Pteropus alecto                 (352) RVELKALLSMCQRPP----KPQEER-PPSPVP----SFRMAWTEKDNGCS----------------RAPLTN-----------NLLP-SAQLQSGKTDLSSVEPIVVMS---------------------------------
Pteropus giganteus              (352) RVELKALLSMCQRPP----KPQEER-PPSPVP----SFRMAWTEKDNGCS----------------RAPLTN-----------NLLP-SAQLQSGKTDLSSVEPIVVMS---------------------------------
Pteropus vampyrus               (352) RVELKALLSMCQRPP----KPQEER-PPSPVP----SFRMAWTEKDNGCS----------------RAPLTN-----------NLLP-SAQLQSGKTDLSSVEPIVVMS---------------------------------
Hipposideros armiger            (352) RVELKALLSMCQRPP----KPQEER-PSSPVP----SFRVAWTEKGHGR-----------------RAPLAN-----------PLLP-SSQLQSGKTDLSSVEPIVAMS---------------------------------
Rhinolophus ferrumequinum       (352) RVELKALLRVFQRPP----KPPEER-PPSPVP----SFRVAWTEKGHGR-----------------RAPLAN-----------HLLS-SSQLQSGKTDLSSVEPIVAMS---------------------------------
Equus asinus                    (353) RIELKALLSMCQRPP----KPQEER-PPSPVP----SFRVAWTEKSKSR-----------------RAPLAN-----------NLLP-SSQLQSGKTDLSSMEPIMAMS---------------------------------
Equus caballus                  (353) RIELKALLSMCQRPP----KPQEER-PPSPVP----SFRVAWTEKSKSR-----------------RAPPAN-----------NLLP-SSQLQSGKTDLSSMEPIMAMS---------------------------------
Ictidomys tridecemlineatus      (353) RVELKALLSMCQRPP----KPQEDR-PPSPVA----SFRVAWTEKSNGG-----------------RAPLAN-----------NLLP-SSQLQSGKTDLSSVEPIVAMS---------------------------------
Marmota flaviventris            (353) RVELKALLSMCQRPP----KPQEDR-PPSPVA----SFRVAWTEKSNGR-----------------RAPLPN-----------NLLP-SSQLQSGKTDLSSVEPIVAMS---------------------------------
Marmota marmota                 (353) RVELKALLSMCQRPP----KPQEDR-PPSPVA----SFRVAWTEKSNGR-----------------RAPLPN-----------NLLP-SSQLQSGKTDLLSVEPIVAMS---------------------------------
Urocitellus parryii             (353) RVELKALLSMCQRPP----KPQEDR-PPSPVA----SFRVAWTEKSNGR-----------------RAPLAN-----------NLLP-SSQLQSGKTDLSSVEPIVAMS---------------------------------
Microcebus murinus              (353) RVELKSLLSMCQRPP----KPQEDR-PPSPVP----SFRVAWTEKSNGRK----------------APPLAN-----------NLLP-SSQLQAGKTDLSSVEPIVAMS---------------------------------
Propithecus coquereli           (353) RVELKSLLSMCQRPP----KPQEDR-PPSPVP----SFRVAWTEKSNGRR----------------APLLAN-----------NLLP-SSQLQSGKTDLSSVEPTVAMS---------------------------------
Callithrix jacchus              (352) RIELKALLNMCQRPP----KPQEER-PPSPVP----SFRVAWTEKNDGQ-----------------RAPLTN-----------NLLP-TSQLQSGKTDLSSVEPIVMMS---------------------------------
Cebus imitator                  (352) RIELKALLNMCQRPP----KPQEER-PPSPVP----SFRVAWTEKSDGQ-----------------RAPLTN-----------NLLP-TSQLQSGKTDLSSVEPIVTMS---------------------------------
Sapajus apella                  (352) RIELKALLNMCQRPP----KPQEER-PPSPVP----SFRVAWTEKSDGQ-----------------RAPLTN-----------NLLP-TSQLQSGKTDLSSVEPIVTMS---------------------------------
Saimiri boliviensis             (352) RTELKALLNMCQRPP----KPREER-LPSPVP----SFRVAWTEKSDGQ-----------------RAPLTS-----------NLLA-TSQLQSGKTDLSSVEPIVTMS---------------------------------
```

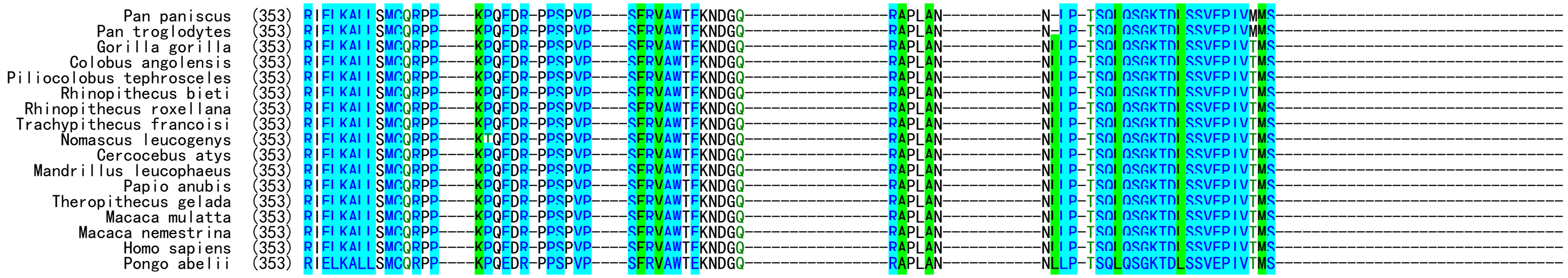


**Fig. S1**. Amino acid sequence alignment of GPR83 orthologs from fish to mammals.

#### Alignment of FAM237A orthologs from fish to mammals

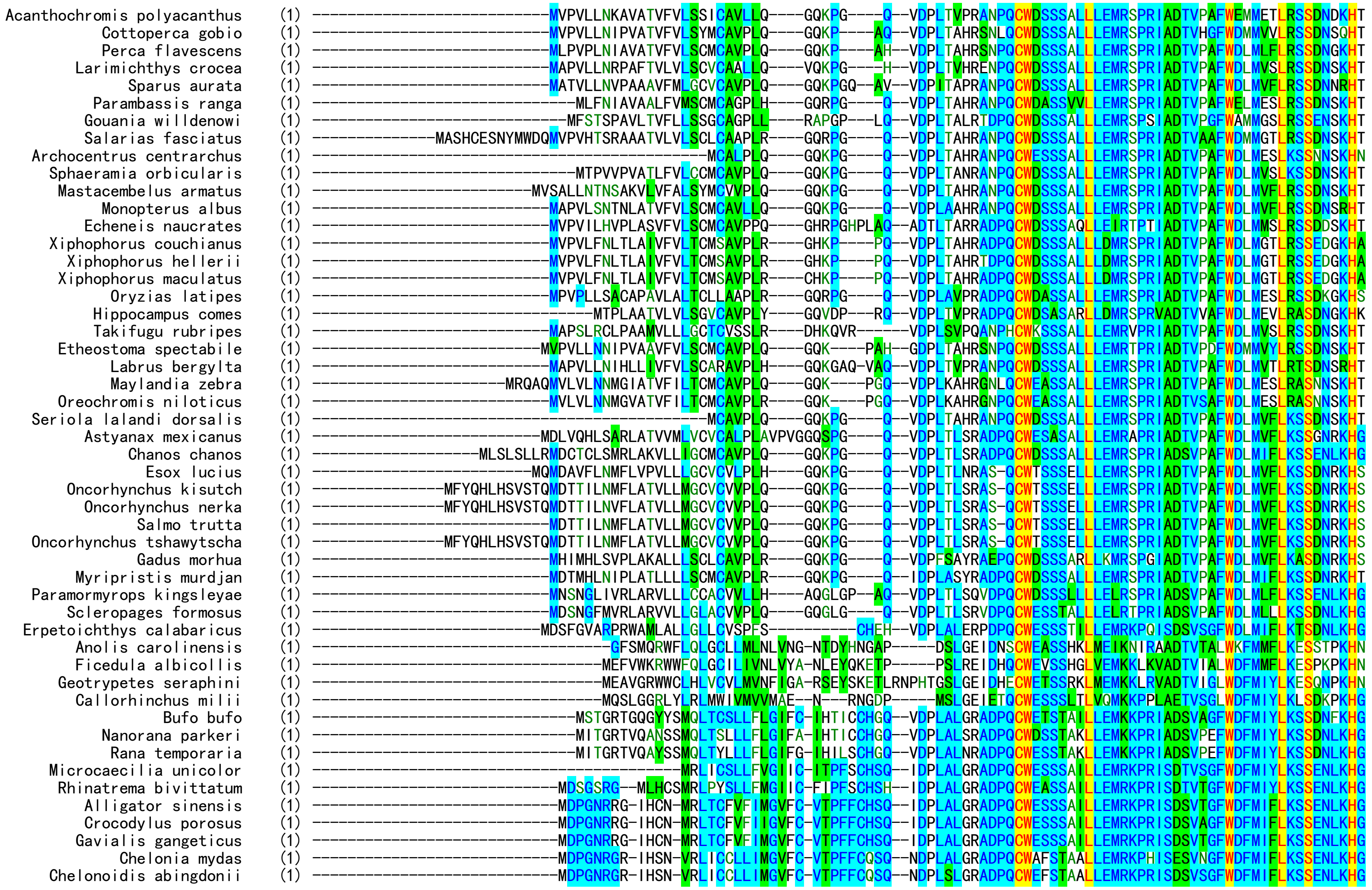

```
              Gopherus evgoodei  (1) ----------------------------MDPGNRGR-IHSN-MRLICSLLIMGVFC-VTPFFCQSQ--NDPLSLGRADPQCWEFSTAALVEMRKPRISDSVSGFWDFMIFLKSSENLKHG
              Mauremys reevesii  (1) ----------------------------MDPGNRGR-IHSN-ARLICSLLIMGVFC-VTPFFCQSQ--NDPLSLGRADPQCWEFSTAALLEMRKPRISDSVSGFWDFMIFLKSSENLKHG
               Chrysemys picta  (1) ----------------------------MDPGNRGR-IHS--VRLICSLLIMGVFC-VTPFFCHSQ--NDPLALGRADPQCWEFSTAALLEMRKPRISDSVSGFWDFMIFLKSSENLKHG
      Trachemys scripta elegans  (1) ----------------------------MDPGNRGR-IHS--VRLICSLLIMGVFC-VTPFFCHSQ--NDPLALGRADPQCWEFSTAALLEMRKPRISDSVSGFWDFMIFLKSSENLKHG
  Terrapene carolina triunguis  (1) ---------------------MESFSDTMDPGNRGR-IHS--VRLICSLLIMGVFC-VTPFFCQSQ--NDPLALGRADPQCWEFSTAALLEMRKPRISDSVSGFWDFMIFLKSSENLKHG
            Pelodiscus sinensis  (1) ----------------------------MDPGNRGR-IHCN-VRLICSLLVMGAFC-VTPFFCQSQ--NDPLALGRADPRCWEFSSAALLEMRKPRISDSVSGFWDFMIFLKSSENLKHG
             Anas platyrhynchos  (1) ----------------------------MDLGNRGRRIHYN-MRLTCSLLIMGVFC-VTPFLCHSQ--IDPLALGRADPQCWESSSAVLLEMRKPRISDSVSGFWDFMIFLKSSENLKHG
              Oxyura jamaicensis  (1) ----------------------------MDLGKRGRRIHYNNMRLTCSLLIMGVFC-VTPFLCHSQ--IDPLALGRADPQCWESSSAVLLEMRKPRISDSVSGFWDFMIFLKSSENLKHG
                Aythya fuligula  (1) ----------------------------MDLGNRGRRIHYN-MRLTCSLLIMGVFC-VTPFLCHSQ--IDPLALGRADPQCWESSSAVLLEMRKPRISDSVSGFWDFMIFLKSSENLKHG
                Cygnus atratus  (1) ----------------------------MDLGNRGRRIHYN-MRLTCSLLIMGVFC-VTPFLCHSQ--IDPLALGRADPQCWESSSAVLLEMRKPRISDSVSGFWDFMIFLKSSENLKHG
                   Cygnus olor  (1) ----------------------------MDLGNRGRRIHYN-MRLTCSLLIMGVFC-VTPFLCHSQ--IDPLALGRADPQCWESSSAVLLEMRKPRISDSVSGFWDFMIFLKSSENLKHG
      Anser cygnoides domesticus  (1) ----------------------------MDLGNRGRRIHYN-MRLTCSLLIMGVFC-VTPFLCHSQ--IDPLALGQADPQCWESSSAVLLEMRKPRISDSVSGFWDFMIFLKSSENLKHG
                  Gallus gallus  (1) ----------------------------MDFGNRGR-IHYN-MRLTYSLLVMGVFC-VTPSLCHSQ--IDPLALGRADPQCWESSSAVLLEMRKPRISDSVSGFWDFMIFLKSSENLKHG
            Phasianus colchicus  (1) ----------------------------MDFGNKGR-IHCN-MRLTYSLLLMTVFC-VTPSLCHSQ--IDPLALGRADPQCWESSSAVLLEMRKPRISDSVSGFWDFMIFLKSSENLKHG
              Numida meleagris  (1) ----------------------------MDFGNRGR-IHCS-MRLTYSLLVMGMFC-VTTFLCHSQ--IDPLALGRADPQCWESSSAVLLEMRKPRISDSVSGFWDFMIFLKSSENLKHG
      Antrostomus carolinensis  (1) ----------------------------MDLQSRRR-IHCN-MRLTCSLLIMGVFC-VTPVLCYSQ--IDPLALGRADPQCWESSSAVLLEMRKPRISDSVSGFWDFMIFLKSSENLKHG
           Aptenodytes forsteri  (1) ----------------------------MDLGSRGR-IHCN-MRLTCSLLIMGAFC-VTPFLCHSQ--IDPLVLGRADPQCWESSSAVLLEMRKPRISDSVSGFWDFMIFLKSSENLKHG
             Pygoscelis adeliae  (1) ----------------------------MDLGSRGR-IHCN-MRLTCSLLIMGAFC-VTPFLCHSQ--IDPLVLGRADPQCWESSSAVLLEMRKPRISDSVSGFWDFMIFLKSSENLKHG
                Nipponia nippon  (1) ---------------------------MMDLGSRGR-IHCN-MRLTCSLLIMGVFC-VTPFLCHSQ--IDPLALGRADPQCWESSSAVLLEMRKPRISDSVSGFWDFMIFLKSSENLKHG
          Chlamydotis macqueenii  (1) ----------------------------MDLGSRGR-IHCN-MRLTCSLLILGAFC-VTPFLCHSQ--IDPLALGQADPQCWESSSTVLLEMRKPRISDSVSGFWDFMIFLKSSENLKHG
              Egretta garzetta  (1) ----------------------------MDLGSRGR-SHCN-MRLTCSLLIMGAFC-VTPFLCHNQ--IDPLALGRADPQCWESSSAVLLEMRKPRISDSVSGFWDFMIFLKSSENLKHG
           Pterocles gutturalis  (1) ------------------------------------------MRLTCSLLIMGAFC-VTPFLCHSQ--IDPLALGRADPQCWESSSAVLLEMRKPRISDSVSGFWDFMIFLKSSENLKHG
                 Falco cherrug  (1) ----------------------------MDLGNRGR-IHCN-MRLICSLLIMGAFC-MSPFLCHSQ--TDPLALGRADPQCWESSSAVLLEMRKPRISDSVSGFWDFMIFLKSSENLKHG
              Falco rusticolus  (1) ----------------------------MDLGNRGR-IHCN-MRLICSLLIMGAFC-MSPFLCHSQ--TDPLALGRADPQCWESSSAVLLEMRKPRISDSVSGFWDFMIFLKSSENLKHG
                Falco naumanni  (1) ----------------------------MDLRNRGR-IHCN-MRLICSLLIMGAFC-TSPFLCHSQ--IDPLALGRADPQCWESSSAVLLEMRKPRISDSVSGFWDFMIFLKSSENLKHG
    Aquila chrysaetos chrysaetos  (1) ----------------------------MDLGSRRI-IHCN-MRLTCTLLIMGAFC-MTPFLCYSQ--IDPLALGRADPQCWESSSAVLLEMRKPRISDSVSGFWDFMIFLKSSENLKHG
             Athene cunicularia  (1) ----------------------------MDLGSRGR-IHCS-MRLTCTLLIMGAFC-MTPFLCHSQ--IDPLALGRADPQCWESSSAVLLEMRKPRISDSVSGFWDFMIFLKSSENLKHG
            Leptosomus discolor  (1) ----------------------------MDLGSRGR-IHCN-MRLTCTLLIMGAFC-MTPFLCHSQ--IDPLALGRADPQCWESSSAVLLEMRKPRISDSVSGFWDFMIFLKSSENLKHG
                    Tyto alba  (1) ----------------------------MDLGSRGR-IHCN-MRLTCILLIMGAFC-MTPFLCHSQ--IDPLALGRADPQCWESSSAVLLEMRKPRISDSVSGFWDFMIFLKSSENLKHG
          Chiroxiphia lanceolata  (1) ----------------------------MDLGSRGR-IHCN-MRLTCSLLIMAAFC-MTPFLCHSQ--TDPLALGWADPQCWESSSAVLLEMRKPRISDSVSGFWDFMIFLKSSENLKHG
        Neopelma chrysocephalum  (1) ----------------------------MDLGSRGR-IHCN-MRLTSSLLIMAAFC-MTPFLCHSQ--TDPLALGWADPQCWESSSAVLLEMRKPRISDSVSGFWDFMIFLKSSENLKHG
             Manacus vitellinus  (1) ----------------------------MDLGSRGR-IHCN-MRLTCSLLIMAAFC-MTPFLCHSQ--TDPLALGWADPQCWESSSAVLLEMRKPRISDSVSGFWDFMIFLKSSENLKHG
               Pipra filicauda  (1) ----------------------------MDLGSRGR-IHCN-MRLTCSLLIMAAFC-MTPFLCHSQ--TDPLALGWADPQCWESSSVVLLEMRKPRISDSVSGFWDFMIFLKSSENLKHG
               Corapipo altera  (1) ----------------------------MDLGSRGR-IQCN-MRLTCSLLIMAAFC-MTPFLCHSQ--TDPLALGWADPQCWESSSAVLLEMRKPRISDSVSGFWDFMIFLKSSENLKHG
             Empidonax traillii  (1) ----------------------------MDLGSRGR-IHCN-MRLTCCLLIMAAFC-MTPFLCHSQ--IDPLALGRADPQCWESSSAVLLEMRKPRISDSVSGFWDFMIFLKSSENLKHG
         Tauraco erythrolophus  (1) ------------------------------------------MRLTCSLLIMGAFC-VTPFLCHSQ--IDPLALGRADPQCWESSSAVLLEMRKPRISDSVSGFWDFMIFLKSSENLKHG
               Calidris pugnax  (1) ----------------------------MDPGSRGR-IHCN-MRLTCSLLIMGAFC-VTPSLCHSQ--IDPLALGRADPQCWESSSAVLLEMRKPRISDSVSGFWDFMIFLKSSENLKHG
                 Columba livia  (1) ----------------------------MDLGSRGR-IHCN-MRLTCSLLIMGAFC-VTPFLCHSQ--IDPLALGRADPQCWESSSAVLLEMRKPRISDSVSGFWDFMIFLKSSENLKHG
              Phaethon lepturus  (1) ----------------------------MDLGSRGR-ILCN-MRLTCSLLIMGALC-VTPFLCHSQ--IDPLALGRADPQCWESSSAVLLEMRKPRISDSVSGFWDFMIFLKSSENLKHG
               Eurypyga helias  (1) ----------------------------MDLGSRGR-IHCN-MRLTCSLLIMGAFC-MTPFLCHSQ--TDPLALGRADPQCWESSSAVLLEMRKPRISDSVSGFWDFMIFLKSSENLKHG
              Nestor notabilis  (1) ----------------------------MDLGNRRR-IHCN-MRLTYSLLIMGAFC-MTPFLCHSQ--IDPLALGRADPQCWQSSSAVLLEMRKPHISDSVSGFWDFMIFLKSSENLKHG
            Strigops habroptila  (1) ----------------------------MDLGNRRR-IHCN-MRLTYSLLIMGAFC-MTPFLCHSQ--IDPLALGRADPQCWESSSAVLLEMRKPHISDSVSGFWDFMVFLKSSENLKHG
           Camarhynchus parvulus  (1) ----------------------------MDLGSRGR-IHCT-MRLTRSLLLMAVFC-ATPLLCHSQ--VDPLALGRADPQCWESSSAVLLEMRKPRISDSVSGFWDFMIFLKSSENLKHG
               Geospiza fortis  (1) ----------------------------MDLGSRGR-IHCT-MRLTRSLLLMAVFC-ATPLLCHSQ--VDPLALGRADPQCWESSSAVLLEMRKPRISDSVSGFWDFMIFLKSSENLKHG
                Molothrus ater  (1) ----------------------------MDLGSRGR-IHCT-MRLTCSLLLMAVFC-VTPFLCHSQ--ADPLALGRADPQCWESSSAVLLEMRKPRISDSVSGFWDFMIFLKSSENLKHG
         Zonotrichia albicollis  (1) ----------------------------MHLGSRRR-IPCT-MRLTCSLLLMAVFC-VTPLLCHSQ--VDPLALGRADPQCWESSSAILLEMRKPRISDSVSGFWDFMIFLKSSENLKHG
               Serinus canaria  (1) ----------------------------MDLGGRGR-IHCT-MRLTCSLLLLAVFC-VTPFLCHSQ--VDPLALGRADPQCWESSSAVLLEMRKPRISDSVSGFWDFMIFLKSSENLKHG
     Lonchura striata domestica  (1) ----------------------------MDLGSRGR-IHCT-MRLTCSLLLMAVFC-VTPFLCHSQ--IDPLALGRADPQCWESSSAVLLEMRKPHISDSVSGFWDFMIFLKSSENLKHG
           Taeniopygia guttata  (1) ----------------------------MDLGSRGR-IHCT-MRLTCSLLLMAVFC-VTPFLCHSQ--IDPLALGRADPQCWESSSAVLLEMRKPHISDSVSGFWDFMIFLKSSENLKHG
             Motacilla alba alba  (1) --MDRNTKRLESLGEELHGSIHHMHSDIMDPGSRGR-IHCT-MRLTCSLLLMAVFC-VTPFLCHSQ--IDPLALGRADPQCWESSSAVLLEMRKPRISDSVSGFWDFMIFLKSSENLKHG
   Onychostruthus taczanowskii  (1) ----------------------------MDPGSRGR-IHCT-MRLTCSLLLMAVFC-VTPFLCHSQ--IDPLALGRADPQCWESSSAVLLEMRKPRISDSVSGFWDFMIFLKSSENLKHG
               Passer montanus  (1) ----------------------------MDPGSRGR-IHCT-MRLTCSLLLMAVFC-VTPFLCHSQ--IDPLALGRADPQCWESSSAVLLEMRKPRISDSVSGFWDFMIFLKSSENLKHG
           Pyrgilauda ruficollis  (1) --MHRNTKRLEGVGEELHGSIHYMHSAIMDPGSRGR-IHCT-MRLTCSLLLIAVFC-VTPVLCHSQ--IDPLALGRADPQCWESSSAVLLEMRKPRISDSVSGFWDFMIFLKSSENLKHG
           Corvus cornix cornix  (1) ----------------------------MDLGSRGR-IHCT-MRLTCSLLLMAVFC-VKPFLCHSQ--IDPLALGRADPQCWESSSAVLLEMRKPRISDSVSGFWDFMIFLKSSENLKHG
```

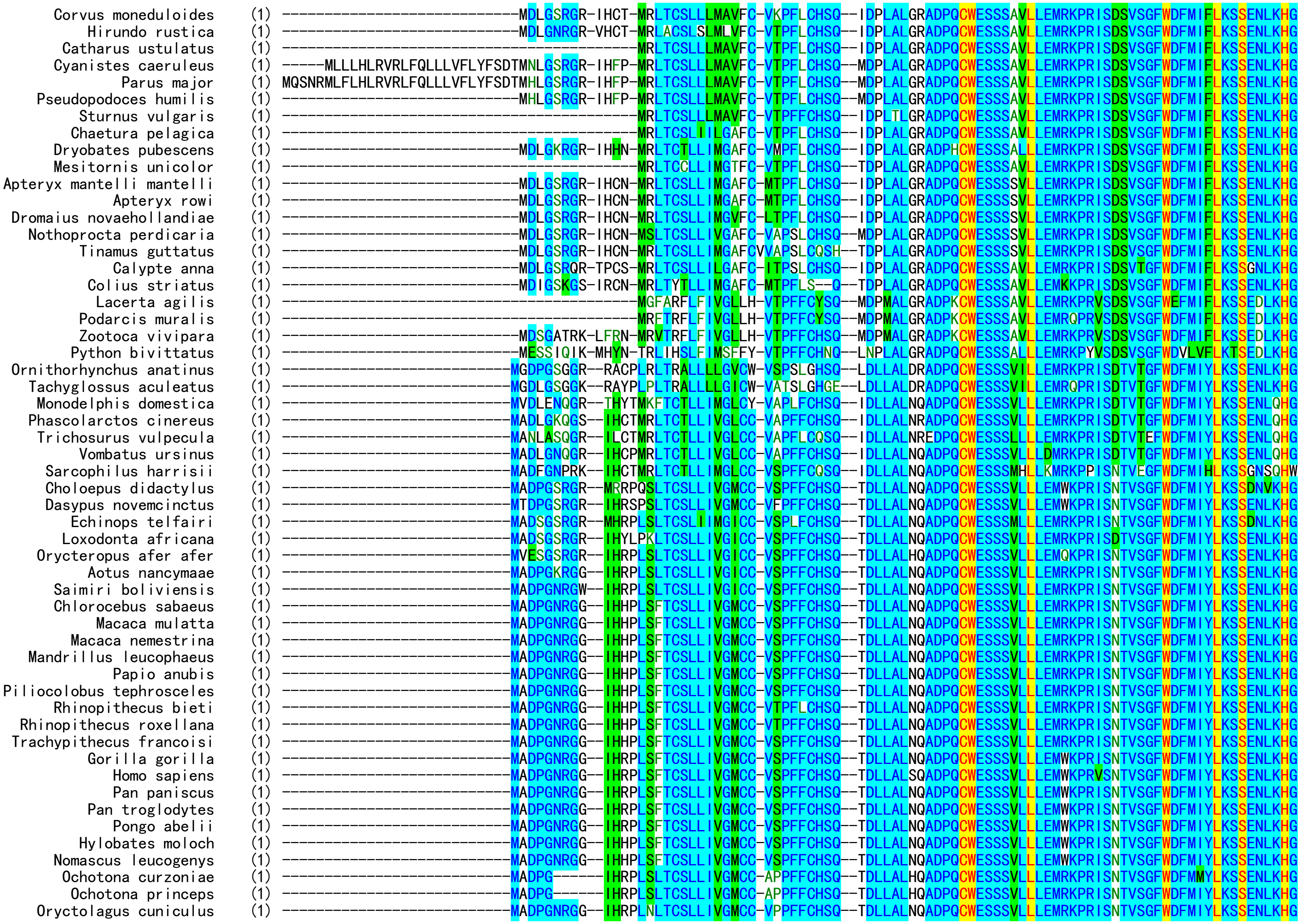
Corvus moneduloides
Hirundo rustica
Catharus ustulatus
Cyanistes caeruleus
Parus major
Pseudopodoces humilis
Sturnus vulgaris
Chaetura pelagica
Dryobates pubescens
Mesitornis unicolor
Apteryx mantelli mantelli
Apteryx rowi
Dromaius novaehollandiae
Nothoprocta perdicaria
Tinamus guttatus
Calypte anna
Colius striatus
Lacerta agilis
Podarcis muralis
Zootoca vivipara
Python bivittatus
Ornithorhynchus anatinus
Tachyglossus aculeatus
Monodelphis domestica
Phascolarctos cinereus
Trichosurus vulpecula
Vombatus ursinus
Sarcophilus harrisii
Choloepus didactylus
Dasypus novemcinctus
Echinops telfairi
Loxodonta africana
Orycteropus afer afer
Aotus nancymaae
Saimiri boliviensis
Chlorocebus sabaeus
Macaca mulatta
Macaca nemestrina
Mandrillus leucophaeus
Papio anubis
Piliocolobus tephrosceles
Rhinopithecus bieti
Rhinopithecus roxellana
Trachypithecus francoisi
Gorilla gorilla
Homo sapiens
Pan paniscus
Pan troglodytes
Pongo abelii
Hylobates moloch
Nomascus leucogenys
Ochotona curzoniae
Ochotona princeps
Oryctolagus cuniculus

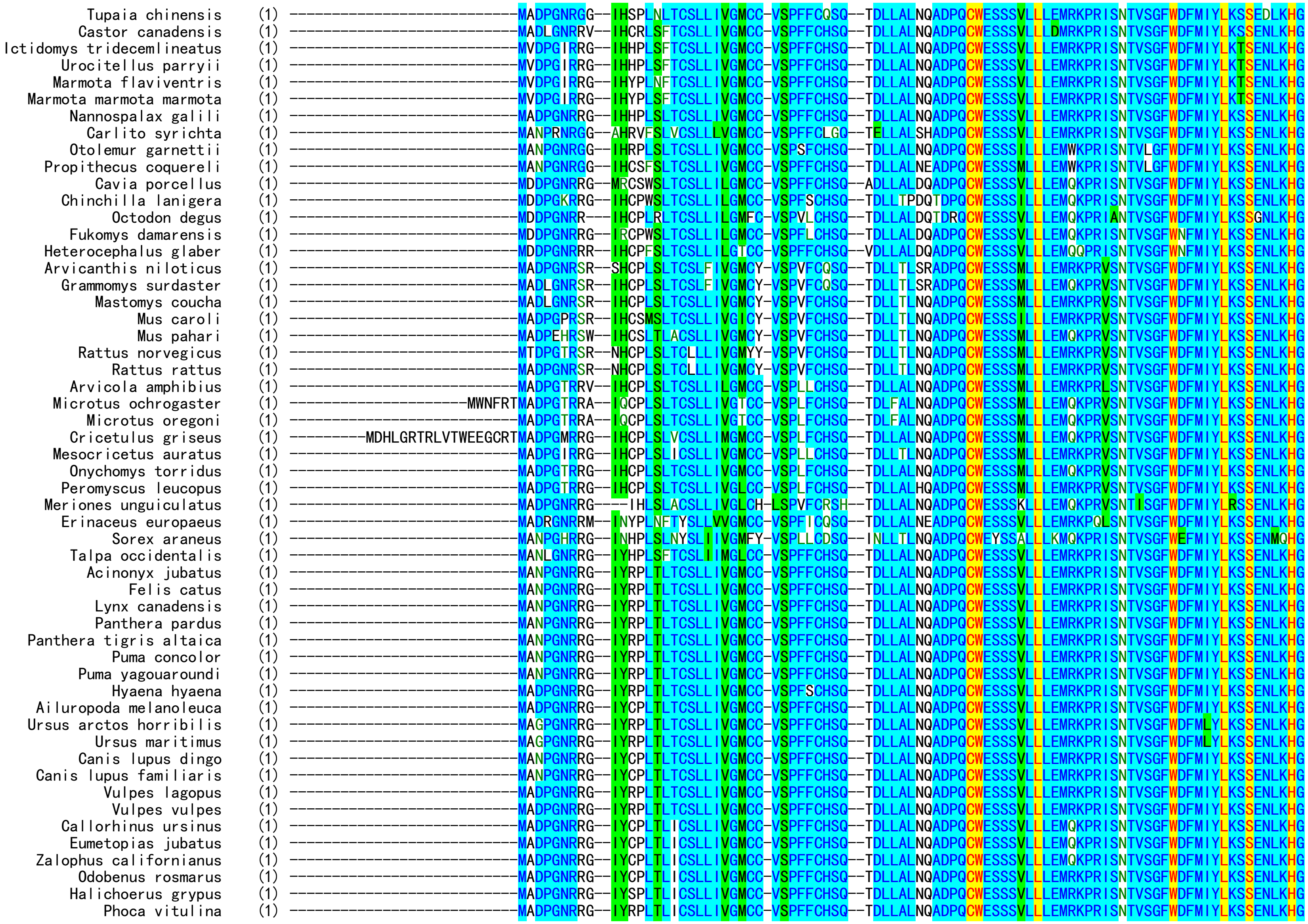

Tupaia chinensis
Castor canadensis
Ictidomys tridecemlineatus
Urocitellus parryii
Marmota flaviventris
Marmota marmota marmota
Nannospalax galili
Carlito syrichta
Otolemur garnettii
Propithecus coquereli
Cavia porcellus
Chinchilla lanigera
Octodon degus
Fukomys damarensis
Heterocephalus glaber
Arvicanthis niloticus
Grammomys surdaster
Mastomys coucha
Mus caroli
Mus pahari
Rattus norvegicus
Rattus rattus
Arvicola amphibius
Microtus ochrogaster
Microtus oregoni
Cricetulus griseus
Mesocricetus auratus
Onychomys torridus
Peromyscus leucopus
Meriones unguiculatus
Erinaceus europaeus
Sorex araneus
Talpa occidentalis
Acinonyx jubatus
Felis catus
Lynx canadensis
Panthera pardus
Panthera tigris altaica
Puma concolor
Puma yagouaroundi
Hyaena hyaena
Ailuropoda melanoleuca
Ursus arctos horribilis
Ursus maritimus
Canis lupus dingo
Canis lupus familiaris
Vulpes lagopus
Vulpes vulpes
Callorhinus ursinus
Eumetopias jubatus
Zalophus californianus
Odobenus rosmarus
Halichoerus grypus
Phoca vitulina

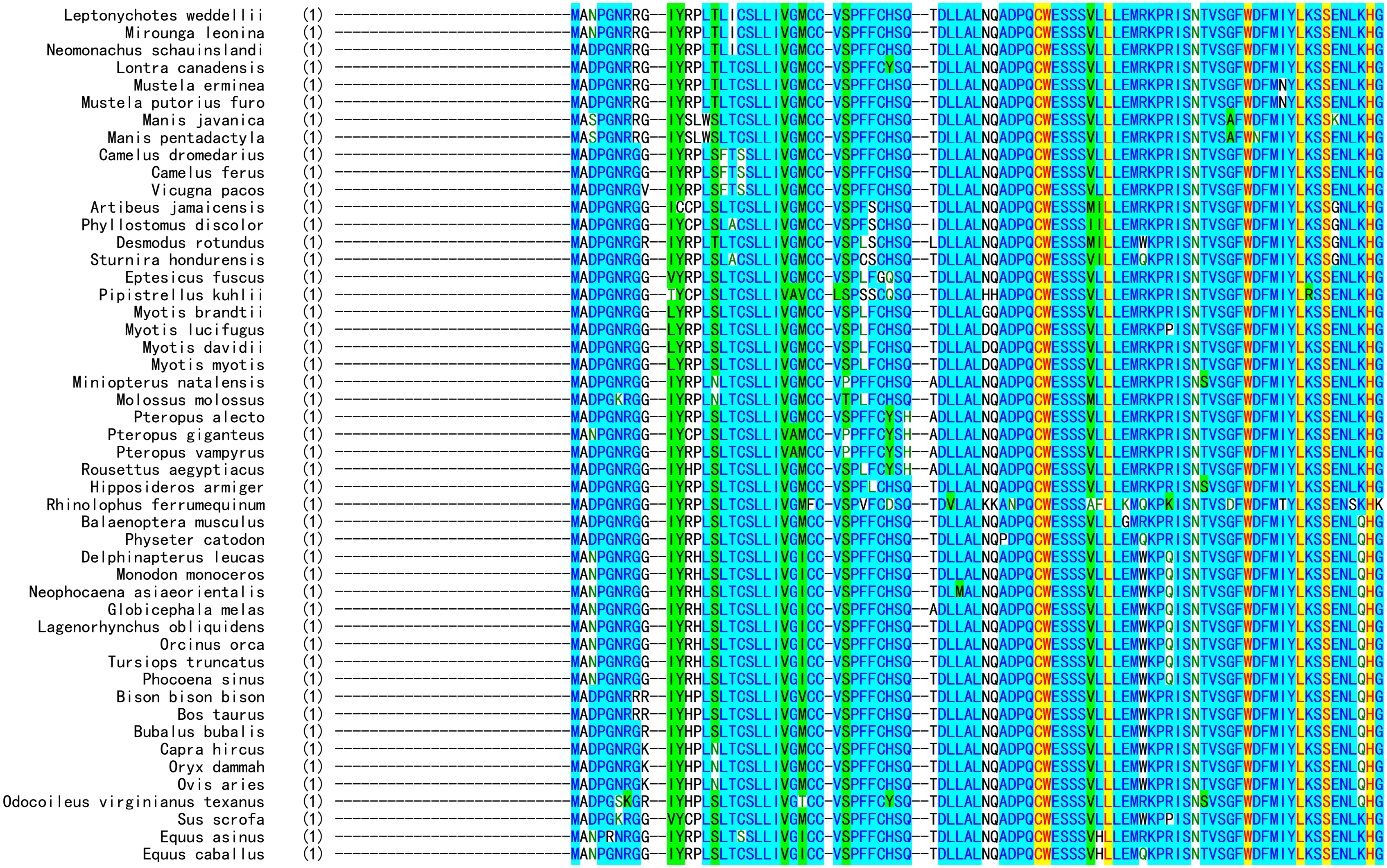
Leptonychotes weddellii (1)
Mirounga leonina (1)
Neomonachus schauinslandi (1)
Lontra canadensis (1)
Mustela erminea (1)
Mustela putorius furo (1)
Manis javanica (1)
Manis pentadactyla (1)
Camelus dromedarius (1)
Camelus ferus (1)
Vicugna pacos (1)
Artibeus jamaicensis (1)
Phyllostomus discolor (1)
Desmodus rotundus (1)
Sturnira hondurensis (1)
Eptesicus fuscus (1)
Pipistrellus kuhlii (1)
Myotis brandtii (1)
Myotis lucifugus (1)
Myotis davidii (1)
Myotis myotis (1)
Miniopterus natalensis (1)
Molossus molossus (1)
Pteropus alecto (1)
Pteropus giganteus (1)
Pteropus vampyrus (1)
Rousettus aegyptiacus (1)
Hipposideros armiger (1)
Rhinolophus ferrumequinum (1)
Balaenoptera musculus (1)
Physeter catodon (1)
Delphinapterus leucas (1)
Monodon monoceros (1)
Neophocaena asiaeorientalis (1)
Globicephala melas (1)
Lagenorhynchus obliquidens (1)
Orcinus orca (1)
Tursiops truncatus (1)
Phocoena sinus (1)
Bison bison bison (1)
Bos taurus (1)
Bubalus bubalis (1)
Capra hircus (1)
Oryx dammah (1)
Ovis aries (1)
Odocoileus virginianus texanus (1)
Sus scrofa (1)
Equus asinus (1)
Equus caballus (1)

```
Acanthochromis polyacanthus (84) TLFWDLARVFWNMYLDCVLSRSHGLGRR---------HITAVHS-LTDKSFRFDSSGG----------NFRTWLSVRVRRRGHIKTLDTKPK--------------------------
Cottoperca gobio (84) ALFWDLARVFWDLYLDCVLSRSHGLGRR---------HIPALHSLITGKSFRFKSSGV----------NSRAWLSVRLRPRGHIKTLKTKSKSNTHYHK-------------------
Perca flavescens (84) ALFWDLARVFWDLYLDCVLSRSHGMGRR---------HITALHSLVTDKPFRFNSSGA----------NSRAWLSVRVMPRGHIKTLKTKPKSHTHYHK-------------------
Larimichthys crocea (84) ALFWDLARVFWDLYLDCVLSRSHGLGRR---------HITAVHSLITDKSFRFNSSGM----------NSRAWLSVRVRRRGHIQTLKTKPKSNTHYHK-------------------
Sparus aurata (86) ALFWDLARVFWDLYLDCVVSRSHGLGRR---------HVTAIHSLITDKSFRFNSSGA----------NSRAWLSVRVRRRGHIKTLKTKPKPNTHYHK-------------------
Parambassis ranga (81) ALFWDLARVFWDLYLDCVLSRSHGLGRR---------HITAVHSLITDKSFRFNSSGT----------NSRAWLSLRVRRRGHIKTLKTNPK--------------------------
Gouania willdenowi (83) ELFWDLARVFWDLYLNCVMSRSHGLGRR---------HVTAVHSLFTDKSFRFNSSGR----------KSRAWLRVRVRSRGHIKTLKTKQQEMHPVINI------------------
Salarias fasciatus (97) ELFWDLARVFWDLYVDCVMSRSHGLGRR---------HTTAVHSLFTDKSFRFNSSGA----------NSRAWLRVRVRRRGHIKTRKTKPK--------------------------
Archocentrus centrarchus (66) ALFWDLAQVFWDIYLDCIVSRSHGLGRR---------HVTAVQSLITDKSFRFNSSGA----------KSHAWLRVRVRRRGQIKTLKTDPK--------------------------
Sphaeramia orbicularis (81) ALFWDLAGVFWDMYADCVLSRSHGLGRR---------HVTAVHSLFTDKTFRFNSSGL----------KTLAWLCVRVRRRGRIETLKTKPNSLYHK---------------------
Mastacembelus armatus (86) ALFWDLAGLFWDIYVDCVLSRSHGLGRR---------HITAVHSLITDKSFRFNSSGV----------NSRAWLSVRVRHRGRIKALKIKPKSNSHYF--------------------
Monopterus albus (84) ALFWDLAGVFWDIYVDCVLSRSHGLGRR---------HITAVHSLITDKSFSFNSSGV----------NSWAWLSVRVRHRGHIKTLKTKPSSNARYYI-------------------
Echeneis naucrates (88) ALFSDLARVFWDMYLDCVMSRSHGLGRR---------HLTAVHSLIPDKTFRFNSSGV----------NSQS-LIIRVRRRGQIKTLKTAPKANTKYHK-------------------
Xiphophorus couchianus (84) ALFWDLARVFWEIYVECVMSRSHGMGRR---------HITSVHSLITDKSFRFNSSRA----------NSWTKLTVRVRRRGHITRLKTEPK--------------------------
Xiphophorus hellerii (84) ALFWDLARVFWEIYVECVMSRSHGMGRR---------HITSVHSLITDKSFRFNSSRA----------NSWTKLTVRVRRRGHITRLKTEPK--------------------------
Xiphophorus maculatus (84) ALFWDLARVFWEIYVECVMSRSHGMGRR---------HITSVHSLITDKSFRYNSSRA----------NSWTKLTVRVRRRGHITRLKTEPK--------------------------
Oryzias latipes (84) ALFWDLARVFWDLYLDCVLSRSHGLGRR---------HITSMHSLISDKSFRFNSSGV----------NSRAWISVRVRRRGHITTLKTNRDK-------------------------
Hippocampus comes (80) TLFWDLARVFWEIYLDCVMSRSHGLGRR---------HVTSVRSLSTDKLFKFNSSGA----------NCSLWLSFRVRCKDHIKTTKSKAT--------------------------
Takifugu rubripes (84) ALFWDLAQVFWDLYLDCVLSRSHGLGRR---------HVTALHSLFTDKSFRLNDSGS----------NYQTWLSVRVRHKGHIKTLRSKVQPSTVYYK-------------------
Etheostoma spectabile (85) VLFWDLVRVFWDLYLDCVLSRSHGMGRR---------HITALQSLITDSKFLVFILMW----------D-------------------------------------------------
Labrus bergylta (87) ALFWDLARVFWDLYLDCVLSRSHGLGRR---------HVTAVHSLITDSK--------------------------------------------------------------------
Maylandia zebra (89) ALFWDLAQVFWDIYLNCIVSRSHGLGRR---------HITAVQSLITDNDGGDNCGTD----------NQSFVVEQEKTTQNPSDSTAQEQSPTRGSM--------------------
Oreochromis niloticus (84) ALFWDLAQVFWDIYLDCIVSRSHGLGRR---------HITAVQSLITDDDGGDNCGTD----------NQSFVVEQEKTTQNPSDSTAQEQSPTRGSM--------------------
Seriola lalandi dorsalis (66) ALYWDLARVFWDIYVDCVLSRSHGLGRR---------HITAVHSLITDKMIQS---------------------------AGHLKLSKSRGE--------------------------
Astyanax mexicanus (89) ALFWDLAQVFWDLYVDCVLSRNHGLGRRHLAQPGER-ITTASRALITDQSFVQESQNHFSK----LKKFSRGWFRIQVQQVGRNNLQHGAHPKSLNIKKVSSF---------------
Chanos chanos (92) ALFWDLAQVFWDIYVDCVLSRTHGLGRRQTSYPQE--RITAMRSLITDKSFVQDVHTNFSK----LKEFTQGWFKIQVQRVGLGGLTGENRLRTKSSSKLQKEFF-------------
Esox lucius (85) ALFWDLAQVFWDIYVDCVMSRTHGLGRRQLKRPRE--QITATRSLITDKSFIQD--TNYSK----LKESAQGCLKIQVQHFGPGVLRHILCARGIKRRSIF-----------------
Oncorhynchus kisutch (95) ALFWDLAQVFWDIYVDCVLSRTHGLGRRQLTWPHE--QITATRSLVTDKSFVQDSQTNVSK----LKESSQGWLKIQVQHFGPGILNHIIRTRGIKSRYIL-----------------
Oncorhynchus nerka (95) ALFWDLAQVFWDIYVDCVLSRTHGLGRRQLTWPHE--QITATRSLVTDKSFVQDSQTNVSK----LKESSQGWLKIQVQHFGPGILNHITRTRGIKSRSIL-----------------
Salmo trutta (83) ALFWDLAQVFWDIYVDCVLSRTHGLGRRQLTWPHE--QITAMRSLVTDKSFVQDSQTNVSK----LKESSQGWLKIQVRHFGPGILNHIIRTRGIKSISIL-----------------
Oncorhynchus tshawytscha (95) ALFWDLAQVFWDIYVDCVLSRTHGLGRRQLTWPHE--QITATRSLVTDSKYLSLT------------------LFSPVYSLGVDTF--------------------------------
Gadus morhua (84) ALFWDLAQVFWDIYVDCVLARSHGLGRRGLKLPKD--RITSVHSLVTDPSQFGTWERG--------SKVTAGLRK-------------------------------------------
Myripristis murdjan (84) ALFWDLAQVFWDIYVDCVLSRTHGLGRRQLKWPNG--HITAVHSLITDSK--------------------------------------------------------------------
Paramormyrops kingsleyae (86) ALFWDLARLFWDIYVDCVLSRTHGLGRRRLATPSS--PATPADSWYHSLKH-------------------------------------------------------------------
Scleropages formosus (84) ALFWDLARLFWDIYVDCVLSRTHGLGRRQLTTSQQ--RITAMYSLVTDQSYVHVSQSNFHD----VPDQDPSTSSHTELLGIHQQRHGTKRNSAS-----------------------
Erpetoichthys calabaricus (83) ALFWNLAQVFWDIYVDCVLSRTHGLGRRQ-LTAGDR-EITALHSLVTDESYIRGHRKDFGK----EKQYTRDLIGIHVHQSGPGGPPPLDQALSFRRPRSRV----------------
Anolis carolinensis (81) DVFNDLAQTFWNMYIDCLLSRSHGVGKRQ----LRFSRNTLTNAQETSEGVANFNSRARR----------------------------------------------------------
Ficedula albicollis (82) ELFNDLAQNFWDMYVDCVLSRSHGMGRRQ----LTSPKYSSTYSHRTLEGSAFTNPF-------------------------------------------------------------
Geotrypetes seraphini (87) ALFDDLAQNFWNIYVDCVLSRSHGLGRRH----LMPPEYVFAYSAKAEAGFDFTKHPF------------------------------------------------------------
Callorhinchus milii (78) ALFLDLAQVFWDTYVDCVLSRSHGLGRR-----QIIARYPLKYPKYTAAGLTRSSDKKLRI---------------------------------------------------------
Bufo bufo (88) VLFWDLAQLFWDIYVDCVLSRTHGLGRRQLKEEGKK--ISNFVSQFTSKRFS----HNRRSVLPSQKELIQHLIDIQVDKSESRLLRSIKSGIKRK----------------------
Nanorana parkeri (88) VLFWDLAQLFWDIYIHCVLSRAHGLGRRQLNEEETK--VTNFISQYTSKTFS----QNRRPLPPDWEELIEQLVGIHVNKSESRLLGNIKRRVKRK----------------------
Rana temporaria (88) ILFWDLAQLFWDIYVHCVLSRAHGLGRRQLNEGEKK--VTNFISQYTSKTFS----QNRRPLSKQWGELLEQLKGINVYKSESRLLGIIKRHMKRK----------------------
Microcaecilia unicolor (76) ALFWDLAQLFWDIYVDCVLSRTHGLGRRQLAEEEQK--ITNLQLQVTGSKRS-SVSHKCV----------------------------------------------------------
Rhinatrema bivittatum (88) ALFWDLAQLFWDIYVDCVLSRTHGLGRRQLTEEEQK--ITNLHLQVTGSMQK-TFSQGPSIPLLKKRELIEGSIGIHVQKSGSRMLERVIKAMDLKRK--------------------
Alligator sinensis (88) ALFWDLAQLFWDIYVDCVLSRTHGLGRRQLAEAEQR--VTTLHSHITGRKQG-AFSHLQRILALKKKELIEDFVSIHVHKSGSTLLGRVTGDLGIKRKQHL-----------------
Crocodylus porosus (88) ALFWDLAQLFWDIYVDCVLSRTHGLGRRQLAEAEQR--VTTLHSHITDRKQG-AFSHLQRTLALKKKMLIEDFVSIHVHKSGSTLLGRVTGDLGIKRKQHL-----------------
Gavialis gangeticus (88) ALFWDLAQLFWDIYVDCVLSRTHGLGRRQLAEAEQR--VTTLHSHITGRKQG-AFSHLQRTLALKKKMLIEDFVSIHVDKSGSTLLGRVTGDLGMKRKQHL-----------------
Chelonia mydas (88) ALFWDLAQLFWDIYVDCVLSRTHGLGRRQLAEAEQK--IIALHSQLTERKQG-TFAHNQRTRVLNKKELIEDLISIHVHKSGSALLRRVIGGLGIKRKQFV-----------------
Chelonoidis abingdonii (88) ALFWDLAQLFWDIYVDCVLSRTHGLGRRQLAEAEQK--ITALHSQLTERKQG-TFSHNQRTQVLNKKELIEDLISIHVHKSGSALLRRVIGGLGIKRKQ-------------------
Gopherus evgoodei (88) ALFWDLAQLFWDIYVDCVLSRTHGLGRRQLAEAEQK--ITALHSQLTERKQG-TFSHNQRTRVLNKKELIEDLS-IHVHKSGSALLRRVIGGLGIKRKQ-------------------
Mauremys reevesii (88) ALFWDLAQLFWDIYVDCVLSRTHGLGRRQLAEAEQK--ITALHSQLTERKQG-TFSHNQRKRVLNKKELIEDLISIHVHKSGSALLRRVIGGLGIKRKQ-------------------
Chrysemys picta (87) ALFWDLAQLFWDIYVDCVLSRTHGLGRRQLAEAEQK--ITALHSQLTEQKQG-TFSHNQRTRVLNKKELIEDLS-IHVHKSGSALLRRVIGGLGIKRKQ-------------------
Trachemys scripta elegans (87) ALFWDLAQLFWDIYVDCVLSRTHGLGRRQLAEAEQK--ITALHSQLTEQKQG-TFSHNQRTRVLNKKELIEDLS-IHVHKSGSALLRRVIGGLGIKRKQ-------------------
```

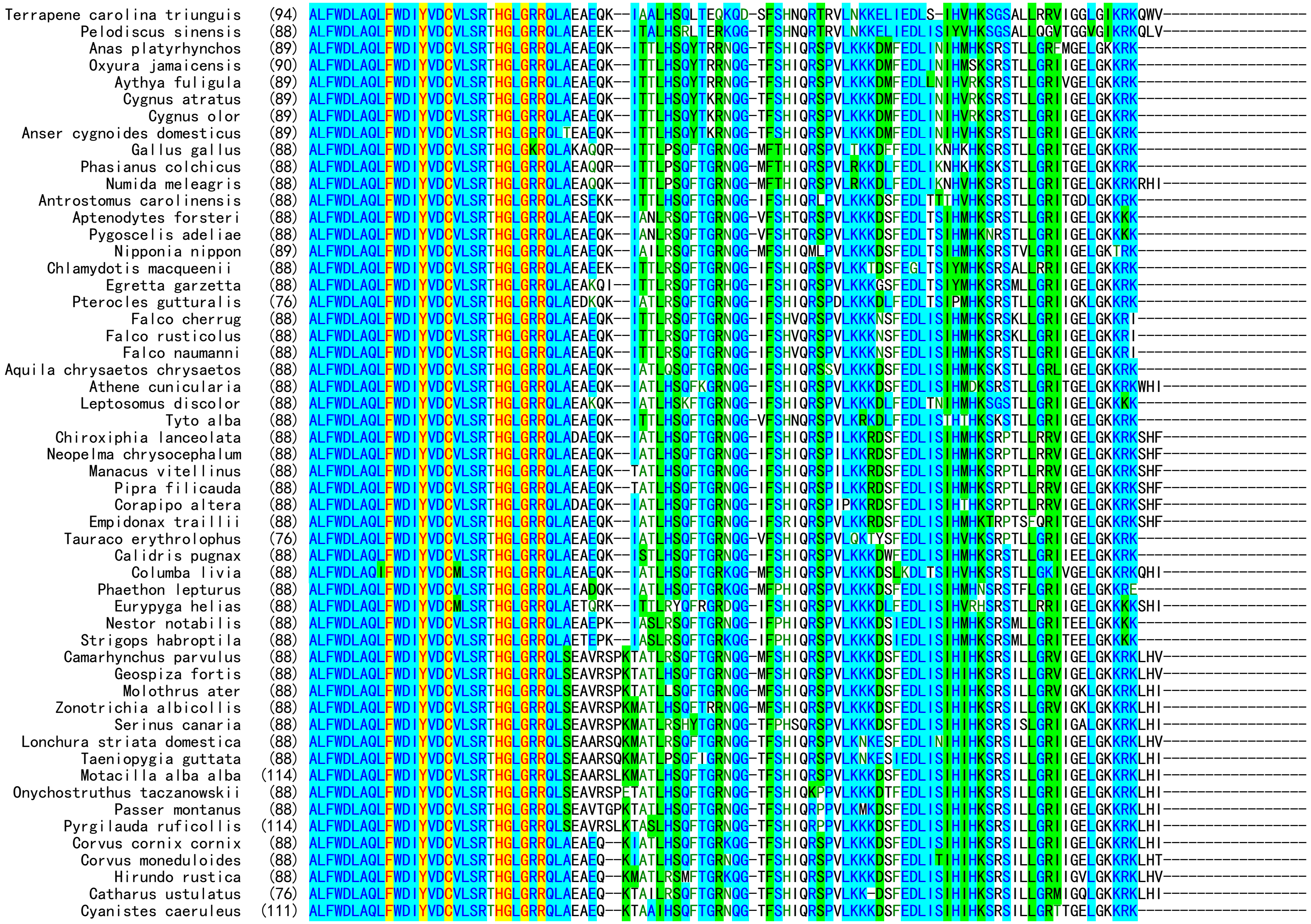

```
                 Parus major  (116) ALFWDLAQLFWDIYVDCVLSRTHGLGRRQLAEAEQ--KTATIHSQFTGKNQG-TFSHIQRSPVLKKKDLFEDLISIHIHKSRSILLGRTTGELGKKRK--------------------
       Pseudopodoces humilis   (88) ALFWDLAQLFWDIYVDCVLSRTHGLGRRQLAEAEQ--KTATIHSQFTGKNQG-TFSHIQRSPVLKKKDSFEDLISIHIHKSRSILLGRTTGELGKKRK--------------------
            Sturnus vulgaris   (76) ALFWDLAQLFWDIYVDCVLSRTHGLGRRQLAKAEQ--KTATLRSQFTGRNQG-TFSHIQRSPVLKK-DSFEDIISIHIDKSRSILLGRITGELGKKRK--------------------
           Chaetura pelagica   (76) ALFWDLAQLFWDIYVDCVLSRTHGLGRRQLAEAEQ---IATLHSQFTRRNQG-IFSHIQRS--LKKKDSFEDLTSIHMHRSNSRLLGRIIGELGKKRKLHT-----------------
          Dryobates pubescens  (88) ALFWDLAQLFWDIYVDCVLSRTHGLGRRQLADAEQK--IATLHSQFMGRNQG-IFPHIQRSPALKKKGSFEDLN-IREHKKKSALLGRIIGQLGKKRELHI-----------------
          Mesitornis unicolor  (76) ALFWDLAQLFWDIYVDCVLSRTHGLGRRQLAEAEQK--IATLHSQFKERNQG-IFPHIQRSPVLKKKGSFEDLTSIHMHKSRSTLLGRTIGETGKKEKKTHLVQDLDL----------
   Apteryx mantelli mantelli   (88) ALFWDLAQLFWDIYVDCVLSRTHGLGRRQLAEAEQK--ITTLHSQFTWRNQG-IFSHIQRMPVLKKKDVFEDLISVYVQKSRSTLLGRVTEELGKKRK--------------------
                Apteryx rowi   (88) ALFWDLAQLFWDIYVDCVLSRTHGLGRRQLAEAEQK--ITTLHSQFTWRNQG-IFSHIQRMPVLKKKDVFEDLISVYVQKSRSTLLGRVTEELGKKRK--------------------
     Dromaius novaehollandiae  (88) ALFWDLAQLFWDIYVDCVLSRTHGLGRRQLAEAEQK--ITTLHSQFTRRNQG-IFSHIQRMPVLKKKDVFEDLISIHVQKSRSTLLGRVTRELGKKRK--------------------
        Nothoprocta perdicaria (88) ALFWDLAQLFWDIYVDCVLSRTHGLGRRQLAEAEHK--ITALHSQFTQRNQG-MFSHIQRMAVLKKKDVFEDLISIHVQKSRSTLLGRVTEELGKKRK--------------------
            Tinamus guttatus   (89) ALFWDLAQLFWDIYVDCVLSRTHGLGRRQLAEAEQK--ITTLHSQFTRKKQG-TFSHIQRMAILKKKDVFEDLISIHVQKSRSTLLGRVTGDLGKKRK--------------------
               Calypte anna    (88) ALFWDLAQLFWDIYVDCVLSRTHGLGRRQLAEAGQK--ITALPSQVTGKKQG-LFSCIQKLPVLKKKDWFEDLKSLHMPKRRSTWLGRIMGELGKKRK--------------------
             Colius striatus   (86) ALFWDLAQIFWDIYVECVLSRTHGLGRRQLAEAEQ---ITILHSQFTGKKKG-TFSHIQRSSVLKKKDMFEDLISIHMHKSRS-ALLRIIGELGEKRNWYI-----------------
              Lacerta agilis   (76) ALFWHLAQIFWDIYMDCVLSRTHGLGKRQLAEAQRR--TAALPSWLILSKQG-IFSQIQMTPLWKKKELKEDIIRIHVHKSRSGSHKRITGYLRMKVKLF------------------
            Podarcis muralis   (76) ALFWHLAQLFWDIYMDCVLSRTHGLGKRQLAEAQRQ--TAALPSWLILSKQG-IFSQIQMTPLWKKKELKENLIRIHVHKSRSASHGRITGLLTIKAKLF------------------
            Zootoca vivipara   (88) ALFWHLAQLFWDIYMDCVLSRTHGLGKRQLAKAQRQ--TAALPSWLILSKQG-IFSQIQMTPLWKKKELKENFIRIHVHKSRSASHGRITGHLTMKAKLF------------------
           Python bivittatus   (88) ALFWNLAQLFWDIYVDCVLSRTHGLGKRQLAGAQHQ--ITAFPSWITRRKQ---------------------------------------------------------------------
     Ornithorhynchus anatinus  (89) ALFWDLAQLFWDIYVDCVISRTHGLGRRQLDGGEEE-KITALHSQTTGGRRG-TYSQFPRTPLLKKKELIEDLISIHVLKSGSRLVGRVTS---LKRK--------------------
       Tachyglossus aculeatus  (89) ALFWDLAQLFWDIYVDCVISRTHGLGRRQLAGEEEE-KIPALHSRTAGG----TYSQEPRTPLLKKKELIEDLISIHVLKSGSRLVGRVTS---LKRK--------------------
         Monodelphis domestica (89) ALFWDLAQLFWDIYVDCVLSRNHGLGRRQLARDE-E-KLSTMHAQHTGS----SYSQFLRAPLLKKKGLIEDLISIHMHKSGSRLVGRVTSDLGIKRK--------------------
       Phascolarctos cinereus  (89) ALFWDLAQLFWDIYVDCVLSRNHGLGRRQLAGDE-E-KLSTMHSQHSGS----TYSQFLRTSLLKKKELIEDLISIHVHKSGSRLIGRVNSDLGIKRK--------------------
        Trichosurus vulpecula  (89) ALFWDLAQLFWDIYVDCVLSRNHGLGRRQLAEDE-E-KLSTMHSQHSGSKSG-TYSQFLRAPLLKKKELIEDLISIHVQKSGSRLIGRVTNDLGMKRK--------------------
            Vombatus ursinus   (89) ALFWDLAQLFWDIYVDCVLSRNHGLGKRQLAGDE-E-KLSTMHSQHSGSKSG-TYSQFQRAPLLKKKELIEDLISIHVHKSGSRLIGRVNSDLGIKRK--------------------
         Sarcophilus harrisii  (89) ALFWDLAQLFWDIYIDCVLSRNHGLGKRQLAGDK-E-KLSTTNLQHTGS----TYSQLLRAPLLKKKELIEDLMSIHKHKSGSRLI--------------------------------
         Choloepus didactylus  (89) ALFWDLAQLFWDIYVDCVLSRNHGLGRRQLARQE-E-KIPAAHSQHTGSQQG-VYFQPLRTPFLKKKEMIEDLISIYVHKSGSRFIGKVTSGLKIKRK--------------------
         Dasypus novemcinctus  (89) ALFWDLAQLFWDIYVDCVLSRNHGLGRRQLDGEE-E---------HTGRKQG-AYSQLLKTTFLKKKEMIEDLISMYVRKSGSGFIGKMTSGLEIKRK--------------------
           Echinops telfairi   (89) ALFWDLAQLFWDIYVDCVLSRNHGLGRRQLAGEE-D-RIPAAHSGHTRSKQG-AYSQLLRTPLLKKKELVEELMHMYVPRSGSRLIGKVTSSLRIKRK--------------------
          Loxodonta africana   (89) ALFWDLAQLFWDIYVDCVLSRNHGLGRRQLAGEE-E-RIPAAHSGHTRSKQG-SYSQLLRIPFLKKKELVEDLINMYVSRSGSRVIGEVTSGLRIKRKLI------------------
         Orycteropus afer afer (89) ALFWDLAQLFWDIYVDCVLSRNHGLGRRQLAGEE-D-RISAAQSGHTRSKQD-AYSHLLRTPFLKKKELVEDVINMYVPRSGSRFTGKVTSGLKIKRK--------------------
            Aotus nancymaae    (89) ALFWDLAQLFWDIYVDCVLSRNHGLGRRQLVGEE-E-KISAAQLQHTRSKQG-TYSQLLRTAFLKKKELIEDLISMHVHRSGSSVIGKVN--QKIKRK--------------------
          Saimiri boliviensis  (89) ALFWDLAQLFWDIYVDCVLSRNHGLGRRQVVGEE-E-KISAAQPQHTRSKQG-TYSQLLRIAFLKKKELIEDLISMHVHRSGSSVIGKVN--QEIKRR--------------------
         Chlorocebus sabaeus   (89) ALFWDLAQLFWDIYVDCVLSRNHGLGRRQLAGEE-E-KISAAQPQHTRSKQG-TYSQQLRTSLLKKKELIEDLISMHVRRSGSSVIGKVN--LEIKRK--------------------
              Macaca mulatta   (89) ALFWDLAQLFWDIYVDCVLSRNHGLGRRQLAGEE-E-KISAAQPQHTRSKQG-TYSQQLRTSLLKKKELIEDLISMHVRRSGSSVIGKVN--LEIKRK--------------------
            Macaca nemestrina  (89) ALFWDLAQLFWDIYVDCVLSRNHGLGRRQLAGEE-E-KISAAQPQHTRSKQG-TYSQQLRTSLLKKKELIEDLISMHVRRSGSSVIGKVN--LEIKRK--------------------
       Mandrillus leucophaeus  (89) ALFWDLAQLFWDIYVDCVLSRNHGLGRRQLAGEE-E-KISAAQPQHTRSKQG-TYSQQLRTSLLKKKELIEDLISMHVRRSGSSVIGKVN--LEIKRK--------------------
                Papio anubis   (89) ALFWDLAQLFWDIYVDCVLSRNHGLGRRQLAGEE-E-KISAAQPQHTRSKQG-TYSQQLRTSLLKKKELIEDLISMHVRRSGSSVIGKVN--LEIKRK--------------------
    Piliocolobus tephrosceles  (89) ALFWDLAQLFWDIYVDCVLSRNHGLGRRQLAGEE-E-KISAAQPQHTRSKQG-TYSQQLRTSLLKKKELIEDLISMHVRRSGSSAIGKVN--LEIKRK--------------------
         Rhinopithecus bieti   (89) ALFWDLAQLFWDIYVDCVLSRNHGLGRRQLAGEE-E-KISAAQPQHTRSKQG-TYSQQLRTSLLKKKELIEDLISMHVRRSGSSAIGKVN--LEIKRE--------------------
      Rhinopithecus roxellana  (89) ALFWDLAQLFWDIYVDCVLSRNHGLGRRQLAGEE-E-KISAAQPQHTRSKQG-TYSQQLRTSLLKKKELIEDLISMHVRRSGSSAIGKVN--LEIKRE--------------------
      Trachypithecus francoisi (89) ALFWDLAQLFWDIYVDCVLSRNHGLGRRQLAGEE-E-KISAAQPQHTRSKQG-TYSQQLRTSLLKKKELIEDLISMHVRRSGSSAIGKVN--LEIKRE--------------------
              Gorilla gorilla  (89) ALFWDLAQLFWDIYVDCVLSRNHGLGRRQLVGEE-E-KISAAQPQHTRSKQG-TYSQLLRTSFLKKKELIEDLISMHVRRSGSSVTGKVN--LEIKRK--------------------
                Homo sapiens   (89) ALFWDLAQLFWDIYVDCVLSRNHGLGRRQLVGEE-E-KISAAQPQHTRSKQG-TYSQLLRTSFLKKKELIEDLISMHVRRSGSSVIGKVN--LEIKRK--------------------
                Pan paniscus   (89) ALFWDLAQLFWDIYVDCVLSRNHGLGRRQLVGEE-E-KISAAQPQHTRSKQGGTYSQLLRTSFLKKKELIEDLISMHVRRSGSSVIGKVN--LEIKRK--------------------
             Pan troglodytes   (89) ALFWDLAQLFWDIYVDCVLSRNHGLGRRQLVGEE-E-KISAAQPQHTRSKQG-TYSQLLRTSFLKKKELIEDLISMHVRRSGSSVIGKVN--LEIKRK--------------------
                Pongo abelii   (89) ALFWDLAQLFWDIYVDCVLSRNHGLGRRQLVGEE-E-KISAAQPQHTRSKQG-TYSQLLRTSFLKKKELIEDLISMHVRRSGSSVIGKVN--LEIKRK--------------------
             Hylobates moloch  (89) ALFWDLAQLFWDIYVDCVLSRNHGLGRRQLVGEE-E-KISAAHPQHTRSKQG-TYSQLLRTSFLKKKELIEDLISMHVRRSGSSVIGKVN--LEIKRK--------------------
          Nomascus leucogenys  (89) ALFWDLAQLFWDIYVDCVLSRNHGLGRRQLVGEE-E-KISAAHPQHTRSKQG-TYSQLLRTSFLKKKELIEDLISMHVRRSGSSVIGKVN--LEIKRK--------------------
           Ochotona curzoniae  (85) ALFWDLAQLFWDIYVDCVLSRNHGLGRRQLTGEE-E-KIPAAQPQHTGSKQG-AFSQLPRMLFLKKKELIQDLINMHAHRSGSRFIGKTK--FEIKRK--------------------
            Ochotona princeps  (85) ALFWDLAQLFWDIYVDCVLSRNHGLGRRQLTGEE-E-KIPAAQPQHTGSKQG-TFSQLPRMLFLKKKELIQDLINMHAHRSGSRFIGKTK--LEIKRK--------------------
         Oryctolagus cuniculus (89) ALFWDLAQLFWDIYVDCVLSRNHGLGRRQLAGEE-E-KISAAQPQYTGSKQD-TFSQLPRTLFLKKKELIGDLISMHVQRSGSRFVGKMN--LEIKKK--------------------
            Tupaia chinensis   (89) ALFWDLAQLFWDIYVDCVLSRNHGLGRRQLAEEE-K-KISATQPQHTGSKQG-TYPQRLRTPLLKKKELIEGLISTHVRRSRSRFIGKVN--LEIKRK--------------------
            Castor canadensis  (89) ALFWDLAQLFWDIYVDCVLSRNHGLGRRQLAGKE-E-KISGVQAQRTGSKKG-TYSQLLRTPFLKKKELIEDSISMHVRKSGSRFTGKVN--LEIKRK--------------------
    Ictidomys tridecemlineatus (89) ALFWDLAQLFWDIYVDCVLSRNHGLGRRQLSVKE-E-KISAAQPQHTGNKQG-AYSQLLRVPFLKKKELIEDWISMYVRRSGSRFVGKVN--LEIKRK--------------------
          Urocitellus parryii  (89) ALFWDLAQLFWDIYVDCVLSRNHGLGRRQLSVKE-E-KISAAQPQHXGNKQG-AYSQLLRVPFLKKKELIEDWISMYMRRSGSRFVGKVN--LEIKRK--------------------
```

```
           Marmota flaviventris  (89) ALFWDLAQLFWDIYVDCVLSRNHGLGRRQLSVKE-E-KISAAQPQHTGNKQG-AYSQILRAPFLKKKELIEDWISMYVRRSGSRFVGKVN--LEIKRK--------------------
       Marmota marmota marmota  (89) ALFWDLAQLFWDIYVDCVLSRNHGLGRRQLSVKE-E-KISAAQPQHTGNKQG-AYSQILRAPFLKKKELIEDWISMYVRRSGSRFVGKVN--LEIKRK--------------------
              Nannospalax galili  (89) ALFWDLAQLFWDIYVDCVLSRNHGLGRRQLAG-E-E-KSSAVQLQHSGSKQG-VYSQRLRTPFLKKKELIEDLISMHMRGSGSKFNGKLN--LEIKRK--------------------
               Carlito syrichta  (89) ALFWD--QLFGDIYVDCVLSRNHGLGRRQLTGEE-E-KVSAAQPRHMGKKQG-VYFQLLRPPFLKKKEMIENLISMHVPRVGPRGTGRVS--LEMKRK--------------------
              Otolemur garnettii  (89) ALFWDLSQLFWDIYVDCVLSRNHGLGRRQLATEE-E-KISAAQSQSTGSKQG-IYSQLLRTPSLKKKELIEDLISMHVRKSRSRFIGKVN--LARKRK--------------------
           Propithecus coquereli  (89) ALFWDLSQLFWDIYVDCVLSRNHGLGRRQLTAEE-E-KISAVQPQNIGSKQG-LYSQLLRTPFLKKKELIEDLINMHVRRSGSRFTGKVN--LETKRK--------------------
                Cavia porcellus  (89) ALFWDLAQLFWDIYVDCVLSRNHGLGRRQLAREG-D-TVSAARPQHGEGKQG-VYSQLLRSPFLKKKELIEDLISMRIRRSGSRFVGKVK--LEIKRK--------------------
             Chinchilla lanigera  (89) ALFWDLAQLFWDIYVDCVLSRNHGLGRRQLTGEG-D-KISAAQPQHAGSKQG-AYSQLLRTPFLKKKELIEDLISMHVRRSGSRFLGKVN--LEIKRKLI------------------
                  Octodon degus  (88) ALFWDLAQLFWDIYVDCVLSRNHGLGRRQLTGEG-V-KASATQPQHAGSKQG-EYSQFLRIPFLKKKELIEDLISMHVRTTGSRFVGKVN--LEIKRK--------------------
             Fukomys damarensis  (89) ALFWDLAQLFWDIYVDCVLSRNHGLGRRQLAGEG-E-KISSAQPQHAGSKQG-AYSQLLRTPFLKKKEFIEELISLHVRKSGSRFVGKVN--LEIRRK--------------------
          Heterocephalus glaber  (89) ALFWDLAQLFWDIYVDCVLSRNHGLGRRQLAGEG-E-KVSSAQPQHAGSKKG-AYSQLLRTPFLKKKELIEDLISMHVRRSGSRFIGKVN--LEIKRK--------------------
          Arvicanthis niloticus  (89) ALFWDLAQLFWDIYVDCVLSRNHGLGRRELAGEE-E-KVSKVLPRHLGIKQG-TYSQLLRTLFLKKKELIEDLTSMQTQRRGSRFTGKKK--LEVKRK--------------------
             Grammomys surdaster  (89) ALFWDLAQLFWDIYVDCVLSRNHGLGRRELAKEE-E-KVSKVLPRHLEIKQG-AYSQLLRTLFLKKKELIEDLTSMQTQRRGSRFTGKEK--LEVKRN--------------------
               Mastomys coucha  (89) ALFWDLAQLFWDIYVDCVLSRNHGLGRRELAGGE-E-KVSKVL-RHSGIKQG-AYSQLLRTLSLKKKELIGDLTSMQTQRRGSRFTGKEK--LEVKRK--------------------
                     Mus caroli  (89) ALFWDLAQLFWDIYVDCVLSRNHGLGRRELTGEE-E-KVSKVLTRHSGIKQG-AYSQLLRTLFLKKKELIEELTSMQTQRRGSRFTGKKK--LEVKRK--------------------
                     Mus pahari  (89) ALFWDLAQLFWDIYVDCVLSRNHGLGRRELTGEE-E-KVSKVLTRHSGIKQG-AYSQLLRTLFLKKKELIEDLTSMQTQRRGSRFTGKEK--LEVKRK--------------------
              Rattus norvegicus  (89) ALFWDLAQLFWDIYVDCVLSRNHGLGRRELAGEE-Q-RVSKVLPRHLGIKQG-AYSQLLRSIYLKKKELIGDLTSMQLQKWGPGFTGKEK--LQIKRNRS------------------
                  Rattus rattus  (89) ALFWDLAQLFWDIYVDCVLSRNHGLGRRELAGEE-Q-KVSKVLPRHLGIKQG-AYSQLLRTIYLKKKELIGDLTSMQLQKWGPGFTGKEK--LQMKRNRS------------------
              Arvicola amphibius  (89) ALFWDLAQLFWDIYVDCVLSRNHGLGRRELAGKEKK-EEKSSTVLHSGIKQG-ANPQLLRTPFSKKKELIEDLISVYMHRQASGFTGKVK--LQVKRKESHLQS--------------
             Microtus ochrogaster  (95) ALFWDLAQLFWDIYVDCVLSRNHGLGRRELAGEEKK-EGKSSTVLHSGIKQG-AYSQLLRSPFLKKKELIEDVISVYMHRRASGFTGKAK--LQVKRKESHLQS--------------
               Microtus oregoni  (89) ALFWDLAQLFWDIYVDCVLSRNHGLGRRELAGEEKK-EEKSSTVLHSGITQG-AYSQLLRSPFLKKKELIEDLISVYMHRRASGFTGKAK--LQVKRKESHLQS--------------
             Cricetulus griseus (107) ALFWDLAQLFWDIYVDCVLSRNHGLGRRELAGEGKK-EEKSSTVMHSGITQG-AYSQPLRSPFLKKKELIEDLISLHMRRRGPRFTGKVK--LQVKRK--------------------
            Mesocricetus auratus  (89) ALFWDLAQLFWDIYVDCVLSRNHGLGRRELAGEGKK-EEKSSTVMHSGITQD-AYSQLLRTPFLKKKELIEDLINMHMHQRGSRFTGKVK--LPGKRK--------------------
              Onychomys torridus  (89) ALFWDLAQLFWDIYIDCVLSRNHGLGRRELAGEEKK-EKQSSTVLHLGIKQG-AYSHLLRTPFLKKKELIGDLISVHMRRRGSRIGGKEK--SEVKRKQSHLQS--------------
            Peromyscus leucopus  (89) ALFWDLAQLFWDIYVDCVLSRNHGLGRRELAGEEKK-EKQSSTVLPSGIKQG-AYSQLLRTPFLKKKKLIGDLISMYMRRRGSRVGGKEK--PEVKRKQSYLQN--------------
           Meriones unguiculatus  (87) ALFWDLAQLFWDIYVDCVLSRNHGLGRRELAGEE-D-KISTVLQRRSEAKQD-AYSQLLRTASSKKKELAEDLISGHMHRPASGFTGKVK--LEVKRKMRLPSELTRALGLRKKIFSY
             Erinaceus europaeus  (89) ALFWDLAQLFWDIYVDCVLSRNHGLGRRELAGEE---K----KPQHARTKQG-AYYRQLRTAFLKNKELIEELLSIHARRSGVKFMGKATSGLERKRK--------------------
                  Sorex araneus  (89) ALFWNLAQLFWDIYIDCILSRNHGLGRRQLAGEG-K-KISDVYPQHTVRKRG-AYSQKLRTPFLKKKELIQDLISMRERRSGSKFIIKVTRDLEIKRK--------------------
            Talpa occidentalis  (89) ALFWDLAQLFWDIYVDCILSRNHGLGRRQLAVEE-G-AISTVHPQHKGIKQG-VSPQQLRTPFLKKKELIEDWISMNLRRSGFKLIGKVTGGLEIKRK--------------------
               Acinonyx jubatus  (89) ALFWDLAQLFWDIYVDCVLSRNHGLGRRQLAGKE-E-EISAVRPQHTGRKQG-AYSQQLRTPFLKKKELIEGVISMHLHGSGSKLIGKVTRGLEIKRK--------------------
                    Felis catus  (89) ALFWDLAQLFWDIYVDCVLSRNHGLGRRQLAGKE-E-EISAVRPQHTGRKQG-AYSQQLRTPFLKKKELIEGVISMHLHGSGSKLIGKVTRGLEIKRK--------------------
               Lynx canadensis  (89) ALFWDLAQLFWDIYVDCVLSRNHGLGRRQLAGKE-E-EISAVRPQHTGRKQG-AYSQQLRTPFLKKKELIEGVISMHLHGSGSKLIGKVTRGLEIKRK--------------------
               Panthera pardus  (89) ALFWDLAQLFWDIYVDCVLSRNHGLGRRQLAGKE-E-EISAVRPQHTGRKQG-AYSQQLRTPFLKKKELIEGVISMHLHGSGSKLIGKVTRGLEIKRK--------------------
         Panthera tigris altaica  (89) ALFWDLAQLFWDIYVDCVLSRNHGLGRRQLAGKE-E-EISAVRPQHTGRKQG-AYSQQLRTPFLKKKELIEGVISMHLHGSGSKLIGKVTRGLEIKRK--------------------
                 Puma concolor  (89) ALFWDLAQLFWDIYVDCVLSRNHGLGRRQLAGKE-E-EISAVRPQHTGRKQG-AYSQQLRTPFLKKKELIEGVISMHLHGSGSKLIGKVTRGLEIKRK--------------------
             Puma yagouaroundi  (89) ALFWDLAQLFWDIYVDCVLSRNHGLGRRQLAGKE-E-EISAVRPQHTGRKQG-AYSQQLRTPFLKKKELIEGVISMHLHGSGSKLIGKVTRGLEIKRK--------------------
                  Hyaena hyaena  (89) ALFWDLAQLFWDIYVDCVLSRNHGLGRRQLAGKE-E-KISTVHPQHTERKQG-VYSQQLRTPFLKKKELIEGVISMHLHRSGSKLIGKVTSGLQIKRK--------------------
         Ailuropoda melanoleuca  (89) ALFWDLAQLFWDIYVDCVLSRNHGLGRRQLAGKE-E-KISAVHPQHTGRKQG-AYSQRLRTPFLKKKELIEGLISMYLHRSGSKFIGKVTSGLEIKRK--------------------
        Ursus arctos horribilis  (89) ALFWDLAQLFWDIYVDCVLSRNHGLGRRQLAGKE-E-KISAVHLQHTGRKQG-AYSQQLRTPFLKKKELIEGLISMYLHRSGSKVIGKVTSGLEIKRK--------------------
              Ursus maritimus  (89) ALFWDLAQLFWDIYVDCVLSRNHGLGRRQLAGKE-E-KISAVHLQHTGRKQG-AYSQQLRTPFLKKKELIEGLISMYLHRSGSKVIGKVTSGLEIKRK--------------------
             Canis lupus dingo  (89) ALFWDLAQLFWDIYVDCVLSRNHGLGRRQLAGKE-E-KISAVHPQHTGRKQG-AYSQQLRTPFLKKKELIEGLISMHLHKSGSKFIGKVTSGLEIKRK--------------------
        Canis lupus familiaris  (89) ALFWDLAQLFWDIYVDCVLSRNHGLGRRQLAGKE-E-KISAVHPQHTGRKQG-AYSQQLRTPFLKKKELIEGLISMHLHKSGSKFIGKVTSGLEIKRK--------------------
                Vulpes lagopus  (89) ALFWDLAQLFWDIYVDCVLSRNHGLGRRQLAGKE-E-KISAVHPQHTGRKQG-AYSQQLRTPFLKKKELIEGLISMRLHKSGSKFIGKVTSGLEIKRK--------------------
                 Vulpes vulpes  (89) ALFWDLAQLFWDIYVDCVLSRNHGLGRRQLAGKE-E-KISAVHPQHTGRKQG-AYSQQLRTPFLKKKELIEGLISMRLHKSGSKFIGKVTSGLEIKRK--------------------
            Callorhinus ursinus  (89) ALFWDLAQLFWDIYVDCVLSRNHGLGRRQLTGKE-E-KISAVHPQHTGRKQG-TYSQQLRTPFLKKKELIEGLISMHLHRSGSKFTGKVTSGLEIKRK--------------------
             Eumetopias jubatus  (89) ALFWDLAQLFWDIYVDCVLSRNHGLGRRQLTGKE-E-KISAVHPQHTGRKQG-TYSQQLRTPFLKKKELIEGLISMHLHRSGSKFIGKVASGLEIKRK--------------------
         Zalophus californianus  (89) ALFWDLAQLFWDIYVDCVLSRNHGLGRRQLTGKE-E-KISAVHPQHTGRKQG-TYSQQLRTPFLKKKELIEGLISMHLHRSGSKFIGKVASGLEIKRK--------------------
             Odobenus rosmarus  (89) ALFWDLAQLFWDIYVDCVLSRNHGLGRRQLTGKE-E-KISAVHPQHTGRKQG-TYSQQLRTPFLKKKELFEVLISMHLHRSGSKFIGKVASGLEIKRK--------------------
            Halichoerus grypus  (89) ALFWDLAQLFWDIYVDCVLSRNHGLGRRQLTGKE-E-KISAVHPQHTGRKQG-AYSQQLRTPFLKKKELIEGLISMHLHRSGSKFIGKVASGLEIKRK--------------------
                 Phoca vitulina  (89) ALFWDLAQLFWDIYVDCVLSRNHGLGRRQLTGKE-E-KISAVHPQHTGRKQG-AYSQQLRTPFLKKKELIEGLISMHLHRSGSKFIGKVASGLEIKRK--------------------
       Leptonychotes weddellii  (89) ALFWDLAQLFWDIYVDCVLSRNHGLGRRQLTGKE-E-KISAVHPQHTGIKQG-AYSQQLRTPFVKNKELIKGLMGMHLHRSGSKFIGKVASGLEIKRK--------------------
               Mirounga leonina  (89) ALFWDLAQLFWDIYVDCVLSRNHGLGRRQLTGKE-E-KISAVHPQHTGIKQG-AYSQQLRAPFVKKKELIEGLMGMHLHRSGSKFIGKVASGLEIKRK--------------------
     Neomonachus schauinslandi  (89) ALFWDLAQLFWDIYVDCVLSRNHGLGRRQLTGKE-E-KISAMHPQHTGRKQG-AYSQQLRTPFLKKKELIEGLMGMHLHRSGSKFIGKVASGLEIKRK--------------------
             Lontra canadensis  (89) ALFWDLAQLFWDIYVDCVLSRNHGLGRRQLSGKE-E-KISAAHPQHAGRNQG-AYSQQLRTPFLKKKELIEGLISMRLHRSGSKFIGKVTSGLEIKRK--------------------
```

```
                 Mustela erminea  (89) ALFWDLAQLFWDIYVDCVLSRNHGLGRRQLSGKE-E--VSAAHPQHAGRNQG-AYSQQLRTPFLKKKELIEGLISMHLHRSGSKFFGKVTSGLEIKRK--------------------
            Mustela putorius furo (89) ALFWDLAQLFWDIYVDCVLSRNHGLGRRQLSGKE-E--ISAAHPQHAGRNQG-AYSQQLRTPFLKKKELIEGLISMHLHRSGSKFFGKVTSGLEIKRK--------------------
                  Manis javanica  (89) ALFWDLAQLFWDIYVDCVLSRNHGLGRRQLAGEE-E-KISAVHPQDKGRKQG-AYSLQLRTPFLKKKELIEDLMNMYIRRSGSKFTGKVTSGLEIKRK--------------------
              Manis pentadactyla  (89) ALFWDLAQLFWDIYVDCVLSRNHGLGRRQLAGEE-E-KISAVHPHDKGRKQG-AYSLQLRTPFLKKKELIEDLINMYIRRSGSKFTGKVTSGLEIKRK--------------------
              Camelus dromedarius (89) ALFWDLAQLFWDIYVDCVLSRNHGLGRRQLAGEE-G-KISAVHPQHTEKKQG-AYSEQPRIPSLKKKELIEDLISMHIRRNGSKFTGKVTTGLEIKRK--------------------
                   Camelus ferus  (89) ALFWDLAQLFWDIYVDCVLSRNHGLGRRQLAGEE-G-KISAVHPQHTEKKQG-AYSEQPRIPSLKKKELIEDLISMHIRRNGSKFTGKVTTGLEIKRK--------------------
                  Vicugna pacos   (89) ALFWDLAQLFWDIYVDCVLSRNHGLGRRQLAGEE-G-KISAVHPQHTEKKQG-AYSEQPRIPSLKKKELIEDLISMHIRRNGSKFTGKVTTGLEIKRK--------------------
             Artibeus jamaicensis (89) ALFWDLAQLFWDIYVDCVLSRNHGLGRRQLAGEE-E-KISAGRPQHMRDKQG-AYSQQLRIPSLKKKKVIEDLLSVYVRRSGAKLTGKVSRGLEIKRK--------------------
            Phyllostomus discolor (89) ALFWDLAQLFWDIYVDCVLSRNHGLGRRQLAGEE-E-KISAGHPQHMREKQG-AYSQQLRIPSLKKKKVIEELLSVRMRRSGATLFGKASRGPEIKRK--------------------
               Desmodus rotundus  (89) ALFWDLAQLFWDIYVDCVVSRNHGLGRRQLAREE-E-KISAGRPQNMRKNQG-AYTQQRRIPSLKKKKVIEDLLSLHMRRSRATLPGKVSRGLEIKRK--------------------
            Sturnira hondurensis  (89) ALFWDLAQLFWDIYVDCVLSRNHGLGRRQLAGEK-E-KISAGHPQHMRGKQG-AYSQQLRIPSLKKKKVMEDLLNAHMRRSGAKLAGEVSRGLEIKTK--------------------
                 Eptesicus fuscus (89) ALFWDLAQLFWDIYVDCVLSRNHGLGRRQLAGEE-E-EISAVHPPHSGRKQG-AHSQQPRIRSLKKK-EIEDLLSLYIRRRGSKLIGKASSGLEMKRK--------------------
             Pipistrellus kuhlii  (89) ALFWDLAQLFWDIYVDCVLSRNHGLGRRQLTEEE-E-EISAAHPPHSGRKQG-AHSQQPRIPSLKKK-EIEDLLSLYIRRRRSKLIRKASSGLEMKRK--------------------
                 Myotis brandtii  (89) ALFWDLAQLFWDIYVDCVLSRNHGLGRRQLAGEE-E-EISAEHPPHSGRKQG-AHSQQPRILSLKKK-KIEDLLSLYIGRRESKSIGKASSGLEIKRK--------------------
                 Myotis lucifugus (89) ALFWDLAQLFWDIYVDCVLSRNHGLGRRQLAGEE-E-EISAVHPPHSGRKQG-AHSQQPRILSLKKK-KIEDLLSLYIRRRESKSIGKASSGLEIKRK--------------------
                  Myotis davidii  (89) ALFWDLAQLFWDIYVDCVLSRNHGLGRRQLAGEE-E-EILAVHPPHSGRKQG-AHSQQPRILSLKKK-EIEDLLSLYIRRRGSKSVGKASSGLEIERK--------------------
                   Myotis myotis  (89) ALFWDLAQLFWDIYVDCVLSRNHGLGRRQLAGEE-E-EMSAVHPPHSGRKQG-AHSQQPRILSLKKK-EIEDLLSLYIHRRGSKSIGKASSGLEIKRK--------------------
          Miniopterus natalensis  (89) ALFWDLAQLFWDIYVDCILSRNHGLGRRQLAGEE-E-RISAVHPQHTGRKQG-AYSQQLQIPSPKKK-ELIEDLLSLYVRS---KVGKASSGPEIKRKQI------------------
               Molossus molossus  (89) ALFWDLAQLFWDIYVDCILSRNHGLGRRQLAGEE-E-KISALHPEHTGRKQG-AYFQQLRISSLKKK-ELIDLLSLQMRRSGSKFIGKASSGLEIKRK--------------------
                 Pteropus alecto  (89) ALFWDLAQLFWDIYVDCVLSRNHGLGRRQLAGEE-E-KISTLHLQHSGKKQG-AYSQQLRIPSLKKKELIEDLLNIHMHRSGSKFFGKMTSGLKMKRK--------------------
              Pteropus giganteus  (89) ALFWDLAQLFWDIYVDCVLSRNHGLGRRQLAGEE-E-KISALHLQHSGKKQG-AYSQQLRIPSLKKKELIEDLLNMHMHRSGSKFFGKMTSGLKMKRK--------------------
               Pteropus vampyrus  (89) ALFWDLAQLFWDIYVDCVLSRNHGLGRRQLAGEE-E-KISALHLQHSGKKQG-AYSQQLRIPSLKKKELIEDLLNMHMHRSGSKFFGKMTSGLKMKRK--------------------
            Rousettus aegyptiacus (89) ALFWDLAQLFWDIYVDCVLSRNHGLGRRQLAGEE-E-KISALRLHHSGKKQG-AYFQQLRIPSLKKKESIEDLLNMHKHRSGSNFFGKITSGLKMKRK--------------------
             Hipposideros armiger (89) ALFWDLAQLFWDIYVDCVLSRNHGLGRRQLAGEE-E-KISAVHLQHTGRKQG-AYSQQLRIPFLKKKELIEDLLMMHVRRSGPKFAGKVTSGLEIKRK--------------------
        Rhinolophus ferrumequinum (89) ALFWELSQRFLDIYVDCVLSKNHGLGKRHLAGKE-E-KISAVHPQHTGRKQG-AYSQQLRIPSGKKKELIENLLSMHLHRSEPKFAGKVTSGLEKKRK--------------------
             Balaenoptera musculus (89) ALFWDLAQLFWDIYVDCVLSRNHGLGRRQLAGEE-E-KISAVHPQHTGRKKG-VYPQQPRIPSLKKKELIEDLISMHAHRSGSKFIGKVTNVLEIKRK--------------------
                 Physeter catodon (89) ALFWDLAQLFWDIYVDCVLSRNHGLGRRQLAGAE-E-KISAVHPQHTGREKG-VYPQQPRIPSLKKKELIEDLISMHAHKSGSKFIGKVTSSLEIKRK--------------------
            Delphinapterus leucas (89) ALFWDLAQLFWDIYVDCVLSRNHGLGRRQLAGEE-E-KISAVHPQHTGRKKG-VYPQQPRIPSLKKKELIEGLINMHVHRSGSKFIGKVTSSLEIKRK--------------------
               Monodon monoceros  (89) ALFWDLAQLFWDIYVDCVLSRNHGLGRRQLAGEE-E-KISAVHPQHTGRKKG-VYPQQPRIPSLKKKELIEGLINMHVHRSGSKFIGKVTSSLEIKRK--------------------
        Neophocaena asiaeorientalis (89) ALFWDLAQLFWDIYVDCVLSRNHGLGRRQLAGEE-E-KISAVHPQHTGRKKG-VYPQQPRITSLKKKELIEGLINMHVHRSGSKFIGKVTSSLEIKRK--------------------
              Globicephala melas (89) ALFWDLAQLFWDIYVDCVLSRNHGLGRRQLAGEE-E-KISAVHPQHAGRKKG-VYPQQPRIPSLKKKELIEGLINMHVHRSGSKFIGKVTSSLEIKRK--------------------
        Lagenorhynchus obliquidens (89) ALFWDLAQLFWDIYVDCVLSRNHGLGRRQLAGEE-E-KISAVHPQHAGRKKG-VYPQQPRIPSLKKKELIEGLINMHVHRSGSKFIGKVTSSLEIKRK--------------------
                    Orcinus orca  (89) ALFWDLAQLFWDIYVDCVLSRNHGLGRRQLAGEE-E-KISAVPPQHAGRKKG-VYPQQPRIPSLKKKELIEGLINMHVHRSGSKFIGKVTSSLEIKRK--------------------
              Tursiops truncatus  (89) ALFWDLAQLFWDIYVDCVLSRNHGLGRRQLAGEE-E-KISAVPPQHAGRKKG-VYPQQPRIPSLKKKELIEGLINMHVHRSGSKFIGKVTSSLEIKRK--------------------
                  Phocoena sinus  (89) ALFWDLAQLFWDIYVDCVLSRNHGLGRRQLAGEE-E-KISAVHPQHTGRKKG-IYPQQPRIPSLKKKELIEGLINMHVHRSGSKFIGKVTSSLEIKRK--------------------
               Bison bison bison  (89) ALFWHLAQLFWDIYVDCVLSRNHGLGRRQLTGEE-V-KISAVHPQQTGRKKD-AYSQQPRTPSLKKKELIEDLISLHVHRSRSKFIGKATSGLEIKRK--------------------
                      Bos taurus  (89) ALFWHLAQLFWDIYVDCVLSRNHGLGRRQLTGEE-V-KISAVHPQQTGRKKG-AYSQQPRTPSLKKKELIEDLISLHVHRSRSKFIGKATSGLEIKRK--------------------
                 Bubalus bubalis  (89) ALFWDLAQLFWDIYVDCVLSRNHGLGRRQLTGEE-V-KISAVHPQHTGRKKD-AYSQQPRTPSLKKKELIEDLISMHVHRSRSKFIGKATSGLEIKRK--------------------
                    Capra hircus  (89) ALFWDLAQLFWDIYVDCVLSRNHGLGRRQLTGEE-V-KISAVHPKHTGRKKG-AYSQQPRNPSLKKKELIEDLISMHVHRSRSKFIGKVTSGLETKRK--------------------
                    Oryx dammah   (89) ALFWDLAQLFWDIYVDCVLSRNHGLGRRQLTGEE-V-KISAVHPQHTGRKKG-AYSQQPRNPSLKKKELIEDLISMHVHRSRSKFIGKATSGLRIKRK--------------------
                      Ovis aries  (89) ALFWDLAQLFWDIYVDCVLSRNHGLGRRQLTGEE-V-KISAVHPQHTGRKKG-AYSQQPRNPSLKKKELIEDLISMHVHRSRSKFIGKATSGLEIKRK--------------------
    Odocoileus virginianus texanus (89) ALFWDLAQLFWDIYVDCVLSRNHGLGRRQLTGEE-V-KMSAVHPQHTGRKKG-AYSQQPRTPSLKKKELIEDLISMHVHRSRSKFIGKVTSGLEIKRK--------------------
                      Sus scrofa  (89) ALFWDLAQLFWDIYVDCVLSRNHGLGRRQLAGEE-E------HPQHTGRKQG-TYSQQLRIPSLKKKEMIEDLISTHIHRSGPKFIGKITSDREIKRKEI------------------
                    Equus asinus  (89) ALFWDLAQLFLDIYVDCVLSRNHGLGRRHLAGEE-E-KISAVHPQHTKRKQG-TYSQQLRAPALKKKQLVEELINTHVRRSGSKFLGKVTSSLEIKRK--------------------
                  Equus caballus  (89) ALFWDLAQLFLDIYVDCVLSRNHGLGRRQLAGEE-E-KISAVHPQHTRRKQG-TYSQQLRAPALKKKQLVEELINTHVRRSGSKFLGKVTSGLEIERK--------------------
```

**Fig. S2.** Amino acid sequence alignment of FAM237A orthologs from fish to mammals.

#### FAM237A-Intein-6×His in pET vector

```
 NsiI
ATG CAT AGC CAG ACC GAC CTG CTG GCT CTT AGT CAA GCG GAT CCG CAG TGC TGG GAA TCC TCT TCA GTG CTG CTC CTG GAA ATG TGG AAA CCG CGC GTT TCG AAC
TAC GTA TCG GTC TGG CTG GAC GAC CGA GAA TCA GTT CGC CTA GGC GTC ACG ACC CTT AGG AGA AGT CAC GAC GAG GAC CTT TAC ACC TTT GGC GCG CAA AGC TTG
 M   H   S   Q   T   D   L   L   A   L   S   Q   A   D   P   Q   C   W   E   S   S   S   V   L   L   L   E   M   W   K   P   R   V   S   N

ACT GTC AGC GGC TTC TGG GAT TTT ATG ATC TAC CTG AAG TCC TCT GAG AAT TTG AAA CAC GGT GCA CTG TTT TGG GAT CTG GCC CAG CTC TTC TGG GAC ATT TAT
TGA CAG TCG CCG AAG ACC CTA AAA TAC TAG ATG GAC TTC AGG AGA CTC TTA AAC TTT GTG CCA CGT GAC AAA ACC CTA GAC CGG GTC GAG AAG ACC CTG TAA ATA
 T   V   S   G   F   W   D   F   M   I   Y   L   K   S   S   E   N   L   K   H   G   A   L   F   W   D   L   A   Q   L   F   W   D   I   Y

GTA GAC TGT GTG CTT AGC CGT AAC CAT GGC TTA TGT ATT TGC GGT GAT GCT CTG GTG GCC CTG CCG GAA GGC GAA AGC GTT CGT ATC GCA GAC ATT GTC CCA GGT
CAT CTG ACA CAC GAA TCG GCA TTG GTA CCG AAT ACA TAA ACG CCA CTA CGA GAC CAC CGG GAC GGC CTT CCG CTT TCG CAA GCA TAG CGT CTG TAA CAG GGT CCA
 V   D   C   V   L   S   R   N   H   G   L   C   I   C   G   D   A   L   V   A   L   P   E   G   E   S   V   R   I   A   D   I   V   P   G

GCT CGC CCG AAC AGT GAT AAT GCG ATC GAC CTG AAA GTA CTC GAT CGT CAC GGC AAC CCA GTG TTA GCG GAT CGT TTG TTT CAT TCT GGT GAG CAT CCT GTT TAT
CGA GCG GGC TTG TCA CTA TTA CGC TAG CTG GAC TTT CAT GAG CTA GCA GTG CCG TTG GGT CAC AAT CGC CTA GCA AAC AAA GTA AGA CCA CTC GTA GGA CAA ATA
 A   R   P   N   S   D   N   A   I   D   L   K   V   L   D   R   H   G   N   P   V   L   A   D   R   L   F   H   S   G   E   H   P   V   Y

ACC GTC CGC ACG GTA GAA GGT CTG CGT GTG ACT GGC ACC GCC AAC CAC CCG CTG CTG TGC TTA GTT GAT GTC GCA GGT GTA CCG ACC CTG TTG TGG AAG TTA ATT
TGG CAG GCG TGC CAT CTT CCA GAC GCA CAC TGA CCG TGG CGG TTG GTG GGC GAC GAC ACG AAT CAA CTA CAG CGT CCA CAT GGC TGG GAC AAC ACC TTC AAT TAA
 T   V   R   T   V   E   G   L   R   V   T   G   T   A   N   H   P   L   L   C   L   V   D   V   A   G   V   P   T   L   L   W   K   L   I

GAT GAG ATC AAA CCA GGC GAC TAC GCT GTG ATT CAG CGT TCC GCG TTC TCA GTT GAT TGT GCT GGC TTT GCC CGC GGT AAA CCG GAG TTC GCA CCT ACC ACT TAT
CTA CTC TAG TTT GGT CCG CTG ATG CGA CAC TAA GTC GCA AGG CGC AAG AGT CAA CTA ACA CGA CCG AAA CGG GCG CCA TTT GGC CTC AAG CGT GGA TGG TGA ATA
 D   E   I   K   P   G   D   Y   A   V   I   Q   R   S   A   F   S   V   D   C   A   G   F   A   R   G   K   P   E   F   A   P   T   T   Y

ACG GTG GGC GTT CCG GGT CTG GTC CGT TTC CTG GAG GCG CAC CAT CGC GAT CCA GAC GCA CAG GCT ATC GCG GAT GAA CTG ACG GAT GGC CGT TTT TAT TAC GCC
TGC CAC CCG CAA GGC CCA GAC CAG GCA AAG GAC CTC CGC GTG GTA GCG CTA GGT CTG CGT GTC CGA TAG CGC CTA CTT GAC TGC CTA CCG GCA AAA ATA ATG CGG
 T   V   G   V   P   G   L   V   R   F   L   E   A   H   H   R   D   P   D   A   Q   A   I   A   D   E   L   T   D   G   R   F   Y   Y   A

AAA GTG GCT AGC GTT ACC GAC GCG GGT GTG CAA CCG GTC TAT TCG TTA CGC GTT GAT ACG GCA GAC CAC GCG TTC ATC ACG AAC GGC TTC GTT TCT CAT GCG GCG
TTT CAC CGA TCG CAA TGG CTG CGC CCA CAC GTT GGC CAG ATA AGC AAT GCG CAA CTA TGC CGT CTG GTG CGC AAG TAG TGC TTG CCG AAG CAA AGA GTA CGC CGC
 K   V   A   S   V   T   D   A   G   V   Q   P   V   Y   S   L   R   V   D   T   A   D   H   A   F   I   T   N   G   F   V   S   H   A   A

NotI
GCC GCA CTC GAG CAC CAC CAC CAC CAC CAC TGA
CGG CGT GAG CTC GTG GTG GTG GTG GTG GTG ACT
 A   A   L   E   H   H   H   H   H   H   *
```

#### SmBiT-FAM237A-Intein-6×His in pET vector

```
NsiI
ATG CAT GTG ACC GGC TAC CGT CTG TTT GAA GAA ATT CTG GGC GGC AGC GGT GGT GGC CAT AGC CAG ACC GAC CTG CTG GCT CTT AGT CAA GCG GAT CCG CAG TGC
TAC GTA CAC TGG CCG ATG GCA GAC AAA CTT CTT TAA GAC CCG CCG TCG CCA CCA CCG GTA TCG GTC TGG CTG GAC GAC CGA GAA TCA GTT CGC CTA GGC GTC ACG
 M   H   V   T   G   Y   R   L   F   E   E   I   L   G   G   S   G   G   G   H   S   Q   T   D   L   L   A   L   S   Q   A   D   P   Q   C

TGG GAA TCC TCT TCA GTG CTG CTC CTG GAA ATG TGG AAA CCG CGC GTT TCG AAC ACT GTC AGC GGC TTC TGG GAT TTT ATG ATC TAC CTG AAG TCC TCT GAG AAT
ACC CTT AGG AGA AGT CAC GAC GAG GAC CTT TAC ACC TTT GGC GCG CAA AGC TTG TGA CAG TCG CCG AAG ACC CTA AAA TAC TAG ATG GAC TTC AGG AGA CTC TTA
 W   E   S   S   S   V   L   L   L   E   M   W   K   P   R   V   S   N   T   V   S   G   F   W   D   F   M   I   Y   L   K   S   S   E   N

TTG AAA CAC GGT GCA CTG TTT TGG GAT CTG GCC CAG CTC TTC TGG GAC ATT TAT GTA GAC TGT GTG CTT AGC CGT AAC CAT GGC TTA TGT ATT TGC GGT GAT GCT
AAC TTT GTG CCA CGT GAC AAA ACC CTA GAC CGG GTC GAG AAG ACC CTG TAA ATA CAT CTG ACA CAC GAA TCG GCA TTG GTA CCG AAT ACA TAA ACG CCA CTA CGA
 L   K   H   G   A   L   F   W   D   L   A   Q   L   F   W   D   I   Y   V   D   C   V   L   S   R   N   H   G   L   C   I   C   G   D   A

CTG GTG GCC CTG CCG GAA GGC GAA AGC GTT CGT ATC GCA GAC ATT GTC CCA GGT GCT CGC CCG AAC AGT GAT AAT GCG ATC GAC CTG AAA GTA CTC GAT CGT CAC
GAC CAC CGG GAC GGC CTT CCG CTT TCG CAA GCA TAG CGT CTG TAA CAG GGT CCA CGA GCG GGC TTG TCA CTA TTA CGC TAG CTG GAC TTT CAT GAG CTA GCA GTG
 L   V   A   L   P   E   G   E   S   V   R   I   A   D   I   V   P   G   A   R   P   N   S   D   N   A   I   D   L   K   V   L   D   R   H

GGC AAC CCA GTG TTA GCG GAT CGT TTG TTT CAT TCT GGT GAG CAT CCT GTT TAT ACC GTC CGC ACG GTA GAA GGT CTG CGT GTG ACT GGC ACC GCC AAC CAC CCG
CCG TTG GGT CAC AAT CGC CTA GCA AAC AAA GTA AGA CCA CTC GTA GGA CAA ATA TGG CAG GCG TGC CAT CTT CCA GAC GCA CAC TGA CCG TGG CGG TTG GTG GGC
 G   N   P   V   L   A   D   R   L   F   H   S   G   E   H   P   V   Y   T   V   R   T   V   E   G   L   R   V   T   G   T   A   N   H   P

CTG CTG TGC TTA GTT GAT GTC GCA GGT GTA CCG ACC CTG TTG TGG AAG TTA ATT GAT GAG ATC AAA CCA GGC GAC TAC GCT GTG ATT CAG CGT TCC GCG TTC TCA
GAC GAC ACG AAT CAA CTA CAG CGT CCA CAT GGC TGG GAC AAC ACC TTC AAT TAA CTA CTC TAG TTT GGT CCG CTG ATG CGA CAC TAA GTC GCA AGG CGC AAG AGT
 L   L   C   L   V   D   V   A   G   V   P   T   L   L   W   K   L   I   D   E   I   K   P   G   D   Y   A   V   I   Q   R   S   A   F   S

GTT GAT TGT GCT GGC TTT GCC CGC GGT AAA CCG GAG TTC GCA CCT ACC ACT TAT ACG GTG GGC GTT CCG GGT CTG GTC CGT TTC CTG GAG GCG CAC CAT CGC GAT
CAA CTA ACA CGA CCG AAA CGG GCG CCA TTT GGC CTC AAG CGT GGA TGG TGA ATA TGC CAC CCG CAA GGC CCA GAC CAG GCA AAG GAC CTC CGC GTG GTA GCG CTA
 V   D   C   A   G   F   A   R   G   K   P   E   F   A   P   T   T   Y   T   V   G   V   P   G   L   V   R   F   L   E   A   H   H   R   D

CCA GAC GCA CAG GCT ATC GCG GAT GAA CTG ACG GAT GGC CGT TTT TAT TAC GCC AAA GTG GCT AGC GTT ACC GAC GCG GGT GTG CAA CCG GTC TAT TCG TTA CGC
GGT CTG CGT GTC CGA TAG CGC CTA CTT GAC TGC CTA CCG GCA AAA ATA ATG CGG TTT CAC CGA TCG CAA TGG CTG CGC CCA CAC GTT GGC CAG ATA AGC AAT GCG
 P   D   A   Q   A   I   A   D   E   L   T   D   G   R   F   Y   Y   A   K   V   A   S   V   T   D   A   G   V   Q   P   V   Y   S   L   R

                                                                     NotI
GTT GAT ACG GCA GAC CAC GCG TTC ATC ACG AAC GGC TTC GTT TCT CAT GCG GCG GCC GCA CTC GAG CAC CAC CAC CAC CAC CAC TGA
CAA CTA TGC CGT CTG GTG CGC AAG TAG TGC TTG CCG AAG CAA AGA GTA CGC CGC CGG CGT GAG CTC GTG GTG GTG GTG GTG GTG ACT
 V   D   T   A   D   H   A   F   I   T   N   G   F   V   S   H   A   A   A   A   L   E   H   H   H   H   H   H   *
```

**Fig. S3.** The nucleotide and amino acid sequences of human FAM237A constructs overexpressed in *E. coli*. The amino acid sequence of mature human FAM237A is shown in red, that of intein in yellow, and that of SmBiT in blue. The restriction enzyme cleavage sites for cloning are shaded.

### Human GPR83 in pcDNA6 vector

```
NheI
GCT AGC ATG GTC CCT CAC CTC TTG CTG CTC TGT CTC CTC CCC TTG GTG CGA GCC ACC GAG CCC CAC GAG GGC CGG GCC GAC GAG CAG AGC GCG GAG GCG GCC CTG
CGA TCG TAC CAG GGA GTG GAG AAC GAC GAG ACA GAG GAG GGG AAC CAC GCT CGG TGG CTC GGG GTG CTC CCG GCC CGG CTG CTC GTC TCG CGC CTC CGC CGG GAC
         M   V   P   H   L   L   L   L   C   L   L   P   L   V   R   A   T   E   P   H   E   G   R   A   D   E   Q   S   A   E   A   A   L

GCC GTG CCC AAT GCC TCG CAC TTC TTC TCT TGG AAC AAC TAC ACC TTC TCC GAC TGG CAG AAC TTT GTG GGC AGG AGG CGC TAC GGC GCT GAG TCC CAG AAC CCC
CGG CAC GGG TTA CGG AGC GTG AAG AAG AGA ACC TTG TTG ATG TGG AAG AGG CTG ACC GTC TTG AAA CAC CCG TCC TCC GCG ATG CCG CGA CTC AGG GTC TTG GGG
 A   V   P   N   A   S   H   F   F   S   W   N   N   Y   T   F   S   D   W   Q   N   F   V   G   R   R   R   Y   G   A   E   S   Q   N   P

ACG GTG AAA GCC CTG CTC ATT GTG GCT TAC TCC TTC ATC ATT GTC TTC TCA CTC TTT GGC AAC GTC CTG GTC TGT CAT GTC ATC TTC AAG AAC CAG CGA ATG CAC
TGC CAC TTT CGG GAC GAG TAA CAC CGA ATG AGG AAG TAG TAA CAG AAG AGT GAG AAA CCG TTG CAG GAC CAG ACA GTA CAG TAG AAG TTC TTG GTC GCT TAC GTG
 T   V   K   A   L   L   I   V   A   Y   S   F   I   I   V   F   S   L   F   G   N   V   L   V   C   H   V   I   F   K   N   Q   R   M   H

TCG GCC ACC AGC CTC TTC ATC GTC AAC CTG GCA GTT GCC GAC ATA ATG ATC ACG CTG CTC AAC ACC CCC TTC ACT TTG GTT CGC TTT GTG AAC AGC ACA TGG ATA
AGC CGG TGG TCG GAG AAG TAG CAG TTG GAC CGT CAA CGG CTG TAT TAC TAG TGC GAC GAG TTG TGG GGG AAG TGA AAC CAA GCG AAA CAC TTG TCG TGT ACC TAT
 S   A   T   S   L   F   I   V   N   L   A   V   A   D   I   M   I   T   L   L   N   T   P   F   T   L   V   R   F   V   N   S   T   W   I

TTT GGG AAG GGC ATG TGC CAT GTC AGC CGC TTT GCC CAG TAC TGC TCA CTG CAC GTC TCA GCA CTG ACA CTG ACA GCC ATT GCG GTG GAT CGC CAC CAG GTC ATC
AAA CCC TTC CCG TAC ACG GTA CAG TCG GCG AAA CGG GTC ATG ACG AGT GAC GTG CAG AGT CGT GAC TGT GAC TGT CGG TAA CGC CAC CTA GCG GTG GTC CAG TAG
 F   G   K   G   M   C   H   V   S   R   F   A   Q   Y   C   S   L   H   V   S   A   L   T   L   T   A   I   A   V   D   R   H   Q   V   I

ATG CAC CCC TTG AAA CCC CGG ATC TCA ATC ACA AAG GGT GTC ATC TAC ATC GCT GTC ATC TGG ACC ATG GCT ACG TTC TTT TCA CTC CCA CAT GCT ATC TGC CAG
TAC GTG GGG AAC TTT GGG GCC TAG AGT TAG TGT TTC CCA CAG TAG ATG TAG CGA CAG TAG ACC TGG TAC CGA TGC AAG AAA AGT GAG GGT GTA CGA TAG ACG GTC
 M   H   P   L   K   P   R   I   S   I   T   K   G   V   I   Y   I   A   V   I   W   T   M   A   T   F   F   S   L   P   H   A   I   C   Q

AAA TTA TTT ACC TTC AAA TAC AGT GAG GAC ATT GTG CGC TCC CTC TGC CTG CCA GAC TTC CCT GAG CCA GCT GAC CTC TTC TGG AAG TAC CTG GAC TTG GCC ACC
TTT AAT AAA TGG AAG TTT ATG TCA CTC CTG TAA CAC GCG AGG GAG ACG GAC GGT CTG AAG GGA CTC GGT CGA CTG GAG AAG ACC TTC ATG GAC CTG AAC CGG TGG
 K   L   F   T   F   K   Y   S   E   D   I   V   R   S   L   C   L   P   D   F   P   E   P   A   D   L   F   W   K   Y   L   D   L   A   T

TTC ATC CTG CTC TAC ATC CTG CCC CTC CTC ATC ATC TCT GTG GCC TAC GCT CGT GTG GCC AAG AAA CTG TGG CTG TGT AAT ATG ATT GGC GAT GTG ACC ACA GAG
AAG TAG GAC GAG ATG TAG GAC GGG GAG GAG TAG TAG AGA CAC CGG ATG CGA GCA CAC CGG TTC TTT GAC ACC GAC ACA TTA TAC TAA CCG CTA CAC TGG TGT CTC
 F   I   L   L   Y   I   L   P   L   L   I   I   S   V   A   Y   A   R   V   A   K   K   L   W   L   C   N   M   I   G   D   V   T   T   E

CAG TAC TTT GCC CTG CGG CGC AAA AAG AAG AAG ACC ATC AAG ATG TTG ATG CTG GTG GTA GTC CTC TTT GCC CTC TGC TGG TTC CCC CTC AAC TGC TAC GTC CTC
GTC ATG AAA CGG GAC GCC GCG TTT TTC TTC TTC TGG TAG TTC TAC AAC TAC GAC CAC CAT CAG GAG AAA CGG GAG ACG ACC AAG GGG GAG TTG ACG ATG CAG GAG
 Q   Y   F   A   L   R   R   K   K   K   K   T   I   K   M   L   M   L   V   V   V   L   F   A   L   C   W   F   P   L   N   C   Y   V   L

CTC CTG TCC AGC AAG GTC ATC CGC ACC AAC AAT GCC CTC TAC TTT GCC TTC CAC TGG TTT GCC ATG AGC AGC ACC TGC TAT AAC CCC TTC ATA TAC TGC TGG CTG
GAG GAC AGG TCG TTC CAG TAG GCG TGG TTG TTA CGG GAG ATG AAA CGG AAG GTG ACC AAA CGG TAC TCG TCG TGG ACG ATA TTG GGG AAG TAT ATG ACG ACC GAC
 L   L   S   S   K   V   I   R   T   N   N   A   L   Y   F   A   F   H   W   F   A   M   S   S   T   C   Y   N   P   F   I   Y   C   W   L

AAC GAG AAC TTC AGG ATT GAG CTA AAG GCA TTA CTG AGC ATG TGT CAA AGA CCT CCC AAG CCT CAG GAG GAC AGG CCA CCC TCC CCA GTT CCT TCC TTC AGG GTG
TTG CTC TTG AAG TCC TAA CTC GAT TTC CGT AAT GAC TCG TAC ACA GTT TCT GGA GGG TTC GGA GTC CTC CTG TCC GGT GGG AGG GGT CAA GGA AGG AAG TCC CAC
```

N E N F R I E L K A L L S M C Q R P P K P Q E D R P P S P V P S F R V

GCC TGG ACA GAG AAG AAT GAT GGC CAG AGG GCT CCC CTT GCC AAT AAC CTC CTG CCC ACC TCC CAA CTC CAG TCT GGG AAG ACA GAC CTG TCA TCT GTG GAA CCC
CGG ACC TGT CTC TTC TTA CTA CCG GTC TCC CGA GGG GAA CGG TTA TTG GAG GAC GGG TGG AGG GTT GAG GTC AGA CCC TTC TGT CTG GAC AGT AGA CAC CTT GGG
A W T E K N D G Q R A P L A N N L L P T S Q L Q S G K T D L S S V E P

AgeI
ATT GTG ACG ATG AGT CCA CCG GTC TGA TAA
TAA CAC TGC TAC TCA GGT GGC CAG ACT ATT
I V T M S P P V * *

#### sLgBiT-GPR83 in pcDNA6 vector

NheI
GCT AGC ATG AAC TCC TTC TCC ACA AGC GCC TTC GGT CCA GTT GCC TTC TCC CTG GGC CTG CTC CTG GTG TTG CCT GCT GCC TTC CCT GCC CCA GTC TTC ACA CTC
CGA TCG TAC TTG AGG AAG AGG TGT TCG CGG AAG CCA GGT CAA CGG AAG AGG GAC CCG GAC GAG GAC CAC AAC GGA CGA CGG AAG GGA CGG GGT CAG AAG TGT GAG
M N S F S T S A F G P V A F S L G L L L V L P A A F P A P V F T L

GAA GAT TTC GTT GGG GAC TGG GAA CAG ACA GCC GCC TAC AAC CTG GAC CAA GTC CTT GAA CAG GGA GGT GTG TCC AGT TTG CTG CAG AAT CTC GCC GTG TCC GTA
CTT CTA AAG CAA CCC CTG ACC CTT GTC TGT CGG CGG ATG TTG GAC CTG GTT CAG GAA CTT GTC CCT CCA CAC AGG TCA AAC GAC GTC TTA GAG CGG CAC AGG CAT
E D F V G D W E Q T A A Y N L D Q V L E Q G G V S S L L Q N L A V S V

ACT CCG ATC CAA AGG ATT GTC CGG AGC GGT GAA AAT GCC CTG AAG ATC GAC ATC CAT GTC ATC ATC CCG TAT GAA GGT CTG AGC GCC GAC CAA ATG GCC CAG ATC
TGA GGC TAG GTT TCC TAA CAG GCC TCG CCA CTT TTA CGG GAC TTC TAG CTG TAG GTA CAG TAG TAG GGC ATA CTT CCA GAC TCG CGG CTG GTT TAC CGG GTC TAG
T P I Q R I V R S G E N A L K I D I H V I I P Y E G L S A D Q M A Q I

GAA GAG GTG TTT AAG GTG GTG TAC CCT GTG GAT GAT CAT CAC TTT AAG GTG ATC CTG CCC TAT GGC ACA CTG GTA ATC GAC GGG GTT ACG CCG AAC ATG CTG AAC
CTT CTC CAC AAA TTC CAC CAC ATG GGA CAC CTA CTA GTA GTG AAA TTC CAC TAG GAC GGG ATA CCG TGT GAC CAT TAG CTG CCC CAA TGC GGC TTG TAC GAC TTG
E E V F K V V Y P V D D H H F K V I L P Y G T L V I D G V T P N M L N

TAT TTC GGA CGG CCG TAT GAA GGC ATC GCC GTG TTC GAC GGC AAA AAG ATC ACT GTA ACA GGG ACC CTG TGG AAC GGC AAC AAA ATT ATC GAC GAG CGC CTG ATC
ATA AAG CCT GCC GGC ATA CTT CCG TAG CGG CAC AAG CTG CCG TTT TTC TAG TGA CAT TGT CCC TGG GAC ACC TTG CCG TTG TTT TAA TAG CTG CTC GCG GAC TAG
Y F G R P Y E G I A V F D G K K I T V T G T L W N G N K I I D E R L I

KpnI
ACC CCC GAC GGC TCC ATG CTG TTC CGA GTA ACC ATC AAC AGT GGT GGC GGC TCT GGT GGT GGC AGC GGC GGT GGT ACC ACC GAG CCC CAC GAG GGC CGG GCC GAC
TGG GGG CTG CCG AGG TAC GAC AAG GCT CAT TGG TAG TTG TCA CCA CCG CCG AGA CCA CCA CCG TCG CCG CCA CCA TGG TGG CTC GGG GTG CTC CCG GCC CGG CTG
T P D G S M L F R V T I N S G G G S G G G S G G G T T E P H E G R A D

GAG CAG AGC GCG GAG GCG GCC CTG GCC GTG CCC AAT GCC TCG CAC TTC TTC TCT TGG AAC AAC TAC ACC TTC TCC GAC TGG CAG AAC TTT GTG GGC AGG AGG CGC
CTC GTC TCG CGC CTC CGC CGG GAC CGG CAC GGG TTA CGG AGC GTG AAG AAG AGA ACC TTG TTG ATG TGG AAG AGG CTG ACC GTC TTG AAA CAC CCG TCC TCC GCG
E Q S A E A A L A V P N A S H F F S W N N Y T F S D W Q N F V G R R R

TAC GGC GCT GAG TCC CAG AAC CCC ACG GTG AAA GCC CTG CTC ATT GTG GCT TAC TCC TTC ATC ATT GTC TTC TCA CTC TTT GGC AAC GTC CTG GTC TGT CAT GTC

ATG CCG CGA CTC AGG GTC TTG GGG TGC CAC TTT CGG GAC GAG TAA CAC CGA ATG AGG AAG TAG TAA CAG AAG AGT GAG AAA CCG TTG CAG GAC CAG ACA GTA CAG
Y G A E S Q N P T V K A L L I V A Y S F I I V F S L F G N V L V C H V

ATC TTC AAG AAC CAG CGA ATG CAC TCG GCC ACC AGC CTC TTC ATC GTC AAC CTG GCA GTT GCC GAC ATA ATG ATC ACG CTG CTC AAC ACC CCC TTC ACT TTG GTT
TAG AAG TTC TTG GTC GCT TAC GTG AGC CGG TGG TCG GAG AAG TAG CAG TTG GAC CGT CAA CGG CTG TAT TAC TAG TGC GAC GAG TTG TGG GGG AAG TGA AAC CAA
I F K N Q R M H S A T S L F I V N L A V A D I M I T L L N T P F T L V

CGC TTT GTG AAC AGC ACA TGG ATA TTT GGG AAG GGC ATG TGC CAT GTC AGC CGC TTT GCC CAG TAC TGC TCA CTG CAC GTC TCA GCA CTG ACA CTG ACA GCC ATT
GCG AAA CAC TTG TCG TGT ACC TAT AAA CCC TTC CCG TAC ACG GTA CAG TCG GCG AAA CGG GTC ATG ACG AGT GAC GTG CAG AGT CGT GAC TGT GAC TGT CGG TAA
R F V N S T W I F G K G M C H V S R F A Q Y C S L H V S A L T L T A I

GCG GTG GAT CGC CAC CAG GTC ATC ATG CAC CCC TTG AAA CCC CGG ATC TCA ATC ACA AAG GGT GTC ATC TAC ATC GCT GTC ATC TGG ACC ATG GCT ACG TTC TTT
CGC CAC CTA GCG GTG GTC CAG TAG TAC GTG GGG AAC TTT GGG GCC TAG AGT TAG TGT TTC CCA CAG TAG ATG TAG CGA CAG TAG ACC TGG TAC CGA TGC AAG AAA
A V D R H Q V I M H P L K P R I S I T K G V I Y I A V I W T M A T F F

TCA CTC CCA CAT GCT ATC TGC CAG AAA TTA TTT ACC TTC AAA TAC AGT GAG GAC ATT GTG CGC TCC CTC TGC CTG CCA GAC TTC CCT GAG CCA GCT GAC CTC TTC
AGT GAG GGT GTA CGA TAG ACG GTC TTT AAT AAA TGG AAG TTT ATG TCA CTC CTG TAA CAC GCG AGG GAG ACG GAC GGT CTG AAG GGA CTC GGT CGA CTG GAG AAG
S L P H A I C Q K L F T F K Y S E D I V R S L C L P D F P E P A D L F

TGG AAG TAC CTG GAC TTG GCC ACC TTC ATC CTG CTC TAC ATC CTG CCC CTC CTC ATC ATC TCT GTG GCC TAC GCT CGT GTG GCC AAG AAA CTG TGG CTG TGT AAT
ACC TTC ATG GAC CTG AAC CGG TGG AAG TAG GAC GAG ATG TAG GAC GGG GAG GAG TAG TAG AGA CAC CGG ATG CGA GCA CAC CGG TTC TTT GAC ACC GAC ACA TTA
W K Y L D L A T F I L L Y I L P L L I I S V A Y A R V A K K L W L C N

ATG ATT GGC GAT GTG ACC ACA GAG CAG TAC TTT GCC CTG CGG CGC AAA AAG AAG AAG ACC ATC AAG ATG TTG ATG CTG GTG GTA GTC CTC TTT GCC CTC TGC TGG
TAC TAA CCG CTA CAC TGG TGT CTC GTC ATG AAA CGG GAC GCC GCG TTT TTC TTC TTC TGG TAG TTC TAC AAC TAC GAC CAC CAT CAG GAG AAA CGG GAG ACG ACC
M I G D V T T E Q Y F A L R R K K K K T I K M L M L V V V L F A L C W

TTC CCC CTC AAC TGC TAC GTC CTC CTC CTG TCC AGC AAG GTC ATC CGC ACC AAC AAT GCC CTC TAC TTT GCC TTC CAC TGG TTT GCC ATG AGC AGC ACC TGC TAT
AAG GGG GAG TTG ACG ATG CAG GAG GAG GAC AGG TCG TTC CAG TAG GCG TGG TTG TTA CGG GAG ATG AAA CGG AAG GTG ACC AAA CGG TAC TCG TCG TGG ACG ATA
F P L N C Y V L L L S S K V I R T N N A L Y F A F H W F A M S S T C Y

AAC CCC TTC ATA TAC TGC TGG CTG AAC GAG AAC TTC AGG ATT GAG CTA AAG GCA TTA CTG AGC ATG TGT CAA AGA CCT CCC AAG CCT CAG GAG GAC AGG CCA CCC
TTG GGG AAG TAT ATG ACG ACC GAC TTG CTC TTG AAG TCC TAA CTC GAT TTC CGT AAT GAC TCG TAC ACA GTT TCT GGA GGG TTC GGA GTC CTC CTG TCC GGT GGG
N P F I Y C W L N E N F R I E L K A L L S M C Q R P P K P Q E D R P P

TCC CCA GTT CCT TCC TTC AGG GTG GCC TGG ACA GAG AAG AAT GAT GGC CAG AGG GCC CCC CTT GCC AAT AAC CTC CTG CCC ACC TCC CAA CTC CAG TCT GGG AAG
AGG GGT CAA GGA AGG AAG TCC CAC CGG ACC TGT CTC TTC TTA CTA CCG GTC TCC CGG GGG GAA CGG TTA TTG GAG GAC GGG TGG AGG GTT GAG GTC AGA CCC TTC
S P V P S F R V A W T E K N D G Q R A P L A N N L L P T S Q L Q S G K

AgeI
ACA GAC CTG TCA TCT GTG GAA CCC ATT GTG ACG ATG AGT TAG ACC GGT
TGT CTG GAC AGT AGA CAC CTT GGG TAA CAC TGC TAC TCA ATC TGG CCA
T D L S S V E P I V T M S *

### GPR83-EGFP in pTRE3G-BI vector

NheI
GCT AGC ATG GTC CCT CAC CTC TTG CTG CTC TGT CTC CTC CCC TTG GTG CGA GCC ACC GAG CCC CAC GAG GGC CGG GCC GAC GAG CAG AGC GCG GAG GCG GCC CTG
CGA TCG TAC CAG GGA GTG GAG AAC GAC GAG ACA GAG GAG GGG AAC CAC GCT CGG TGG CTC GGG GTG CTC CCG GCC CGG CTG CTC GTC TCG CGC CTC CGC CGG GAC
M V P H L L L L C L L P L V R A T E P H E G R A D E Q S A E A A L

GCC GTG CCC AAT GCC TCG CAC TTC TTC TCT TGG AAC AAC TAC ACC TTC TCC GAC TGG CAG AAC TTT GTG GGC AGG AGG CGC TAC GGC GCT GAG TCC CAG AAC CCC
CGG CAC GGG TTA CGG AGC GTG AAG AAG AGA ACC TTG TTG ATG TGG AAG AGG CTG ACC GTC TTG AAA CAC CCG TCC TCC GCG ATG CCG CGA CTC AGG GTC TTG GGG
A V P N A S H F F S W N N Y T F S D W Q N F V G R R R Y G A E S Q N P

ACG GTG AAA GCC CTG CTC ATT GTG GCT TAC TCC TTC ATC ATT GTC TTC TCA CTC TTT GGC AAC GTC CTG GTC TGT CAT GTC ATC TTC AAG AAC CAG CGA ATG CAC
TGC CAC TTT CGG GAC GAG TAA CAC CGA ATG AGG AAG TAG TAA CAG AAG AGT GAG AAA CCG TTG CAG GAC CAG ACA GTA CAG TAG AAG TTC TTG GTC GCT TAC GTG
T V K A L L I V A Y S F I I V F S L F G N V L V C H V I F K N Q R M H

TCG GCC ACC AGC CTC TTC ATC GTC AAC CTG GCA GTT GCC GAC ATA ATG ATC ACG CTG CTC AAC ACC CCC TTC ACT TTG GTT CGC TTT GTG AAC AGC ACA TGG ATA
AGC CGG TGG TCG GAG AAG TAG CAG TTG GAC CGT CAA CGG CTG TAT TAC TAG TGC GAC GAG TTG TGG GGG AAG TGA AAC CAA GCG AAA CAC TTG TCG TGT ACC TAT
S A T S L F I V N L A V A D I M I T L L N T P F T L V R F V N S T W I

TTT GGG AAG GGC ATG TGC CAT GTC AGC CGC TTT GCC CAG TAC TGC TCA CTG CAC GTC TCA GCA CTG ACA CTG ACA GCC ATT GCG GTG GAT CGC CAC CAG GTC ATC
AAA CCC TTC CCG TAC ACG GTA CAG TCG GCG AAA CGG GTC ATG ACG AGT GAC GTG CAG AGT CGT GAC TGT GAC TGT CGG TAA CGC CAC CTA GCG GTG GTC CAG TAG
F G K G M C H V S R F A Q Y C S L H V S A L T L T A I A V D R H Q V I

ATG CAC CCC TTG AAA CCC CGG ATC TCA ATC ACA AAG GGT GTC ATC TAC ATC GCT GTC ATC TGG ACC ATG GCT ACG TTC TTT TCA CTC CCA CAT GCT ATC TGC CAG
TAC GTG GGG AAC TTT GGG GCC TAG AGT TAG TGT TTC CCA CAG TAG ATG TAG CGA CAG TAG ACC TGG TAC CGA TGC AAG AAA AGT GAG GGT GTA CGA TAG ACG GTC
M H P L K P R I S I T K G V I Y I A V I W T M A T F F S L P H A I C Q

AAA TTA TTT ACC TTC AAA TAC AGT GAG GAC ATT GTG CGC TCC CTC TGC CTG CCA GAC TTC CCT GAG CCA GCT GAC CTC TTC TGG AAG TAC CTG GAC TTG GCC ACC
TTT AAT AAA TGG AAG TTT ATG TCA CTC CTG TAA CAC GCG AGG GAG ACG GAC GGT CTG AAG GGA CTC GGT CGA CTG GAG AAG ACC TTC ATG GAC CTG AAC CGG TGG
K L F T F K Y S E D I V R S L C L P D F P E P A D L F W K Y L D L A T

TTC ATC CTG CTC TAC ATC CTG CCC CTC CTC ATC ATC TCT GTG GCC TAC GCT CGT GTG GCC AAG AAA CTG TGG CTG TGT AAT ATG ATT GGC GAT GTG ACC ACA GAG
AAG TAG GAC GAG ATG TAG GAC GGG GAG GAG TAG TAG AGA CAC CGG ATG CGA GCA CAC CGG TTC TTT GAC ACC GAC ACA TTA TAC TAA CCG CTA CAC TGG TGT CTC
F I L L Y I L P L L I I S V A Y A R V A K K L W L C N M I G D V T T E

CAG TAC TTT GCC CTG CGG CGC AAA AAG AAG AAG ACC ATC AAG ATG TTG ATG CTG GTG GTA GTC CTC TTT GCC CTC TGC TGG TTC CCC CTC AAC TGC TAC GTC CTC
GTC ATG AAA CGG GAC GCC GCG TTT TTC TTC TTC TGG TAG TTC TAC AAC TAC GAC CAC CAT CAG GAG AAA CGG GAG ACG ACC AAG GGG GAG TTG ACG ATG CAG GAG
Q Y F A L R R K K K K T I K M L M L V V V L F A L C W F P L N C Y V L

CTC CTG TCC AGC AAG GTC ATC CGC ACC AAC AAT GCC CTC TAC TTT GCC TTC CAC TGG TTT GCC ATG AGC AGC ACC TGC TAT AAC CCC TTC ATA TAC TGC TGG CTG
GAG GAC AGG TCG TTC CAG TAG GCG TGG TTG TTA CGG GAG ATG AAA CGG AAG GTG ACC AAA CGG TAC TCG TCG TGG ACG ATA TTG GGG AAG TAT ATG ACG ACC GAC
L L S S K V I R T N N A L Y F A F H W F A M S S T C Y N P F I Y C W L

AAC GAG AAC TTC AGG ATT GAG CTA AAG GCA TTA CTG AGC ATG TGT CAA AGA CCT CCC AAG CCT CAG GAG GAC AGG CCA CCC TCC CCA GTT CCT TCC TTC AGG GTG
TTG CTC TTG AAG TCC TAA CTC GAT TTC CGT AAT GAC TCG TAC ACA GTT TCT GGA GGG TTC GGA GTC CTC CTG TCC GGT GGG AGG GGT CAA GGA AGG AAG TCC CAC
N E N F R I E L K A L L S M C Q R P P K P Q E D R P P S P V P S F R V

```
GCC TGG ACA GAG AAG AAT GAT GGC CAG AGG GCC CCC CTT GCC AAT AAC CTC CTG CCC ACC TCC CAA CTC CAG TCT GGG AAG ACA GAC CTG TCA TCT GTG GAA CCC
CGG ACC TGT CTC TTC TTA CTA CCG GTC TCC CGG GGG GAA CGG TTA TTG GAG GAC GGG TGG AGG GTT GAG GTC AGA CCC TTC TGT CTG GAC AGT AGA CAC CTT GGG
 A   W   T   E   K   N   D   G   Q   R   A   P   L   A   N   N   L   L   P   T   S   Q   L   Q   S   G   K   T   D   L   S   S   V   E   P

                         AgeI
ATT GTG ACG ATG AGT CCA CCG GTC GCC ACC ATG GTG AGC AAG GGC GAG GAG CTG TTC ACC GGG GTG GTG CCC ATC CTG GTC GAG CTG GAC GGC GAC GTA AAC GGC
TAA CAC TGC TAC TCA GGT GGC CAG CGG TGG TAC CAC TCG TTC CCG CTC CTC GAC AAG TGG CCC CAC CAC GGG TAG GAC CAG CTC GAC CTG CCG CTG CAT TTG CCG
 I   V   T   M   S   P   P   V   A   T   M   V   S   K   G   E   E   L   F   T   G   V   V   P   I   L   V   E   L   D   G   D   V   N   G

CAC AAG TTC AGC GTG TCC GGC GAG GGC GAG GGC GAT GCC ACC TAC GGC AAG CTG ACC CTG AAG TTC ATC TGC ACC ACC GGC AAG CTG CCC GTG CCC TGG CCC ACC
GTG TTC AAG TCG CAC AGG CCG CTC CCG CTC CCG CTA CGG TGG ATG CCG TTC GAC TGG GAC TTC AAG TAG ACG TGG TGG CCG TTC GAC GGG CAC GGG ACC GGG TGG
 H   K   F   S   V   S   G   E   G   E   G   D   A   T   Y   G   K   L   T   L   K   F   I   C   T   T   G   K   L   P   V   P   W   P   T

CTC GTG ACC ACC CTG ACC TAC GGC GTG CAG TGC TTC AGC CGC TAC CCC GAC CAC ATG AAG CAG CAC GAC TTC TTC AAG TCC GCC ATG CCC GAA GGC TAC GTC CAG
GAG CAC TGG TGG GAC TGG ATG CCG CAC GTC ACG AAG TCG GCG ATG GGG CTG GTG TAC TTC GTC GTG CTG AAG AAG TTC AGG CGG TAC GGG CTT CCG ATG CAG GTC
 L   V   T   T   L   T   Y   G   V   Q   C   F   S   R   Y   P   D   H   M   K   Q   H   D   F   F   K   S   A   M   P   E   G   Y   V   Q

GAG CGC ACC ATC TTC TTC AAG GAC GAC GGC AAC TAC AAG ACC CGC GCC GAG GTG AAG TTC GAG GGC GAC ACC CTG GTG AAC CGC ATC GAG CTG AAG GGC ATC GAC
CTC GCG TGG TAG AAG AAG TTC CTG CTG CCG TTG ATG TTC TGG GCG CGG CTC CAC TTC AAG CTC CCG CTG TGG GAC CAC TTG GCG TAG CTC GAC TTC CCG TAG CTG
 E   R   T   I   F   F   K   D   D   G   N   Y   K   T   R   A   E   V   K   F   E   G   D   T   L   V   N   R   I   E   L   K   G   I   D

TTC AAG GAG GAC GGC AAC ATC CTG GGG CAC AAG CTG GAG TAC AAC TAC AAC AGC CAC AAC GTC TAT ATC ATG GCC GAC AAG CAG AAG AAC GGC ATC AAG GTG AAC
AAG TTC CTC CTG CCG TTG TAG GAC CCC GTG TTC GAC CTC ATG TTG ATG TTG TCG GTG TTG CAG ATA TAG TAC CGG CTG TTC GTC TTC TTG CCG TAG TTC CAC TTG
 F   K   E   D   G   N   I   L   G   H   K   L   E   Y   N   Y   N   S   H   N   V   Y   I   M   A   D   K   Q   K   N   G   I   K   V   N

TTC AAG ATC CGC CAC AAC ATC GAG GAC GGC AGC GTG CAG CTC GCC GAC CAC TAC CAG CAG AAC ACC CCC ATC GGC GAC GGC CCC GTG CTG CTG CCC GAC AAC CAC
AAG TTC TAG GCG GTG TTG TAG CTC CTG CCG TCG CAC GTC GAG CGG CTG GTG ATG GTC GTC TTG TGG GGG TAG CCG CTG CCG GGG CAC GAC GAC GGG CTG TTG GTG
 F   K   I   R   H   N   I   E   D   G   S   V   Q   L   A   D   H   Y   Q   Q   N   T   P   I   G   D   G   P   V   L   L   P   D   N   H

TAC CTG AGC ACC CAG TCC GCC CTG AGC AAA GAC CCC AAC GAG AAG CGC GAT CAC ATG GTC CTG CTG GAG TTC GTG ACC GCC GCC GGG ATC ACT CTC GGC ATG GAC
ATG GAC TCG TGG GTC AGG CGG GAC TCG TTT CTG GGG TTG CTC TTC GCG CTA GTG TAC CAG GAC GAC CTC AAG CAC TGG CGG CGG CCC TAG TGA GAG CCG TAC CTG
 Y   L   S   T   Q   S   A   L   S   K   D   P   N   E   K   R   D   H   M   V   L   L   E   F   V   T   A   A   G   I   T   L   G   M   D

                         NotI
GAG CTG TAC AAG TAA AGC GGC CGC
CTC GAC ATG TTC ATT TCG CCG GCG
 E   L   Y   K   *
```

#### IRES-mediated coexpression of GPR83 and EGFP in pTRE3G-BI vector

```
  NheI
GCT AGC ATG GTC CCT CAC CTC TTG CTG CTC TGT CTC CTC CCC TTG GTG CGA GCC ACC GAG CCC CAC GAG GGC CGG GCC GAC GAG CAG AGC GCG GAG GCG GCC CTG
CGA TCG TAC CAG GGA GTG GAG AAC GAC GAG ACA GAG GAG GGG AAC CAC GCT CGG TGG CTC GGG GTG CTC CCG GCC CGG CTG CTC GTC TCG CGC CTC CGC CGG GAC
         M   V   P   H   L   L   L   L   C   L   L   P   L   V   R   A   T   E   P   H   E   G   R   A   D   E   Q   S   A   E   A   A   L

GCC GTG CCC AAT GCC TCG CAC TTC TTC TCT TGG AAC AAC TAC ACC TTC TCC GAC TGG CAG AAC TTT GTG GGC AGG AGG CGC TAC GGC GCT GAG TCC CAG AAC CCC
CGG CAC GGG TTA CGG AGC GTG AAG AAG AGA ACC TTG TTG ATG TGG AAG AGG CTG ACC GTC TTG AAA CAC CCG TCC TCC GCG ATG CCG CGA CTC AGG GTC TTG GGG
 A   V   P   N   A   S   H   F   F   S   W   N   N   Y   T   F   S   D   W   Q   N   F   V   G   R   R   R   Y   G   A   E   S   Q   N   P

ACG GTG AAA GCC CTG CTC ATT GTG GCT TAC TCC TTC ATC ATT GTC TTC TCA CTC TTT GGC AAC GTC CTG GTC TGT CAT GTC ATC TTC AAG AAC CAG CGA ATG CAC
TGC CAC TTT CGG GAC GAG TAA CAC CGA ATG AGG AAG TAG TAA CAG AAG AGT GAG AAA CCG TTG CAG GAC CAG ACA GTA CAG TAG AAG TTC TTG GTC GCT TAC GTG
 T   V   K   A   L   L   I   V   A   Y   S   F   I   I   V   F   S   L   F   G   N   V   L   V   C   H   V   I   F   K   N   Q   R   M   H

TCG GCC ACC AGC CTC TTC ATC GTC AAC CTG GCA GTT GCC GAC ATA ATG ATC ACG CTG CTC AAC ACC CCC TTC ACT TTG GTT CGC TTT GTG AAC AGC ACA TGG ATA
AGC CGG TGG TCG GAG AAG TAG CAG TTG GAC CGT CAA CGG CTG TAT TAC TAG TGC GAC GAG TTG TGG GGG AAG TGA AAC CAA GCG AAA CAC TTG TCG TGT ACC TAT
 S   A   T   S   L   F   I   V   N   L   A   V   A   D   I   M   I   T   L   L   N   T   P   F   T   L   V   R   F   V   N   S   T   W   I

TTT GGG AAG GGC ATG TGC CAT GTC AGC CGC TTT GCC CAG TAC TGC TCA CTG CAC GTC TCA GCA CTG ACA CTG ACA GCC ATT GCG GTG GAT CGC CAC CAG GTC ATC
AAA CCC TTC CCG TAC ACG GTA CAG TCG GCG AAA CGG GTC ATG ACG AGT GAC GTG CAG AGT CGT GAC TGT GAC TGT CGG TAA CGC CAC CTA GCG GTG GTC CAG TAG
 F   G   K   G   M   C   H   V   S   R   F   A   Q   Y   C   S   L   H   V   S   A   L   T   L   T   A   I   A   V   D   R   H   Q   V   I

ATG CAC CCC TTG AAA CCC CGG ATC TCA ATC ACA AAG GGT GTC ATC TAC ATC GCT GTC ATC TGG ACC ATG GCT ACG TTC TTT TCA CTC CCA CAT GCT ATC TGC CAG
TAC GTG GGG AAC TTT GGG GCC TAG AGT TAG TGT TTC CCA CAG TAG ATG TAG CGA CAG TAG ACC TGG TAC CGA TGC AAG AAA AGT GAG GGT GTA CGA TAG ACG GTC
 M   H   P   L   K   P   R   I   S   I   T   K   G   V   I   Y   I   A   V   I   W   T   M   A   T   F   F   S   L   P   H   A   I   C   Q

AAA TTA TTT ACC TTC AAA TAC AGT GAG GAC ATT GTG CGC TCC CTC TGC CTG CCA GAC TTC CCT GAG CCA GCT GAC CTC TTC TGG AAG TAC CTG GAC TTG GCC ACC
TTT AAT AAA TGG AAG TTT ATG TCA CTC CTG TAA CAC GCG AGG GAG ACG GAC GGT CTG AAG GGA CTC GGT CGA CTG GAG AAG ACC TTC ATG GAC CTG AAC CGG TGG
 K   L   F   T   F   K   Y   S   E   D   I   V   R   S   L   C   L   P   D   F   P   E   P   A   D   L   F   W   K   Y   L   D   L   A   T

TTC ATC CTG CTC TAC ATC CTG CCC CTC CTC ATC ATC TCT GTG GCC TAC GCT CGT GTG GCC AAG AAA CTG TGG CTG TGT AAT ATG ATT GGC GAT GTG ACC ACA GAG
AAG TAG GAC GAG ATG TAG GAC GGG GAG GAG TAG TAG AGA CAC CGG ATG CGA GCA CAC CGG TTC TTT GAC ACC GAC ACA TTA TAC TAA CCG CTA CAC TGG TGT CTC
 F   I   L   L   Y   I   L   P   L   L   I   I   S   V   A   Y   A   R   V   A   K   K   L   W   L   C   N   M   I   G   D   V   T   T   E

CAG TAC TTT GCC CTG CGG CGC AAA AAG AAG AAG ACC ATC AAG ATG TTG ATG CTG GTG GTA GTC CTC TTT GCC CTC TGC TGG TTC CCC CTC AAC TGC TAC GTC CTC
GTC ATG AAA CGG GAC GCC GCG TTT TTC TTC TTC TGG TAG TTC TAC AAC TAC GAC CAC CAT CAG GAG AAA CGG GAG ACG ACC AAG GGG GAG TTG ACG ATG CAG GAG
 Q   Y   F   A   L   R   R   K   K   K   K   T   I   K   M   L   M   L   V   V   V   L   F   A   L   C   W   F   P   L   N   C   Y   V   L

CTC CTG TCC AGC AAG GTC ATC CGC ACC AAC AAT GCC CTC TAC TTT GCC TTC CAC TGG TTT GCC ATG AGC AGC ACC TGC TAT AAC CCC TTC ATA TAC TGC TGG CTG
GAG GAC AGG TCG TTC CAG TAG GCG TGG TTG TTA CGG GAG ATG AAA CGG AAG GTG ACC AAA CGG TAC TCG TCG TGG ACG ATA TTG GGG AAG TAT ATG ACG ACC GAC
 L   L   S   S   K   V   I   R   T   N   N   A   L   Y   F   A   F   H   W   F   A   M   S   S   T   C   Y   N   P   F   I   Y   C   W   L

AAC GAG AAC TTC AGG ATT GAG CTA AAG GCA TTA CTG AGC ATG TGT CAA AGA CCT CCC AAG CCT CAG GAG GAC AGG CCA CCC TCC CCA GTT CCT TCC TTC AGG GTG
TTG CTC TTG AAG TCC TAA CTC GAT TTC CGT AAT GAC TCG TAC ACA GTT TCT GGA GGG TTC GGA GTC CTC CTG TCC GGT GGG AGG GGT CAA GGA AGG AAG TCC CAC
 N   E   N   F   R   I   E   L   K   A   L   L   S   M   C   Q   R   P   P   K   P   Q   E   D   R   P   P   S   P   V   P   S   F   R   V
```

```
GCC TGG ACA GAG AAG AAT GAT GGC CAG AGG GCC CCC CTT GCC AAT AAC CTC CTG CCC ACC TCC CAA CTC CAG TCT GGG AAG ACA GAC CTG TCA TCT GTG GAA CCC
CGG ACC TGT CTC TTC TTA CTA CCG GTC TCC CGG GGG GAA CGG TTA TTG GAG GAC GGG TGG AGG GTT GAG GTC AGA CCC TTC TGT CTG GAC AGT AGA CAC CTT GGG
 A   W   T   E   K   N   D   G   Q   R   A   P   L   A   N   N   L   L   P   T   S   Q   L   Q   S   G   K   T   D   L   S   S   V   E   P

                        AgeI
ATT GTG ACG ATG AGT CCA CCG GTC TGA TAA GCG GCC GCG CCC CTC TCC CTC CCC CCC CCC TAA CGT TAC TGG CCG AAG CCG CTT GGA ATA AGG CCG GTG TGC GTT
TAA CAC TGC TAC TCA GGT GGC CAG ACT ATT CGC CGG CGC GGG GAG AGG GAG GGG GGG GGG ATT GCA ATG ACC GGC TTC GGC GAA CCT TAT TCC GGC CAC ACG CAA
 I   V   T   M   S   P   P   V   *

TGT CTA TAT GTT ATT TTC CAC CAT ATT GCC GTC TTT TGG CAA TGT GAG GGC CCG GAA ACC TGG CCC TGT CTT CTT GAC GAG CAT TCC TAG GGG TCT TTC CCC TCT
ACA GAT ATA CAA TAA AAG GTG GTA TAA CGG CAG AAA ACC GTT ACA CTC CCG GGC CTT TGG ACC GGG ACA GAA GAA CTG CTC GTA AGG ATC CCC AGA AAG GGG AGA

CGC CAA AGG AAT GCA AGG TCT GTT GAA TGT CGT GAA GGA AGC AGT TCC TCT GGA AGC TTC TTG AAG ACA AAC AAC GTC TGT AGC GAC CCT TTG CAG GCA GCG GAA
GCG GTT TCC TTA CGT TCC AGA CAA CTT ACA GCA CTT CCT TCG TCA AGG AGA CCT TCG AAG AAC TTC TGT TTG TTG CAG ACA TCG CTG GGA AAC GTC CGT CGC CTT

CCC CCC ACC TGG CGA CAG GTG CCT CTG CGG CCA AAA GCC ACG TGT ATA AGA TAC ACC TGC AAA GGC GGC ACA ACC CCA GTG CCA CGT TGT GAG TTG GAT AGT TGT
GGG GGG TGG ACC GCT GTC CAC GGA GAC GCC GGT TTT CGG TGC ACA TAT TCT ATG TGG ACG TTT CCG CCG TGT TGG GGT CAC GGT GCA ACA CTC AAC CTA TCA ACA

GGA AAG AGT CAA ATG GCT CTC CTC AAG CGT ATT CAA CAA GGG GCT GAA GGA TGC CCA GAA GGT ACC CCA TTG TAT GGG ATC TGA TCT GGG GCC TCG GTG CAC ATG
CCT TTC TCA GTT TAC CGA GAG GAG TTC GCA TAA GTT GTT CCC CGA CTT CCT ACG GGT CTT CCA TGG GGT AAC ATA CCC TAG ACT AGA CCC CGG AGC CAC GTG TAC

CTT TAC ATG TGT TTA GTC GAG GTT AAA AAA ACG TCT AGG CCC CCC GAA CCA CGG GGA CGT GGT TTT CCT TTG AAA AAC ACG ATG ATA AT
GAA ATG TAC ACA AAT CAG CTC CAA TTT TTT TGC AGA TCC GGG GGG CTT GGT GCC CCT GCA CCA AAA GGA AAC TTT TTG TGC TAC TAT TA

ATG GCC ACA ACC ATG GTG AGC AAG GGC GAG GAG CTG TTC ACC GGG GTG GTG CCC ATC CTG GTC GAG CTG GAC GGC GAC GTA AAC GGC CAC AAG TTC AGC GTG TCC
TAC CGG TGT TGG TAC CAC TCG TTC CCG CTC CTC GAC AAG TGG CCC CAC CAC GGG TAG GAC CAG CTC GAC CTG CCG CTG CAT TTG CCG GTG TTC AAG TCG CAC AGG
 M   A   T   T   M   V   S   K   G   E   E   L   F   T   G   V   V   P   I   L   V   E   L   D   G   D   V   N   G   H   K   F   S   V   S

GGC GAG GGC GAG GGC GAT GCC ACC TAC GGC AAG CTG ACC CTG AAG TTC ATC TGC ACC ACC GGC AAG CTG CCC GTG CCC TGG CCC ACC CTC GTG ACC ACC CTG ACC
CCG CTC CCG CTC CCG CTA CGG TGG ATG CCG TTC GAC TGG GAC TTC AAG TAG ACG TGG TGG CCG TTC GAC GGG CAC GGG ACC GGG TGG GAG CAC TGG TGG GAC TGG
 G   E   G   E   G   D   A   T   Y   G   K   L   T   L   K   F   I   C   T   T   G   K   L   P   V   P   W   P   T   L   V   T   T   L   T

TAC GGC GTG CAG TGC TTC AGC CGC TAC CCC GAC CAC ATG AAG CAG CAC GAC TTC TTC AAG TCC GCC ATG CCC GAA GGC TAC GTC CAG GAG CGC ACC ATC TTC TTC
ATG CCG CAC GTC ACG AAG TCG GCG ATG GGG CTG GTG TAC TTC GTC GTG CTG AAG AAG TTC AGG CGG TAC GGG CTT CCG ATG CAG GTC CTC GCG TGG TAG AAG AAG
 Y   G   V   Q   C   F   S   R   Y   P   D   H   M   K   Q   H   D   F   F   K   S   A   M   P   E   G   Y   V   Q   E   R   T   I   F   F

AAG GAC GAC GGC AAC TAC AAG ACC CGC GCC GAG GTG AAG TTC GAG GGC GAC ACC CTG GTG AAC CGC ATC GAG CTG AAG GGC ATC GAC TTC AAG GAG GAC GGC AAC
TTC CTG CTG CCG TTG ATG TTC TGG GCG CGG CTC CAC TTC AAG CTC CCG CTG TGG GAC CAC TTG GCG TAG CTC GAC TTC CCG TAG CTG AAG TTC CTC CTG CCG TTG
 K   D   D   G   N   Y   K   T   R   A   E   V   K   F   E   G   D   T   L   V   N   R   I   E   L   K   G   I   D   F   K   E   D   G   N

ATC CTG GGG CAC AAG CTG GAG TAC AAC TAC AAC AGC CAC AAC GTC TAT ATC ATG GCC GAC AAG CAG AAG AAC GGC ATC AAG GTG AAC TTC AAG ATC CGC CAC AAC
TAG GAC CCC GTG TTC GAC CTC ATG TTG ATG TTG TCG GTG TTG CAG ATA TAG TAC CGG CTG TTC GTC TTC TTG CCG TAG TTC CAC TTG AAG TTC TAG GCG GTG TTG
 I   L   G   H   K   L   E   Y   N   Y   N   S   H   N   V   Y   I   M   A   D   K   Q   K   N   G   I   K   V   N   F   K   I   R   H   N

ATC GAG GAC GGC AGC GTG CAG CTC GCC GAC CAC TAC CAG CAG AAC ACC CCC ATC GGC GAC GGC CCC GTG CTG CTG CCC GAC AAC CAC TAC CTG AGC ACC CAG TCC
TAG CTC CTG CCG TCG CAC GTC GAG CGG CTG GTG ATG GTC GTC TTG TGG GGG TAG CCG CTG CCG GGG CAC GAC GAC GGG CTG TTG GTG ATG GAC TCG TGG GTC AGG
 I   E   D   G   S   V   Q   L   A   D   H   Y   Q   Q   N   T   P   I   G   D   G   P   V   L   L   P   D   N   H   Y   L   S   T   Q   S
```

```
GCC CTG AGC AAA GAC CCC AAC GAG AAG CGC GAT CAC ATG GTC CTG CTG GAG TTC GTG ACC GCC GCC GGG ATC ACT CTC GGC ATG GAC GAG CTG TAC AAG TAA GATATC
CGG GAC TCG TTT CTG GGG TTG CTC TTC GCG CTA GTG TAC CAG GAC GAC CTC AAG CAC TGG CGG CGG CCC TAG TGA GAG CCG TAC CTG CTC GAC ATG TTC ATT CTATAG
A   L   S   K   D   P   N   E   K   R   D   H   M   V   L   L   E   F   V   T   A   A   G   I   T   L   G   M   D   E   L   Y   K   *
```

**Fig. S4.** The nucleotide and amino acid sequences of human GPR83 constructs expressed in mammalian cells. The amino acid sequence of the full-length human GPR83 (1-423) or mature human GPR83 (17-423) is shown in red. The N-terminal signal peptide of full-length GPR83 and sLgBiT is shaded. The amino acid sequence of sLgBiT is shown in blue, and that of EGFP in green. The restriction enzyme cleavage sites for cloning are shaded. The nucleotide sequence of IRES is shown in pink.

#### GPR83-LgBiT in MCS2 of pTRE3G-BI

```
     EcoRI
1    GAA TTC ATG GTC CCT CAC CTC TTG CTG CTC TGT CTC CTC CCC TTG GTG CGA GCC ACC GAG CCC CAC GAG GGC CGG GCC GAC GAG CAG AGC GCG GAG GCG GCC CTG
     CTT AAG TAC CAG GGA GTG GAG AAC GAC GAG ACA GAG GAG GGG AAC CAC GCT CGG TGG CTC GGG GTG CTC CCG GCC CGG CTG CTC GTC TCG CGC CTC CGC CGG GAC
              M   V   P   H   L   L   L   L   C   L   L   P   L   V   R   A   T   E   P   H   E   G   R   A   D   E   Q   S   A   E   A   A   L

106  GCC GTG CCC AAT GCC TCG CAC TTC TTC TCT TGG AAC AAC TAC ACC TTC TCC GAC TGG CAG AAC TTT GTG GGC AGG AGG CGC TAC GGC GCT GAG TCC CAG AAC CCC
     CGG CAC GGG TTA CGG AGC GTG AAG AAG AGA ACC TTG TTG ATG TGG AAG AGG CTG ACC GTC TTG AAA CAC CCG TCC TCC GCG ATG CCG CGA CTC AGG GTC TTG GGG
      A   V   P   N   A   S   H   F   F   S   W   N   N   Y   T   F   S   D   W   Q   N   F   V   G   R   R   R   Y   G   A   E   S   Q   N   P

211  ACG GTG AAA GCC CTG CTC ATT GTG GCT TAC TCC TTC ATC ATT GTC TTC TCA CTC TTT GGC AAC GTC CTG GTC TGT CAT GTC ATC TTC AAG AAC CAG CGA ATG CAC
     TGC CAC TTT CGG GAC GAG TAA CAC CGA ATG AGG AAG TAG TAA CAG AAG AGT GAG AAA CCG TTG CAG GAC CAG ACA GTA CAG TAG AAG TTC TTG GTC GCT TAC GTG
      T   V   K   A   L   L   I   V   A   Y   S   F   I   I   V   F   S   L   F   G   N   V   L   V   C   H   V   I   F   K   N   Q   R   M   H

316  TCG GCC ACC AGC CTC TTC ATC GTC AAC CTG GCA GTT GCC GAC ATA ATG ATC ACG CTG CTC AAC ACC CCC TTC ACT TTG GTT CGC TTT GTG AAC AGC ACA TGG ATA
     AGC CGG TGG TCG GAG AAG TAG CAG TTG GAC CGT CAA CGG CTG TAT TAC TAG TGC GAC GAG TTG TGG GGG AAG TGA AAC CAA GCG AAA CAC TTG TCG TGT ACC TAT
      S   A   T   S   L   F   I   V   N   L   A   V   A   D   I   M   I   T   L   L   N   T   P   F   T   L   V   R   F   V   N   S   T   W   I

421  TTT GGG AAG GGC ATG TGC CAT GTC AGC CGC TTT GCC CAG TAC TGC TCA CTG CAC GTC TCA GCA CTG ACA CTG ACA GCC ATT GCG GTG GAT CGC CAC CAG GTC ATC
     AAA CCC TTC CCG TAC ACG GTA CAG TCG GCG AAA CGG GTC ATG ACG AGT GAC GTG CAG AGT CGT GAC TGT GAC TGT CGG TAA CGC CAC CTA GCG GTG GTC CAG TAG
      F   G   K   G   M   C   H   V   S   R   F   A   Q   Y   C   S   L   H   V   S   A   L   T   L   T   A   I   A   V   D   R   H   Q   V   I

526  ATG CAC CCC TTG AAA CCC CGG ATC TCA ATC ACA AAG GGT GTC ATC TAC ATC GCT GTC ATC TGG ACC ATG GCT ACG TTC TTT TCA CTC CCA CAT GCT ATC TGC CAG
     TAC GTG GGG AAC TTT GGG GCC TAG AGT TAG TGT TTC CCA CAG TAG ATG TAG CGA CAG TAG ACC TGG TAC CGA TGC AAG AAA AGT GAG GGT GTA CGA TAG ACG GTC
      M   H   P   L   K   P   R   I   S   I   T   K   G   V   I   Y   I   A   V   I   W   T   M   A   T   F   F   S   L   P   H   A   I   C   Q

631  AAA TTA TTT ACC TTC AAA TAC AGT GAG GAC ATT GTG CGC TCC CTC TGC CTG CCA GAC TTC CCT GAG CCA GCT GAC CTC TTC TGG AAG TAC CTG GAC TTG GCC ACC
     TTT AAT AAA TGG AAG TTT ATG TCA CTC CTG TAA CAC GCG AGG GAG ACG GAC GGT CTG AAG GGA CTC GGT CGA CTG GAG AAG ACC TTC ATG GAC CTG AAC CGG TGG
      K   L   F   T   F   K   Y   S   E   D   I   V   R   S   L   C   L   P   D   F   P   E   P   A   D   L   F   W   K   Y   L   D   L   A   T

736  TTC ATC CTG CTC TAC ATC CTG CCC CTC CTC ATC ATC TCT GTG GCC TAC GCT CGT GTG GCC AAG AAA CTG TGG CTG TGT AAT ATG ATT GGC GAT GTG ACC ACA GAG
     AAG TAG GAC GAG ATG TAG GAC GGG GAG GAG TAG TAG AGA CAC CGG ATG CGA GCA CAC CGG TTC TTT GAC ACC GAC ACA TTA TAC TAA CCG CTA CAC TGG TGT CTC
      F   I   L   L   Y   I   L   P   L   L   I   I   S   V   A   Y   A   R   V   A   K   K   L   W   L   C   N   M   I   G   D   V   T   T   E

841  CAG TAC TTT GCC CTG CGG CGC AAA AAG AAG AAG ACC ATC AAG ATG TTG ATG CTG GTG GTA GTC CTC TTT GCC CTC TGC TGG TTC CCC CTC AAC TGC TAC GTC CTC
     GTC ATG AAA CGG GAC GCC GCG TTT TTC TTC TTC TGG TAG TTC TAC AAC TAC GAC CAC CAT CAG GAG AAA CGG GAG ACG ACC AAG GGG GAG TTG ACG ATG CAG GAG
      Q   Y   F   A   L   R   R   K   K   K   K   T   I   K   M   L   M   L   V   V   V   L   F   A   L   C   W   F   P   L   N   C   Y   V   L

946  CTC CTG TCC AGC AAG GTC ATC CGC ACC AAC AAT GCC CTC TAC TTT GCC TTC CAC TGG TTT GCC ATG AGC AGC ACC TGC TAT AAC CCC TTC ATA TAC TGC TGG CTG
     GAG GAC AGG TCG TTC CAG TAG GCG TGG TTG TTA CGG GAG ATG AAA CGG AAG GTG ACC AAA CGG TAC TCG TCG TGG ACG ATA TTG GGG AAG TAT ATG ACG ACC GAC
      L   L   S   S   K   V   I   R   T   N   N   A   L   Y   F   A   F   H   W   F   A   M   S   S   T   C   Y   N   P   F   I   Y   C   W   L

1051 AAC GAG AAC TTC AGG ATT GAG CTA AAG GCA TTA CTG AGC ATG TGT CAA AGA CCT CCC AAG CCT CAG GAG GAC AGG CCA CCC TCC CCA GTT CCT TCC TTC AGG GTG
     TTG CTC TTG AAG TCC TAA CTC GAT TTC CGT AAT GAC TCG TAC ACA GTT TCT GGA GGG TTC GGA GTC CTC CTG TCC GGT GGG AGG GGT CAA GGA AGG AAG TCC CAC
      N   E   N   F   R   I   E   L   K   A   L   L   S   M   C   Q   R   P   P   K   P   Q   E   D   R   P   P   S   P   V   P   S   F   R   V
```

```
1156 GCC TGG ACA GAG AAG AAT GAT GGC CAG AGG GCT CCC CTT GCC AAT AAC CTC CTG CCC ACC TCC CAA CTC CAG TCT GGG AAG ACA GAC CTG TCA TCT GTG GAA CCC
     CGG ACC TGT CTC TTC TTA CTA CCG GTC TCC CGA GGG GAA CGG TTA TTG GAG GAC GGG TGG AGG GTT GAG GTC AGA CCC TTC TGT CTG GAC AGT AGA CAC CTT GGG
      A   W   T   E   K   N   D   G   Q   R   A   P   L   A   N   N   L   L   P   T   S   Q   L   Q   S   G   K   T   D   L   S   S   V   E   P

                         AgeI
1261 ATT GTG ACG ATG AGT ACC GGT ACC GGC GGA GGG TCT AGC AGT GGC GGT GGG ATG GTC TTC ACA CTC GAA GAT TTC GTT GGG GAC TGG GAA CAG ACA GCC GCC TAC
     TAA CAC TGC TAC TCA TGG CCA TGG CCG CCT CCC AGA TCG TCA CCG CCA CCC TAC CAG AAG TGT GAG CTT CTA AAG CAA CCC CTG ACC CTT GTC TGT CGG CGG ATG
      I   V   T   M   S   T   G   T   G   G   G   S   S   S   G   G   G   M   V   F   T   L   E   D   F   V   G   D   W   E   Q   T   A   A   Y

1366 AAC CTG GAC CAA GTC CTT GAA CAG GGA GGT GTG TCC AGT TTG CTG CAG AAT CTC GCC GTG TCC GTA ACT CCG ATC CAA AGG ATT GTC CGG AGC GGT GAA AAT GCC
     TTG GAC CTG GTT CAG GAA CTT GTC CCT CCA CAC AGG TCA AAC GAC GTC TTA GAG CGG CAC AGG CAT TGA GGC TAG GTT TCC TAA CAG GCC TCG CCA CTT TTA CGG
      N   L   D   Q   V   L   E   Q   G   G   V   S   S   L   L   Q   N   L   A   V   S   V   T   P   I   Q   R   I   V   R   S   G   E   N   A

1471 CTG AAG ATC GAC ATC CAT GTC ATC ATC CCG TAT GAA GGT CTG AGC GCC GAC CAA ATG GCC CAG ATC GAA GAG GTG TTT AAG GTG GTG TAC CCT GTG GAT GAT CAT
     GAC TTC TAG CTG TAG GTA CAG TAG TAG GGC ATA CTT CCA GAC TCG CGG CTG GTT TAC CGG GTC TAG CTT CTC CAC AAA TTC CAC CAC ATG GGA CAC CTA CTA GTA
      L   K   I   D   I   H   V   I   I   P   Y   E   G   L   S   A   D   Q   M   A   Q   I   E   E   V   F   K   V   V   Y   P   V   D   D   H

1576 CAC TTT AAG GTG ATC CTG CCC TAT GGC ACA CTG GTA ATC GAC GGG GTT ACG CCG AAC ATG CTG AAC TAT TTC GGA CGG CCG TAT GAA GGC ATC GCC GTG TTC GAC
     GTG AAA TTC CAC TAG GAC GGG ATA CCG TGT GAC CAT TAG CTG CCC CAA TGC GGC TTG TAC GAC TTG ATA AAG CCT GCC GGC ATA CTT CCG TAG CGG CAC AAG CTG
      H   F   K   V   I   L   P   Y   G   T   L   V   I   D   G   V   T   P   N   M   L   N   Y   F   G   R   P   Y   E   G   I   A   V   F   D

1681 GGC AAA AAG ATC ACT GTA ACA GGG ACC CTG TGG AAC GGC AAC AAA ATT ATC GAC GAG CGC CTG ATC ACC CCC GAC GGC TCC ATG CTG TTC CGA GTA ACC ATC AAC
     CCG TTT TTC TAG TGA CAT TGT CCC TGG GAC ACC TTG CCG TTG TTT TAA TAG CTG CTC GCG GAC TAG TGG GGG CTG CCG AGG TAC GAC AAG GCT CAT TGG TAG TTG
      G   K   K   I   T   V   T   G   T   L   W   N   G   N   K   I   I   D   E   R   L   I   T   P   D   G   S   M   L   F   R   V   T   I   N

             NdeI
1786 AGT TAA CAT ATG
     TCA ATT GTA TAC
      S   *
```

#### SmBiT-ARRB2 in MCS-1 of pTRE3G-BI

```
      BamHI
1    GGA TCC ATG GGA GTG ACC GGC TAC CGG CTG TTC GAG GAG ATT CTG GGC GGC TCT AGT GGT GGA GGG GGC AGC GGC GGA GGT GGG GAG AAA CCC GGG ACC AGG GTC
     CCT AGG TAC CCT CAC TGG CCG ATG GCC GAC AAG CTC CTC TAA GAC CCG CCG AGA TCA CCA CCT CCC CCG TCG CCG CCT CCA CCC CTC TTT GGG CCC TGG TCC CAG
              M   G   V   T   G   Y   R   L   F   E   E   I   L   G   G   S   S   G   G   G   G   S   G   G   G   G   E   K   P   G   T   R   V

106  TTC AAG AAG TCG AGC CCT AAC TGC AAG CTC ACC GTG TAC TTG GGC AAG CGG GAC TTC GTA GAT CAC CTG GAC AAA GTG GAC CCT GTA GAT GGC GTG GTG CTT GTG
     AAG TTC TTC AGC TCG GGA TTG ACG TTC GAG TGG CAC ATG AAC CCG TTC GCC CTG AAG CAT CTA GTG GAC CTG TTT CAC CTG GGA CAT CTA CCG CAC CAC GAA CAC
      F   K   K   S   S   P   N   C   K   L   T   V   Y   L   G   K   R   D   F   V   D   H   L   D   K   V   D   P   V   D   G   V   V   L   V

211  GAC CCT GAC TAC CTG AAG GAC CGC AAA GTG TTT GTG ACC CTC ACC TGC GCC TTC CGC TAT GGC CGT GAA GAC CTG GAT GTG CTG GGC TTG TCC TTC CGC AAA GAC
     CTG GGA CTG ATG GAC TTC CTG GCG TTT CAC AAA CAC TGG GAG TGG ACG CGG AAG GCG ATA CCG GCA CTT CTG GAC CTA CAC GAC CCG AAC AGG AAG GCG TTT CTG
      D   P   D   Y   L   K   D   R   K   V   F   V   T   L   T   C   A   F   R   Y   G   R   E   D   L   D   V   L   G   L   S   F   R   K   D
```

316 CTG TTC ATC GCC ACC TAC CAG GCC TTC CCC CCG GTG CCC AAC CCA CCC CGG CCC CCC ACC CGC CTG CAG GAC CGG CTG CTG AGG AAG CTG GGC CAG CAT GCC CAC
GAC AAG TAG CGG TGG ATG GTC CGG AAG GGG GGC CAC GGG TTG GGT GGG GCC GGG GGG TGG GCG GAC GTC CTG GCC GAC GAC TCC TTC GAC CCG GTC GTA CGG GTG
L F I A T Y Q A F P P V P N P P R P P T R L Q D R L L R K L G Q H A H

421 CCC TTC TTC TTC ACC ATA CCC CAG AAT CTT CCA TGC TCC GTC ACA CTG CAG CCA GGC CCA GAG GAT ACA GGA AAG GCC TGC GGC GTA GAC TTT GAG ATT CGA GCC
GGG AAG AAG AAG TGG TAT GGG GTC TTA GAA GGT ACG AGG CAG TGT GAC GTC GGT CCG GGT CTC CTA TGT CCT TTC CGG ACG CCG CAT CTG AAA CTC TAA GCT CGG
P F F F T I P Q N L P C S V T L Q P G P E D T G K A C G V D F E I R A

526 TTC TGT GCT AAA TCA CTA GAA GAG AAA AGC CAC AAA AGG AAC TCT GTG CGG CTG GTG ATC CGA AAG GTG CAG TTC GCC CCG GAG AAA CCC GGC CCC CAG CCT TCA
AAG ACA CGA TTT AGT GAT CTT CTC TTT TCG GTG TTT TCC TTG AGA CAC GCC GAC CAC TAG GCT TTC CAC GTC AAG CGG GGC CTC TTT GGG CCG GGG GTC GGA AGT
F C A K S L E E K S H K R N S V R L V I R K V Q F A P E K P G P Q P S

631 GCC GAA ACC ACA CGC CAC TTC CTC ATG TCT GAC CGG TCC CTG CAC CTC GAG GCT TCC CTG GAC AAG GAG CTG TAC TAC CAT GGG GAG CCC CTC AAT GTA AAT GTC
CGG CTT TGG TGT GCG GTG AAG GAG TAC AGA CTG GCC AGG GAC GTG GAG CTC CGA AGG GAC CTG TTC CTC GAC ATG ATG GTA CCC CTC GGG GAG TTA CAT TTA CAG
A E T T R H F L M S D R S L H L E A S L D K E L Y Y H G E P L N V N V

736 CAC GTC ACC AAC AAC TCC ACC AAG ACC GTC AAG AAG ATC AAA GTC TCT GTG AGA CAG TAC GCC GAC ATC TGC CTC TTC AGC ACC GCC CAG TAC AAG TGT CCT GTG
GTG CAG TGG TTG TTG AGG TGG TTC TGG CAG TTC TTC TAG TTT CAG AGA CAC TCT GTC ATG CGG CTG TAG ACG GAG AAG TCG TGG CGG GTC ATG TTC ACA GGA CAC
H V T N N S T K T V K K I K V S V R Q Y A D I C L F S T A Q Y K C P V

841 GCT CAA CTC GAA CAA GAT GAC CAG GTA TCT CCC AGC TCC ACA TTC TGT AAG GTG TAC ACC ATA ACC CCA CTG CTC AGC GAC AAC CGG GAG AAG CGG GGT CTC GCC
CGA GTT GAG CTT GTT CTA CTG GTC CAT AGA GGG TCG AGG TGT AAG ACA TTC CAC ATG TGG TAT TGG GGT GAC GAG TCG CTG TTG GCC CTC TTC GCC CCA GAG CGG
A Q L E Q D D Q V S P S S T F C K V Y T I T P L L S D N R E K R G L A

946 CTG GAT GGG AAA CTC AAG CAC GAG GAC ACC AAC CTG GCT TCC AGC ACC ATC GTG AAG GAG GGT GCC AAC AAG GAG GTG CTG GGA ATC CTG GTG TCC TAC AGG GTC
GAC CTA CCC TTT GAG TTC GTG CTC CTG TGG TTG GAC CGA AGG TCG TGG TAG CAC TTC CTC CCA CGG TTG TTC CTC CAC GAC CCT TAG GAC CAC AGG ATG TCC CAG
L D G K L K H E D T N L A S S T I V K E G A N K E V L G I L V S Y R V

1051 AAG GTG AAG CTG GTG GTG TCT CGA GGC GGG GAT GTC TCT GTG GAG CTG CCT TTT GTT CTT ATG CAC CCC AAG CCC CAC GAC CAC ATC CCC CTC CCC AGA CCC CAG
TTC CAC TTC GAC CAC CAC AGA GCT CCG CCC CTA CAG AGA CAC CTC GAC GGA AAA CAA GAA TAC GTG GGG TTC GGG GTG CTG GTG TAG GGG GAG GGG TCT GGG GTC
K V K L V V S R G G D V S V E L P F V L M H P K P H D H I P L P R P Q

1156 TCA GCC GCT CCG GAG ACA GAT GTC CCT GTG GAC ACC AAC CTC ATT GAA TTT GAT ACC AAC TAT GCC ACA GAT GAT GAC ATT GTG TTT GAG GAC TTT GCC CGG CTT
AGT CGG CGA GGC CTC TGT CTA CAG GGA CAC CTG TGG TTG GAG TAA CTT AAA CTA TGG TTG ATA CGG TGT CTA CTA CTG TAA CAC AAA CTC CTG AAA CGG GCC GAA
S A A P E T D V P V D T N L I E F D T N Y A T D D D I V F E D F A R L

Not I
1261 CGG CTG AAG GGG ATG AAG GAT GAC GAC TAT GAT GAT CAA CTC TGC TAG GCG GCC GC
GCC GAC TTC CCC TAC TTC CTA CTG CTG ATA CTA CTA GTT GAG ACG ATC CGC CGG CG
R L K G M K D D D Y D D Q L C *

**Fig. S5.** The nucleotide and amino acid sequences of GPR83-LgBiT and SmBiT-ARRB2 in pTRE3G-BI vector for coexpression in mammalian cells under a controllable manner. The amino acid sequence of the full-length human GPR83 (1-423) is shown in red, and that of ARRB2 in green. The amino acid sequence of LgBiT and SmBiT is shown in blue. The restriction enzyme cleavage sites for cloning are shaded.

1 130

#### Alignment of PCSK1N/proSAAS orthologs

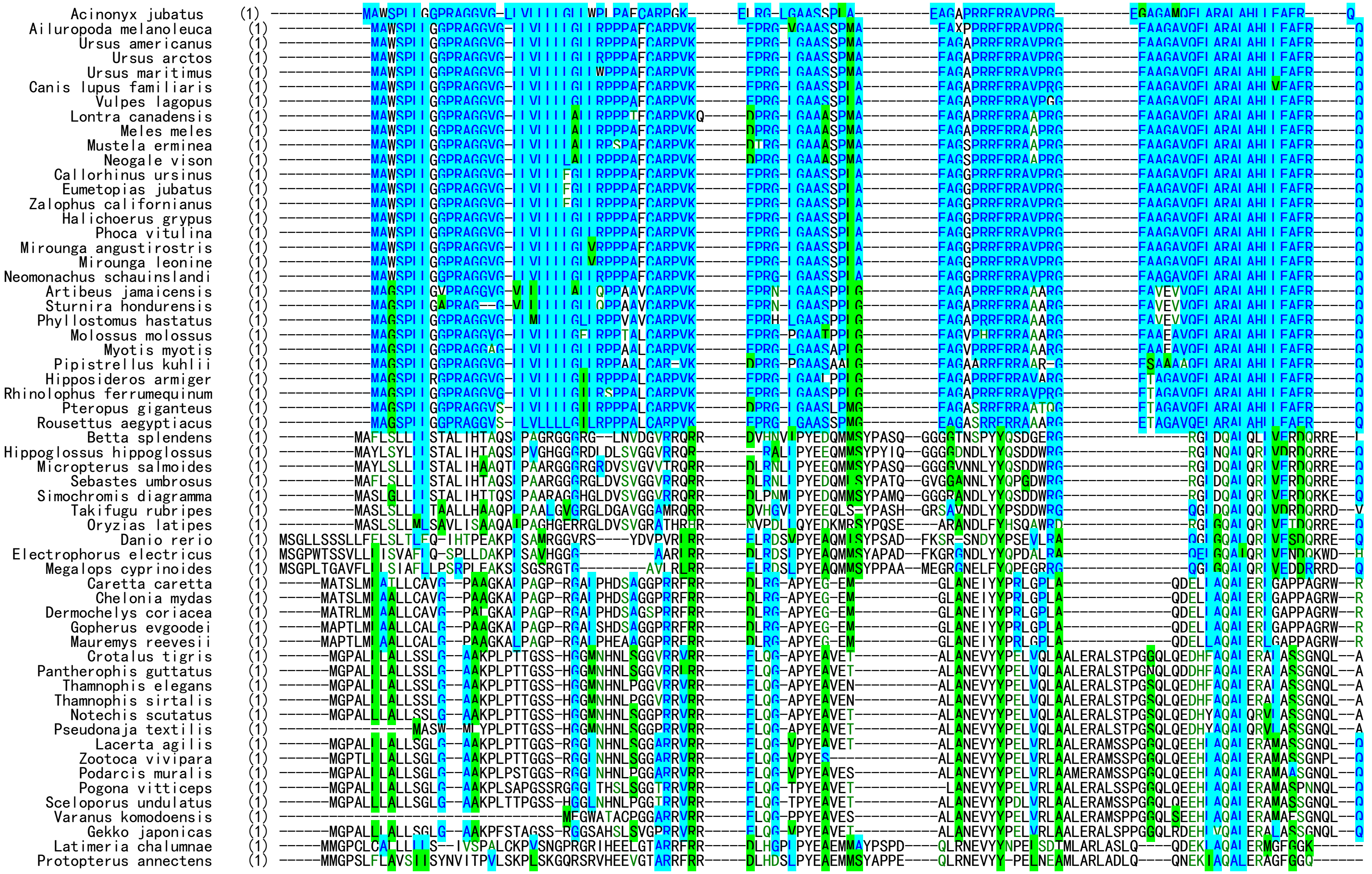

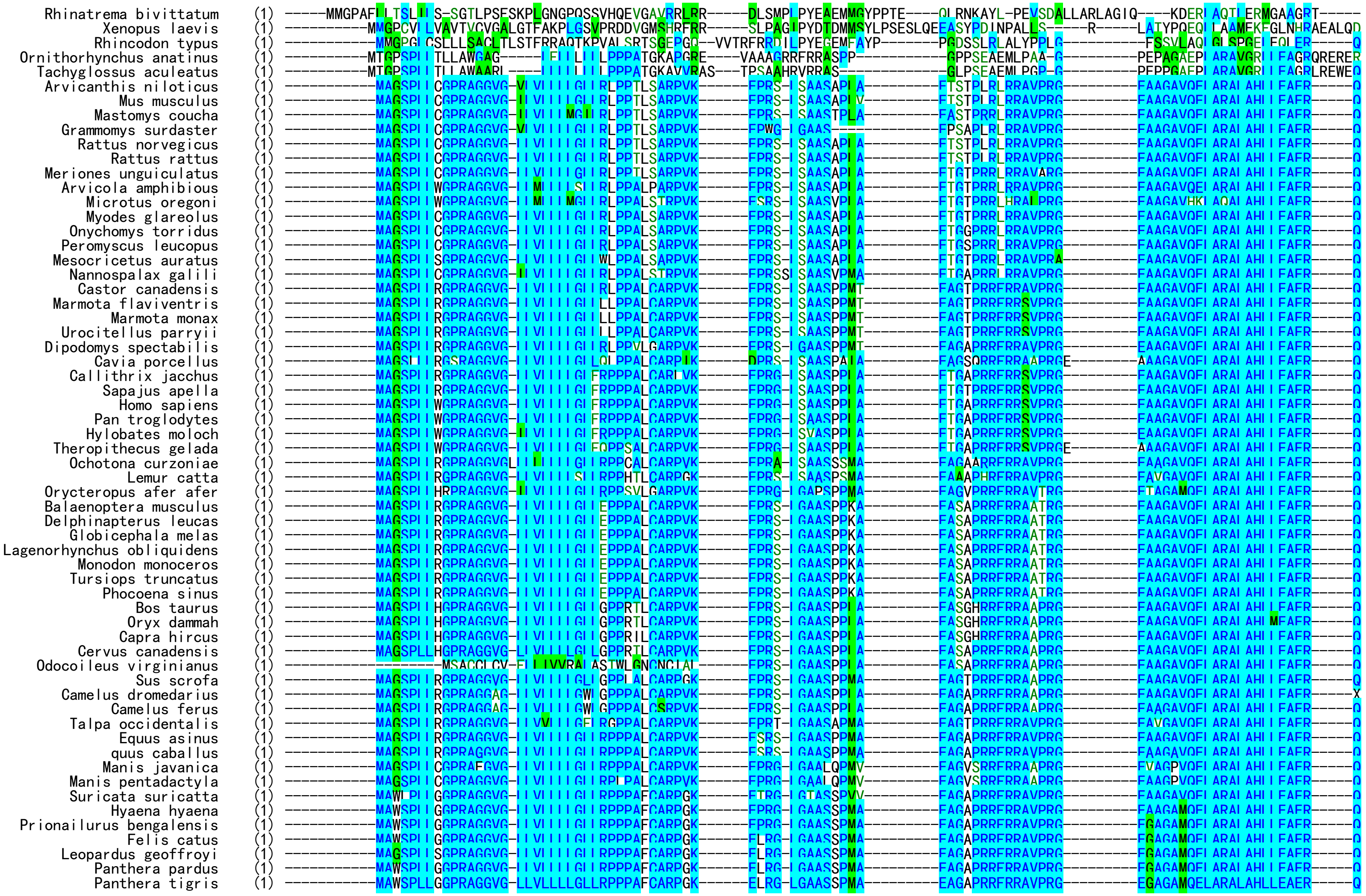
Rhinatrema bivittatum
Xenopus laevis
Rhincodon typus
Ornithorhynchus anatinus
Tachyglossus aculeatus
Arvicanthis niloticus
Mus musculus
Mastomys coucha
Grammomys surdaster
Rattus norvegicus
Rattus rattus
Meriones unguiculatus
Arvicola amphibious
Microtus oregoni
Myodes glareolus
Onychomys torridus
Peromyscus leucopus
Mesocricetus auratus
Nannospalax galili
Castor canadensis
Marmota flaviventris
Marmota monax
Urocitellus parryii
Dipodomys spectabilis
Cavia porcellus
Callithrix jacchus
Sapajus apella
Homo sapiens
Pan troglodytes
Hylobates moloch
Theropithecus gelada
Ochotona curzoniae
Lemur catta
Orycteropus afer afer
Balaenoptera musculus
Delphinapterus leucas
Globicephala melas
Lagenorhynchus obliquidens
Monodon monoceros
Tursiops truncatus
Phocoena sinus
Bos taurus
Oryx dammah
Capra hircus
Cervus canadensis
Odocoileus virginianus
Sus scrofa
Camelus dromedarius
Camelus ferus
Talpa occidentalis
Equus asinus
quus caballus
Manis javanica
Manis pentadactyla
Suricata suricatta
Hyaena hyaena
Prionailurus bengalensis
Felis catus
Leopardus geoffroyi
Panthera pardus
Panthera tigris

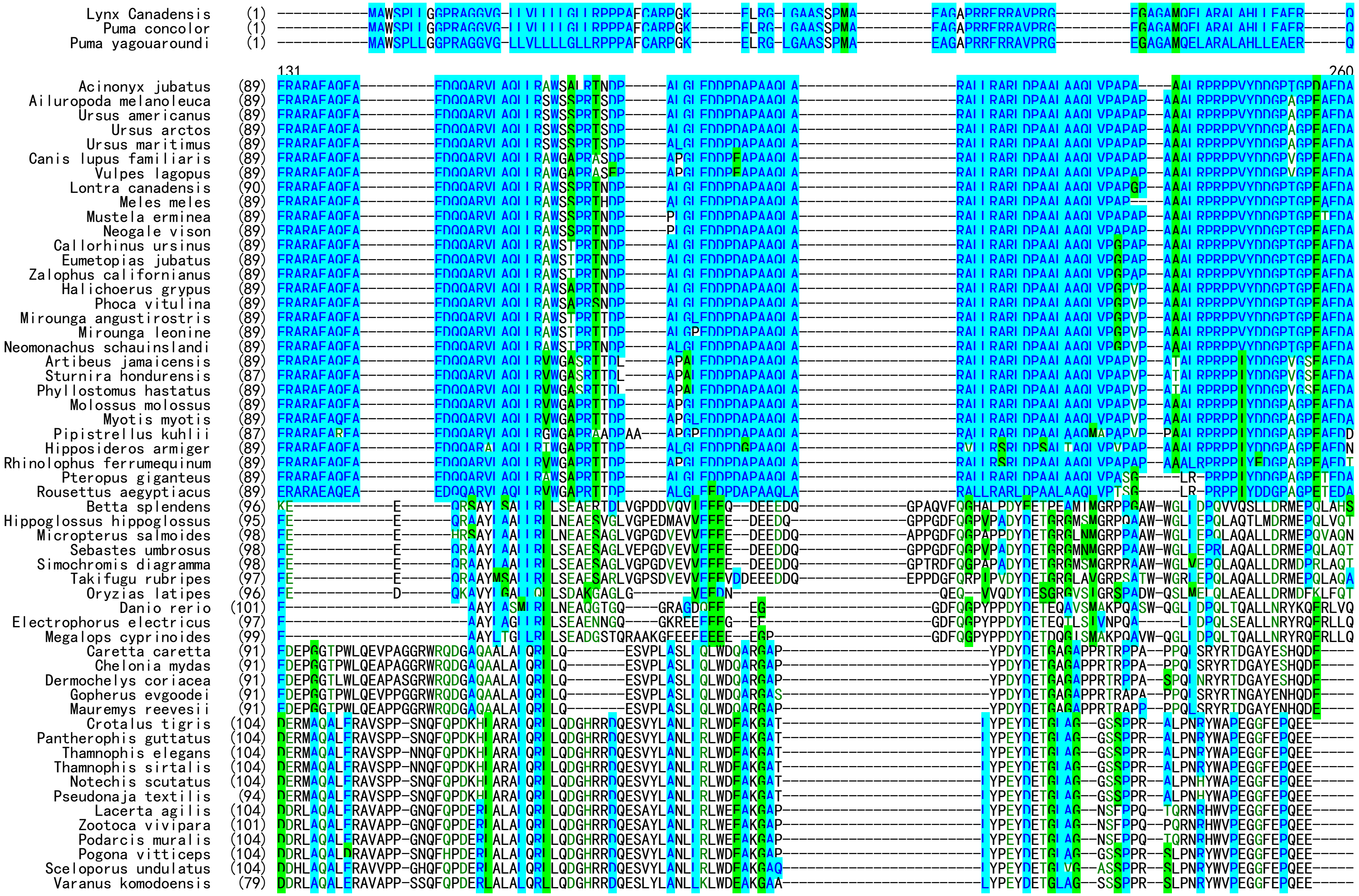

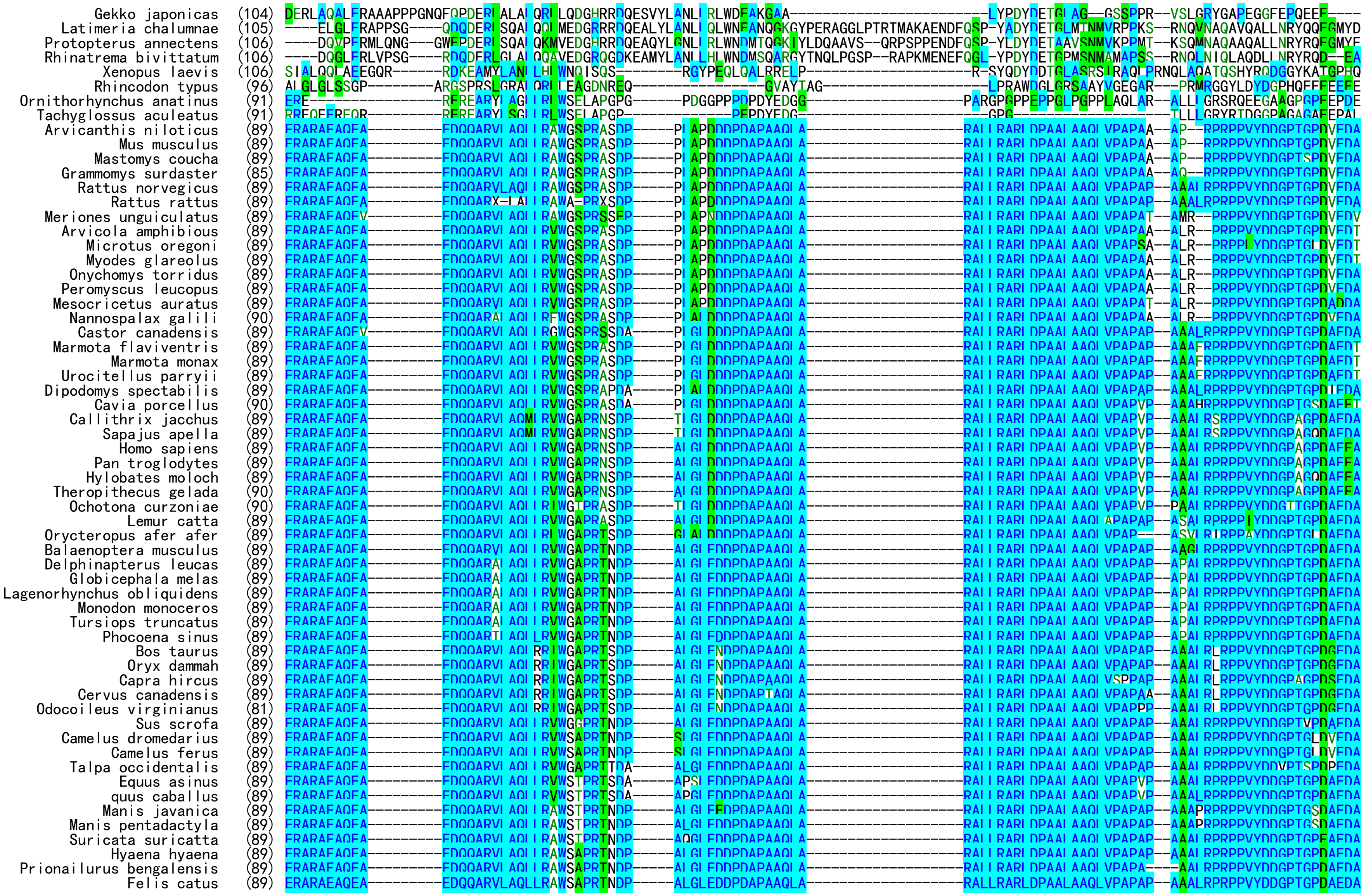

Gekko japonicas (104)
Latimeria chalumnae (105)
Protopterus annectens (106)
Rhinatrema bivittatum (106)
Xenopus laevis (106)
Rhincodon typus (96)
Ornithorhynchus anatinus (91)
Tachyglossus aculeatus (91)
Arvicanthis niloticus (89)
Mus musculus (89)
Mastomys coucha (89)
Grammomys surdaster (85)
Rattus norvegicus (89)
Rattus rattus (89)
Meriones unguiculatus (89)
Arvicola amphibious (89)
Microtus oregoni (89)
Myodes glareolus (89)
Onychomys torridus (89)
Peromyscus leucopus (89)
Mesocricetus auratus (89)
Nannospalax galili (90)
Castor canadensis (89)
Marmota flaviventris (89)
Marmota monax (89)
Urocitellus parryii (89)
Dipodomys spectabilis (89)
Cavia porcellus (90)
Callithrix jacchus (89)
Sapajus apella (89)
Homo sapiens (89)
Pan troglodytes (89)
Hylobates moloch (89)
Theropithecus gelada (90)
Ochotona curzoniae (90)
Lemur catta (89)
Orycteropus afer afer (89)
Balaenoptera musculus (89)
Delphinapterus leucas (89)
Globicephala melas (89)
Lagenorhynchus obliquidens (89)
Monodon monoceros (89)
Tursiops truncatus (89)
Phocoena sinus (89)
Bos taurus (89)
Oryx dammah (89)
Capra hircus (89)
Cervus canadensis (89)
Odocoileus virginianus (81)
Sus scrofa (89)
Camelus dromedarius (89)
Camelus ferus (89)
Talpa occidentalis (89)
Equus asinus (89)
quus caballus (89)
Manis javanica (89)
Manis pentadactyla (89)
Suricata suricatta (89)
Hyaena hyaena (89)
Prionailurus bengalensis (89)
Felis catus (89)

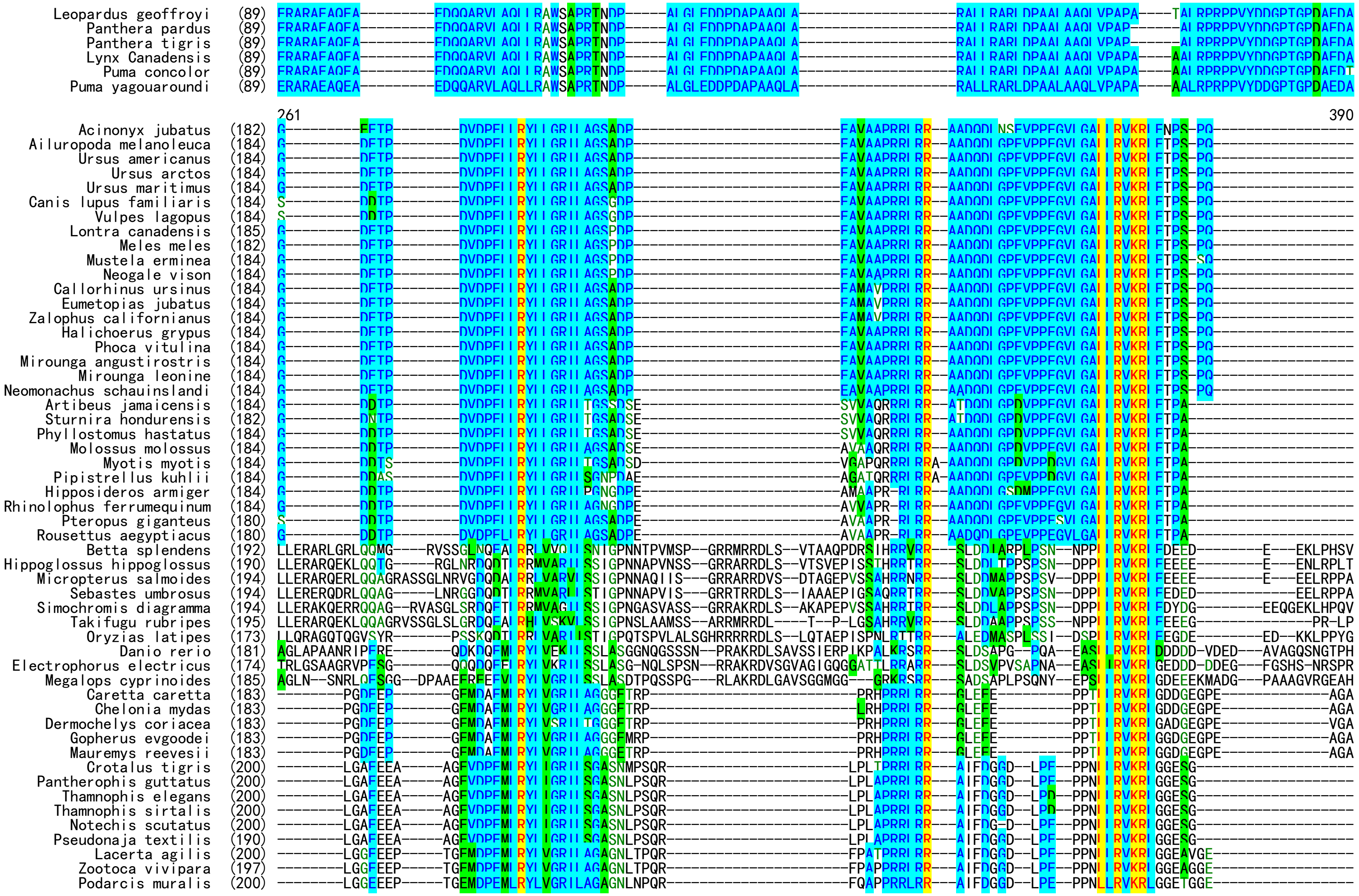
Leopardus geoffroyi (89)
Panthera pardus (89)
Panthera tigris (89)
Lynx Canadensis (89)
Puma concolor (89)
Puma yagouaroundi (89)
261
390
Acinonyx jubatus (182)
Ailuropoda melanoleuca (184)
Ursus americanus (184)
Ursus arctos (184)
Ursus maritimus (184)
Canis lupus familiaris (184)
Vulpes lagopus (184)
Lontra canadensis (185)
Meles meles (182)
Mustela erminea (184)
Neogale vison (184)
Callorhinus ursinus (184)
Eumetopias jubatus (184)
Zalophus californianus (184)
Halichoerus grypus (184)
Phoca vitulina (184)
Mirounga angustirostris (184)
Mirounga leonine (184)
Neomonachus schauinslandi (184)
Artibeus jamaicensis (184)
Sturnira hondurensis (182)
Phyllostomus hastatus (184)
Molossus molossus (184)
Myotis myotis (184)
Pipistrellus kuhlii (184)
Hipposideros armiger (184)
Rhinolophus ferrumequinum (184)
Pteropus giganteus (180)
Rousettus aegyptiacus (180)
Betta splendens (192)
Hippoglossus hippoglossus (190)
Micropterus salmoides (194)
Sebastes umbrosus (194)
Simochromis diagramma (194)
Takifugu rubripes (195)
Oryzias latipes (173)
Danio rerio (181)
Electrophorus electricus (174)
Megalops cyprinoides (185)
Caretta caretta (183)
Chelonia mydas (183)
Dermochelys coriacea (183)
Gopherus evgoodei (183)
Mauremys reevesii (183)
Crotalus tigris (200)
Pantherophis guttatus (200)
Thamnophis elegans (200)
Thamnophis sirtalis (200)
Notechis scutatus (200)
Pseudonaja textilis (190)
Lacerta agilis (200)
Zootoca vivipara (197)
Podarcis muralis (200)

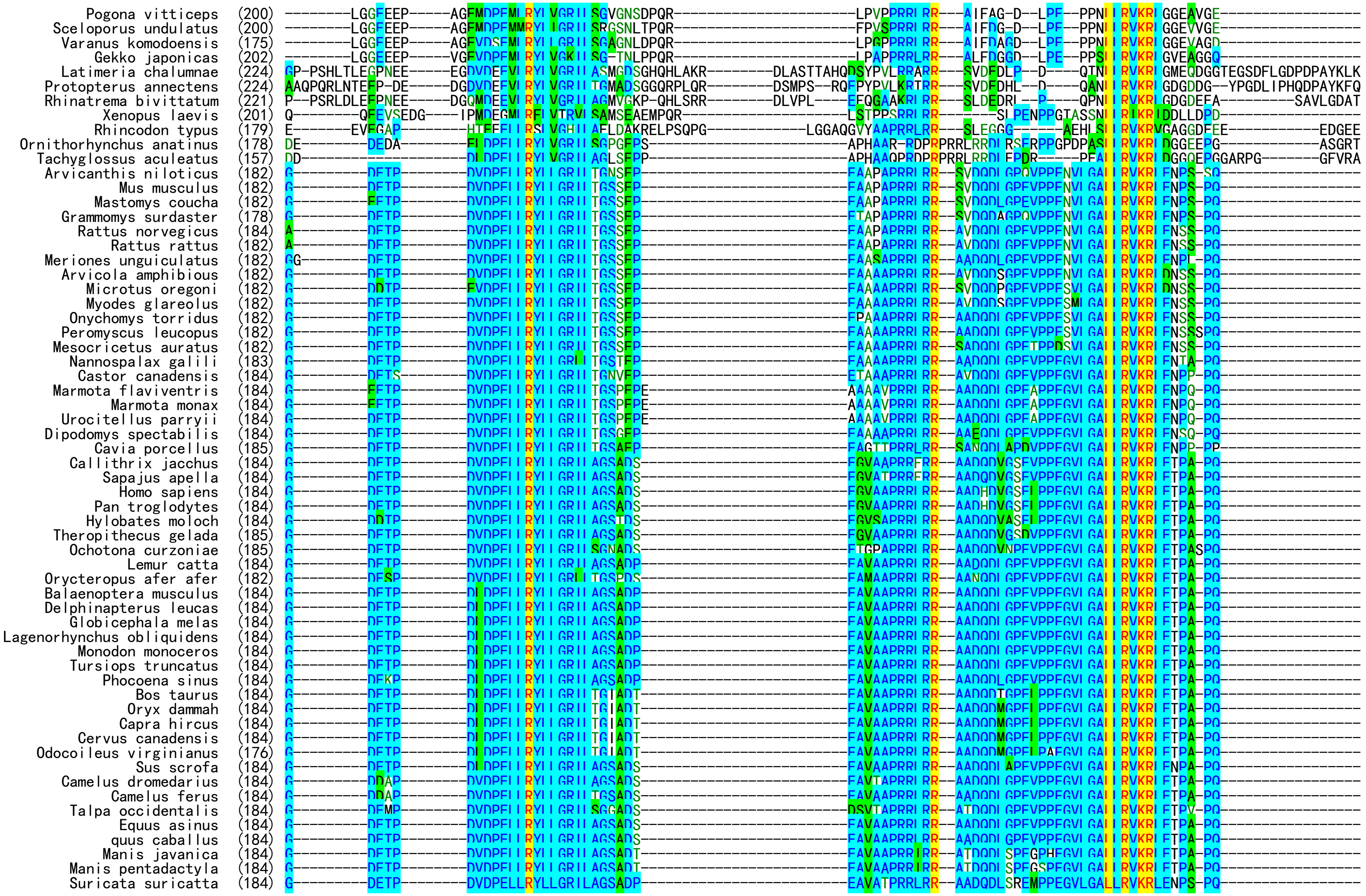
Pogona vitticeps (200)
Sceloporus undulatus (200)
Varanus komodoensis (175)
Gekko japonicas (202)
Latimeria chalumnae (224)
Protopterus annectens (224)
Rhinatrema bivittatum (221)
Xenopus laevis (201)
Rhincodon typus (179)
Ornithorhynchus anatinus (178)
Tachyglossus aculeatus (157)
Arvicanthis niloticus (182)
Mus musculus (182)
Mastomys coucha (182)
Grammomys surdaster (178)
Rattus norvegicus (184)
Rattus rattus (182)
Meriones unguiculatus (182)
Arvicola amphibious (182)
Microtus oregoni (182)
Myodes glareolus (182)
Onychomys torridus (182)
Peromyscus leucopus (182)
Mesocricetus auratus (182)
Nannospalax galili (183)
Castor canadensis (184)
Marmota flaviventris (184)
Marmota monax (184)
Urocitellus parryii (184)
Dipodomys spectabilis (184)
Cavia porcellus (185)
Callithrix jacchus (184)
Sapajus apella (184)
Homo sapiens (184)
Pan troglodytes (184)
Hylobates moloch (184)
Theropithecus gelada (185)
Ochotona curzoniae (185)
Lemur catta (184)
Orycteropus afer afer (182)
Balaenoptera musculus (184)
Delphinapterus leucas (184)
Globicephala melas (184)
Lagenorhynchus obliquidens (184)
Monodon monoceros (184)
Tursiops truncatus (184)
Phocoena sinus (184)
Bos taurus (184)
Oryx dammah (184)
Capra hircus (184)
Cervus canadensis (184)
Odocoileus virginianus (176)
Sus scrofa (184)
Camelus dromedarius (184)
Camelus ferus (184)
Talpa occidentalis (184)
Equus asinus (184)
quus caballus (184)
Manis javanica (184)
Manis pentadactyla (184)
Suricata suricatta (184)

```
          Hyaena hyaena  (184) G---------DETP--------DVDPELLRYLLGRILAGSADP-------------------------EAVAAPRRLRR--AADQDLSREVPPEGVLGALLRVKRLENPS-PQ-----------------
Prionailurus bengalensis (182) G---------EETP--------DVDPELLRYLLGRILAGSVDP-------------------------EAVAAPRRLRR--AADQDLSPEVPPEGVLGALLRVKRLENPS-PQ-----------------
             Felis catus (184) G---------EETP--------DVDPELLRYLLGRILAGSADS-------------------------EAVAAPRRLRR--AADQDLSPEVPPEGVLGALLRVKRLENPS-PQ-----------------
    Leopardus geoffroyi  (182) G---------EETP--------DVDPELLRYLLGRILAGSADP-------------------------EAVAAPRRLRR--AADQDLSPEVPPEGVLGALLRVKRLENPS-PQ-----------------
         Panthera pardus (180) G---------EETP--------DVDPELLRYLLGRILAGSADP-------------------------EAVAAPRRLRR--AADQDLSPEVPPEGVLGALLRVKRLENPS-PQ-----------------
         Panthera tigris (180) G---------EETP--------DVDPELLRYLLGRILAGSADP-------------------------EAVAAPRRLRR--AADQDLSPEVPPEGVLGALLRVKRLENPS-PQ-----------------
         Lynx Canadensis (182) G---------EETP--------DVDPELLRYLLGRILAGSADP-------------------------EAVAAPRRLRR--AADQDLSPEVPPEGVLGALLRVKRLENPS-PQ-----------------
           Puma concolor (182) G---------EETP--------DVDPELLRYLLGRILAGSADP-------------------------EAVAAPRRLRR--AADQDLSPEVPPEGVLGALLRVKRLENPS-PQ-----------------
       Puma yagouaroundi (182) G---------EETP--------DVDPELLRYLLGRILAGSADP-------------------------EAVAAPRRLRR--AADQDLSPEVPPEGVLGALLRVKRLENPS-PQ-----------------

                               391                                                         453
         Acinonyx jubatus (250) APVRRLLPP------------------------------------------------------
   Ailuropoda melanoleuca (252) APMRRLLPP------------------------------------------------------
        Ursus americanus (252) APVRRLLPP------------------------------------------------------
            Ursus arctos (252) APVRRLLPP------------------------------------------------------
         Ursus maritimus (252) APVRRLLPP------------------------------------------------------
     Canis lupus familiaris (252) APVRRLLPP------------------------------------------------------
          Vulpes lagopus (252) AQVRRLLPP------------------------------------------------------
       Lontra canadensis (253) APVRRLLPP------------------------------------------------------
             Meles meles (250) APVRRLLPP------------------------------------------------------
         Mustela erminea (252) APVRRLLPP------------------------------------------------------
           Neogale vison (252) APVRRLLPP------------------------------------------------------
      Callorhinus ursinus (252) APVRRLLPP------------------------------------------------------
       Eumetopias jubatus (252) APVRRLLPP------------------------------------------------------
     Zalophus californianus (252) APVRRLLPP------------------------------------------------------
       Halichoerus grypus (252) APVRRLLPP------------------------------------------------------
           Phoca vitulina (252) APVRRLLPP------------------------------------------------------
   Mirounga angustirostris (252) APVRRLLPP------------------------------------------------------
         Mirounga leonine (252) APVRRLLPP------------------------------------------------------
   Neomonachus schauinslandi (252) APVCRLLPH------------------------------------------------------
       Artibeus jamaicensis (251) PPARRLLPP------------------------------------------------------
      Sturnira hondurensis (249) PPARRLLPP------------------------------------------------------
      Phyllostomus hastatus (251) PPARRLLPP------------------------------------------------------
        Molossus molossus (251) PPAQRRLVPA-----------------------------------------------------
            Myotis myotis (252) PQARRLLPP------------------------------------------------------
     Pipistrellus kuhlii (252) PLARRLLPP------------------------------------------------------
     Hipposideros armiger (250) PPARRLLPP------------------------------------------------------
  Rhinolophus ferrumequinum (250) PPARRLLPP------------------------------------------------------
       Pteropus giganteus (246) PPARRLLPP------------------------------------------------------
     Rousettus aegyptiacus (246) PPARRLLPP------------------------------------------------------
          Betta splendens (298) TGLQRMKRIDTMATTSMEKLNIGNHWRKRR-ALNYDHHILIDQILDYMRE-------------
   Hippoglossus hippoglossus (295) AGLQRMKRIDTMATAAEEELNHGIRRRRRRAAMNYDPQILIDQIVNYMRE-------------
     Micropterus salmoides (303) AGLQRMKRIDTMATAAMEELNNGGRRRRRRAVLNYDPQMLIDQIVEYMRE-------------
        Sebastes umbrosus (297) TGLQRMKRIDTMATAGVEELNHGSRRRRRREALNYDPQILIDQILEYM---------------
     Simochromis diagramma (305) VGLQRMKRIDTTSTGDAEELNHGSRRRRRRAALNYDPQILVDQILEYMRE-------------
        Takifugu rubripes (296) AGLQRMKRID-------EDLGQGSRRRRRRAVLTFDPQLLVRQVMEYMRE-------------
          Oryzias latipes (279) ATLQRIKRIDTLAAS--EETNHVSR-RHRRAAVNYNSKLQMDQILEYMRG-------------
              Danio rerio (291) TGLQRMKRIDTDLPPP-------KKHSRKRRALSYDPALIAQHILQYLPA-------------
    Electrophorus electricus (284) VGLQRMKRIDTDLTPP-------QKHSRKRRAVTYDPNVIAQHILQYLPA-------------
      Megalops cyprinoides (297) VGLQRMKRIDSDLQQ--------QKHKRRRRALTYDPNLLAQRILQYLPQ-------------
           Caretta caretta (248) PQLQRAKRTEEEEE----------------AGGGRRGVYGGEPLLRYLPE-------------
            Chelonia mydas (248) PQLQRAKRTEEEEE----------------AGGGRRGVYGGEPLLRYLPE-------------
     Dermochelys coriacea (248) PQLQRAKRTEEEEE----------------AGGGRRGAYGGEPLLRYLPE-------------
         Gopherus evgoodei (248) PQLQRAKRTEEEEEE--------------AAVGGRRGAYGGEPLLRYLPE-------------
        Mauremys reevesii (248) PQLQRAKRTEEEEEE--------------AAVGGRRGAYGGEPLLRYLPE-------------
           Crotalus tigris (269) DELQRVKRTEGEG------------------VGGRRSMSGGTQRLRYLSE-------------
     Pantherophis guttatus (269) DELQRVKRTEGEG------------------VGGRRSMSGGTQRLRYLSE-------------
        Thamnophis elegans (269) DELQRVKRTEGEG------------------VGGRRSMSGGTQRLRYLSE-------------
       Thamnophis sirtalis (269) DELQRVKRTEGEG------------------VGGRRSMSGGTQRLRYLSE-------------
        Notechis scutatus (268) DELQRVKQTEGEG------------------VGGRRSMSGGTQRVRYLSE-------------
      Pseudonaja textilis (259) DELQRVKQTEGEG------------------VGGRRSMSGGTQRLRYLSE-------------
```

```
            Lacerta agilis (271) PQLQRVKRTEGEG------------------AGGRRGVSGGVQRLRYLPE-------------
           Zootoca vivipara (268) PQLQRVKRTEGEG------------------AGGRRGVSGGMQRLRYLPE-------------
           Podarcis muralis (271) PQLQRVKRTEGEG------------------AGGRRGVSGGVQRLRYLPE-------------
          Pogona vitticeps (270) TQLQRVKRTEGEG------------------AGGRRGATGGTQRLRYVSE-------------
      Sceloporus undulatus (270) SQLQRVKRTEGEG------------------VGGRRGVAGGSQRLQYIPE-------------
        Varanus komodoensis (246) PQLQRVKRTEGEG------------------AGGKRGGTGGSQRLRYLPE-------------
           Gekko japonicas (272) PQLQRIKRTDGGGRK--------------KDGGSSKADAKSKRHVEYLKSFKAISDIAAVADK
      Latimeria chalumnae (331) GGLQRAKRIDEQLDGL-------AAPIRRKRHAGYDEAEFAERVLKYLPD-------------
     Protopterus annectens (327) GGLQRMKRIDDQLEEP-------SKSIRSRRYAAYDQGSLAERALKYLPE-------------
     Rhinatrema bivittatum (313) YRLQRAKRAEEPVDEP-------ATALRSKRYAAYSDPEFPEHFLKYLPE-------------
            Xenopus laevis (275) YVANGLLRRK-----------------RIDGDLEARPGHYTDQLLKYLPD-------------
          Rhincodon typus (262) EGGARYPGPDRRRAKG-------FGVRPRLARAPLPSGQLARRLARLLPD-------------
  Ornithorhynchus anatinus (255) RPKRGPLHPGAGLAQQVQRYRPH----------------------------------------
     Tachyglossus aculeatus (231) RPARG------------LPYRPH----------------------------------------
      Arvicanthis niloticus (250) APARRIIPP------------------------------------------------------
              Mus musculus (250) APARRIIPP------------------------------------------------------
          Mastomys coucha (250) APARRIIPP------------------------------------------------------
        Grammomys surdaster (246) APARRIIPP------------------------------------------------------
         Rattus norvegicus (252) APARRIIPP------------------------------------------------------
              Rattus rattus (250) APARRIIPP------------------------------------------------------
     Meriones unguiculatus (251) APARRIIPP------------------------------------------------------
       Arvicola amphibious (250) GPARRIIPP------------------------------------------------------
          Microtus oregoni (250) APARRIIPP------------------------------------------------------
          Myodes glareolus (250) APARRIIPP------------------------------------------------------
        Onychomys torridus (250) APARRIIPP------------------------------------------------------
        Peromyscus leucopus (251) APARRIIPP------------------------------------------------------
       Mesocricetus auratus (250) APARRIIPP------------------------------------------------------
        Nannospalax galili (251) APARRIIPA------------------------------------------------------
         Castor canadensis (252) APARRIIPP------------------------------------------------------
      Marmota flaviventris (253) APARRIIPP------------------------------------------------------
             Marmota monax (253) APARRIIPP------------------------------------------------------
       Urocitellus parryii (253) APARRIIPP------------------------------------------------------
      Dipodomys spectabilis (252) APARRIIPP------------------------------------------------------
          Cavia porcellus (253) APARRIIPP------------------------------------------------------
        Callithrix jacchus (252) MPARRIIPP------------------------------------------------------
            Sapajus apella (252) MPARRIIPP------------------------------------------------------
              Homo sapiens (252) VPARRIIPP------------------------------------------------------
          Pan troglodytes (252) VPARRIIPP------------------------------------------------------
           Hylobates moloch (252) VPARRIIPP------------------------------------------------------
       Theropithecus gelada (253) VPARRIIPP------------------------------------------------------
        Ochotona curzoniae (254) APARRIIPP------------------------------------------------------
              Lemur catta (252) APARRIIPP------------------------------------------------------
       Orycteropus afer afer (250) APVRRIIPP------------------------------------------------------
      Balaenoptera musculus (252) APARRIIP-------------------------------------------------------
     Delphinapterus leucas (252) APARRIIP-------------------------------------------------------
        Globicephala melas (252) APARRIIP-------------------------------------------------------
 Lagenorhynchus obliquidens (252) APARRIIP-------------------------------------------------------
        Monodon monoceros (252) APARRIIP-------------------------------------------------------
        Tursiops truncatus (252) APARRIIP-------------------------------------------------------
            Phocoena sinus (252) APARRIIP-------------------------------------------------------
                Bos taurus (252) APARRIIP-------------------------------------------------------
              Oryx dammah (252) APARRIIP-------------------------------------------------------
              Capra hircus (252) APARRIIP-------------------------------------------------------
         Cervus canadensis (252) APARRIIP-------------------------------------------------------
     Odocoileus virginianus (244) APARRIIP-------------------------------------------------------
                Sus scrofa (252) APARRIIP-------------------------------------------------------
        Camelus dromedarius (252) APARRIIP-------------------------------------------------------
             Camelus ferus (252) APARRIIP-------------------------------------------------------
       Talpa occidentalis (252) APGRRIIPP------------------------------------------------------
              Equus asinus (252) APARRIIPP------------------------------------------------------
             quus caballus (252) APARRILPP------------------------------------------------------
```

```
        Manis javanica  (252) APGRRIIPPLAD---------------------------------------------------
    Manis pentadactyla  (252) APGRRIIPPLAA---------------------------------------------------
    Suricata suricatta  (252) APVRRIIPP------------------------------------------------------
        Hyaena hyaena   (252) APVRRIIPA------------------------------------------------------
Prionailurus bengalensis (250) APVRRIIPP------------------------------------------------------
           Felis catus  (252) APVRRIIPP------------------------------------------------------
   Leopardus geoffroyi  (250) APVRRIIPP------------------------------------------------------
      Panthera pardus   (248) APVRRIIPP------------------------------------------------------
      Panthera tigris   (248) APVRRIIPP------------------------------------------------------
      Lynx Canadensis   (250) APVRRIIPP------------------------------------------------------
        Puma concolor   (250) APVRRIIPP------------------------------------------------------
     Puma yagouaroundi  (250) APVRRLLPP------------------------------------------------------
```

**Fig. S6.** Amino acid sequence alignment of PCSK1N/proSAAS orthologs from different species.
